# Inducible mechanisms of disease tolerance provide an alternative strategy of acquired immunity to malaria

**DOI:** 10.1101/2020.10.01.322180

**Authors:** Wiebke Nahrendorf, Alasdair Ivens, Philip J. Spence

## Abstract

Immunity to malaria is often considered slow to develop but this only applies to defense mechanisms that function to eliminate parasites (resistance). In contrast, immunity to severe disease can be acquired quickly and without the need for improved pathogen control (tolerance). We show that a single malaria episode is sufficient to induce host adaptations that can minimise inflammation, prevent tissue damage and avert endothelium activation, a hallmark of severe disease. Furthermore, monocytes are functionally reprogrammed in tolerised hosts to prevent their differentiation into inflammatory macrophages and instead promote mechanisms of stress tolerance to protect their niche. This alternative fate is not underpinned by epigenetic reprogramming of bone marrow progenitors but is imprinted within the remodelled spleen. Crucially, all of these adaptations operate independently of pathogen load and limit the damage caused by malaria parasites in subsequent infections. Inducible mechanisms of disease tolerance therefore provide an alternative strategy of acquired immunity.

## Introduction

Mechanisms of host resistance can eliminate pathogens, but it is disease tolerance that functions to preserve life. Tolerance mechanisms of host defense do not have a direct impact on pathogen load and instead act to minimise tissue damage caused by the pathogen - and the immune response targeting it. They also function to protect vital homeostatic processes, such as energy metabolism, under conditions of infection-induced stress (Martins et al., 2019). We have taken tremendous steps to understand acquired resistance mechanisms that can eliminate pathogens and provide sterile immunity. On the other hand, it is unclear whether mechanisms of disease tolerance can persist after pathogen clearance to provide an alternative strategy of acquired immunity. And it is these mechanisms that are likely to be at the forefront of host defense when sterile immunity can not be generated.

We propose that immunity to severe life-threatening malaria is underpinned by acquired mechanisms of disease tolerance. The majority of malaria-induced deaths occur in children under the age of 5 infected with *Plasmodium falciparum* (Weiss et al., 2019). A landmark prospective study in Tanzania followed 882 children from birth and showed that the risk of developing severe malaria is highest in the first few infections of life, and very few children (< 1.8 %) have more than one severe episode (Goncalves et al., 2014). These data therefore support the longstanding view that immunity against severe malaria is acquired rapidly - often before 12-months of age (Gupta et al., 1999; Marsh and Snow, 1999). Crucially, this study further showed that children who survive severe malaria are frequently reinfected and experience episodes of febrile malaria with similar or even higher parasite densities (Goncalves et al., 2014). Immunity to severe forms of malaria is therefore not due to improved parasite elimination (resistance) but instead underpinned by the improved ability of the host to limit the pathological consequences of infection (tolerance).

In order to survive a first malaria episode a large body of evidence supports that disease tolerance is critical: since acute malaria causes hypoglycaemia, the ability to maintain blood glucose levels within dynamic range – possibly in crosstalk with iron metabolism (Weis et al., 2017) – can determine whether the host lives or dies (Cumnock et al., 2018). Furthermore, the induction of hemeoxygenase 1 (HMOX1) by nitric oxide (Jeney et al., 2014) leads to the detoxification of free heme to bilirubin (Seixas et al., 2009), thus protecting renal proximal tubule epithelial cells, which in turn prevents acute kidney injury (Ramos et al., 2019). Nonetheless, while these metabolic adaptations are doubtless essential for the survival of naive hosts there is no evidence as yet that they can be acquired (or learned) and then applied more effectively in subsequent infections to provide clinical immunity.

Instead, the most effective way to induce disease tolerance may be through host control of inflammation (Medzhitov et al., 2012), which can limit collateral tissue damage and avert fatal metabolic perturbations. In malaria, the ability to control systemic inflammation may also minimise the detrimental effects of parasite sequestration by reducing activation of the endothelium and restricting available binding sites in the microvasculature (Schofield and Grau, 2005). Field studies support the idea that dampening the systemic inflammatory response to *P. falciparum* can save lives; lower levels of pro-inflammatory plasma cytokines are found in children in Malawi who survive severe malaria compared to those who subsequently died (Mandala et al., 2017). Furthermore, inflammation can be reduced in the absence of improved parasite clearance; Ghanaian children in a high transmission area have higher parasite densities but less systemic inflammation and far fewer febrile episodes compared to children in a lower transmission setting (Ademolue et al., 2017).

The blood cycle in malaria unleashes a plethora of parasite-derived as well as host tissue damage-associated signals (e.g. free heme from ruptured red cells), which are sensed by innate immune cells. Additionally, pronounced changes in host physiology (such as hypoxia and acidaemia) are hallmarks of severe disease (von Seidlein et al., 2012). Together all of these diverse signals trigger monocytes and macrophages to produce pro-inflammatory molecules, many of which have been associated with a poor prognosis including the prototypical myeloid-derived cytokines TNF and IL-6 (Mandala et al., 2017). Monocytes and macrophages can also directly cause severe malarial anaemia through extensive bystander phagocytosis of healthy uninfected red cells in the spleen and impairment of erythropoiesis in the bone marrow (Jakeman et al., 1999; Pathak and Ghosh, 2016). Given these key roles in malaria pathogenesis, the ability to rapidly alter the monocyte and macrophage response after a single malaria episode has the potential to not only minimise inflammation-induced tissue damage but additionally avoid life-threatening complications. Importantly, an emerging body of literature shows that the response of myeloid cells to infection is not hardwired, as previously thought, but can in fact be reprogrammed by pathogens and their products. This was first demonstrated by stimulating bone marrow-derived macrophages with bacterial lipopolysaccharide (LPS) *in vitro*, which reduces the production of pro-inflammatory cytokines and increases the release of antimicrobial effector molecules upon re-stimulation (Foster et al., 2007). The ability of monocytes and macrophages to learn from a previous exposure and specialise their response to repeated pathogen encounters through cell intrinsic modifications is termed innate memory (Netea et al., 2016). Although myeloid cells are usually short-lived and terminally differentiated, memory can nevertheless be imprinted through the epigenetic reprogramming of either long-lived tissue-resident macrophages (Wendeln et al., 2018) or progenitor cells in the bone marrow (Kaufmann et al., 2018).

Innate memory is not antigen-specific and has been shown to operate independently of pathogen load (Dominguez-Andres and Netea, 2018; Seeley and Ghosh, 2017). Furthermore, monocytes obtained from Malian children produce less pro-inflammatory cytokines when stimulated *in vitro* if they were isolated after an episode of febrile malaria (as compared to before infection) (Portugal et al., 2014) suggesting that human monocytes can be intrinsically modified by malaria parasites. We therefore hypothesise that innate memory - leading to the functional specialisation of monocytes and macrophages to limit inflammation and associated pathology - offers the most compelling explanation for how immunity to severe malaria can be acquired so quickly and without the need for enhanced parasite control.

## Results

To investigate acquired mechanisms of disease tolerance and the role of innate memory *in vivo* we needed to examine monocyte progenitors in the bone marrow and long-lived tissue-resident macrophages in the spleen. And since these tissues are not readily accessible in human malaria we needed a model that would recapitulate at least some key features of human infection. Given that a meta-analysis of malariatherapy data shows naive human hosts quickly adapt to tolerate chronic parasitaemia (for example, by increasing their pyrogenic threshold (Gatton and Cheng, 2002)) we chose a rodent malaria parasite (*Plasmodium chabaudi*) that establishes chronic recrudescing infections in laboratory mice. Importantly, experimental infections were initiated with sporozoites, since we have previously shown that mosquito transmission resets expression of the large sub-telomeric multi-gene families that control parasite virulence (Spence et al., 2013). And furthermore, we used two parasite genotypes to try and uncouple the relative contribution of parasite-derived versus damage-associated signals in promoting mechanisms of tolerance. *P. chabaudi* AS causes a mild infection, characterised by a low pathogen load and few clinical symptoms (Figures 1A, 1B and 1C). In contrast, *P. chabaudi* AJ (which has more than 140,000 SNPs cf. *P. chabaudi* AS (Otto et al., 2014)) leads to acute hyperparasitaemia and severe anaemia (Figures 1B and 1C), accompanied by hypothermia and prostration (Figure S1A). AJ shares many key features with AS such as synchrony, chronicity and a persisting low-grade anaemia (Figures 1B, 1C, S1B and S1C), and yet whilst AS sequesters in key immune sites such as spleen and bone marrow (Brugat et al., 2014) we find no evidence that AJ sequesters in host tissues (Figure S1D).

**Figure 1.**
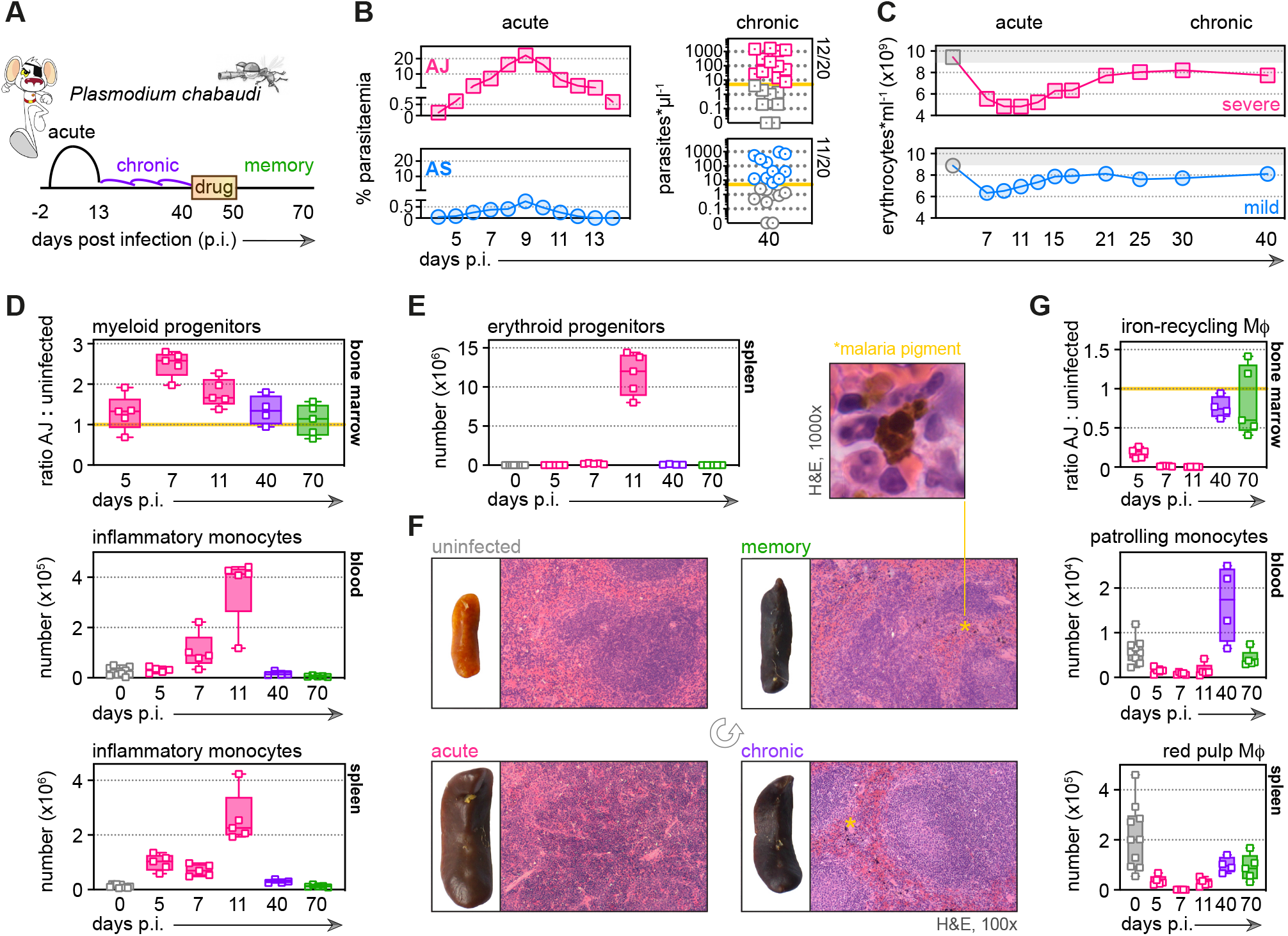
Malaria triggers emergency myelopoiesis and obliterates tissue-resident macrophages. (A) C57Bl/6 mice were infected with *Plasmodium chabaudi* AJ or AS sporozoites; the blood-stage of infection started 2-days later after the release of merozoites from the liver. Mice were chronically infected for 40-days, at which point we administered the antimalarial drug chloroquine. Memory responses were assessed 30-days thereafter. Note that we exclusively use days post infection (p.i.) to refer to the blood-stage of malaria. (B) Acute parasitaemia was monitored daily using Giemsa stained thin blood films and chronic infection was verified 40-days p.i. by qPCR (n = 20 per group). Symbols below the limit of detection (5 parasites*μl^−1^) are coloured grey and these mice were excluded from the study. (C) The mean number of erythrocytes*ml^−1^ is shown before (grey symbols) and during infection (n = 10 for AJ and n = 14 for AS). Severe anaemia is defined as > 50 % loss of red cells. (D and E) Inflammatory monocytes and progenitors from uninfected mice (0-days p.i.), AJ infected mice (5, 7, 11 and 40-days p.i.) and once-infected mice (memory, 70-days p.i.) were analysed by flow cytometry (n = 4-5 per time point, box-plots show median and IQR). Uninfected age-matched controls were analysed at each time point and pooled for graphing (n = 10); absolute counts are shown for blood and spleen. Myeloid progenitors in (D) are shown as a ratio of infected:uninfected at each time point because bone marrow cellularity increases with age. (F) Paraffin embedded spleen sections were H&E stained (11-days p.i. for acute AJ infection) - examples of malaria pigment in chronically infected and once-infected mice are marked with an asterisk. (G) Tissue-resident macrophages (MΦ) from uninfected mice, AJ infected mice and once-infected mice were analysed by flow cytometry (n = 4-5 per time point, box-plots show median and IQR). Absolute counts (blood and spleen) and cell ratios (bone marrow) are shown exactly as described for (D and E). *see also Figures S1 and S2*

### Malaria triggers emergency myelopoiesis and obliterates tissue-resident macrophages

To ask whether malaria can functionally reprogramme myeloid cells we must first understand their response to acute infection in a naive host; we started by mapping their dynamics in our severe model of disease. We found that the bone marrow quickly prioritises myelopoiesis by increasing the number of granulocyte monocyte progenitors (GMP) (Figure 1D). Consequently, a huge number of inflammatory monocytes and neutrophils are released into circulation and recruited into their target organ - the spleen (Figures 1D and S2A). Furthermore, megakaryocyte erythroid progenitors (MEP) appear *de novo* in the spleen (Figure 1E); this extramedullary mechanism of erythropoiesis likely represents a division of labour in an attempt to compensate for the loss of erythroid progenitors in the bone marrow (Pathak and Ghosh, 2016). We also observed major histological changes in tissue structure and integrity with reduced cellularity in the bone marrow contrasting starkly with marked splenomegaly, which was accompanied by a complete loss of organisation between red and white pulp (Figures 1F, S2B and S2C).

Strikingly, we found that long-lived prenatally seeded tissue-resident macrophages rapidly disappear during acute infection, including specialised iron-recycling macrophages (Figure 1G). Since red pulp macrophages are the only cells that can store and recycle iron in the spleen their disappearance thus means that ferric iron, which can be revealed histologically with Prussian Blue staining, is completely absent at the peak of infection (Figures S2D and S2E). We could further demonstrate that patrolling monocytes (Carlin et al., 2013), often regarded as the tissue-resident macrophages of the vasculature (Mildner et al., 2017), also disappear early in infection (Figure 1G). These findings therefore place inflammatory monocytes at the centre of the acute phase response, since they now provide the only route through which to phagocytose and clear infected red cells.

### Monocytes differentiate into inflammatory macrophages in naive hosts

We therefore carefully characterised the fate and function of inflammatory monocytes in the spleen by RNA-sequencing (Figure 2A) and used clueGO to reveal the complexity and diversity of their response to a first encounter with malaria parasites (Bindea et al., 2009; Mlecnik et al., 2014). ClueGO assigns significant gene ontology (GO) terms based on differential gene expression and groups them into functional networks by relatedness. When we merged all linked nodes into supergroups (see methods) we found that more than one third of all GO terms were related to regulation of defence (Figures 2B and 2C). Furthermore, clueGO identified interferon signaling as a potential regulator of monocyte fate (Figure 2B) and interferon-inducible guanylate binding proteins were highly upregulated (Figure S3A).

**Figure 2.**
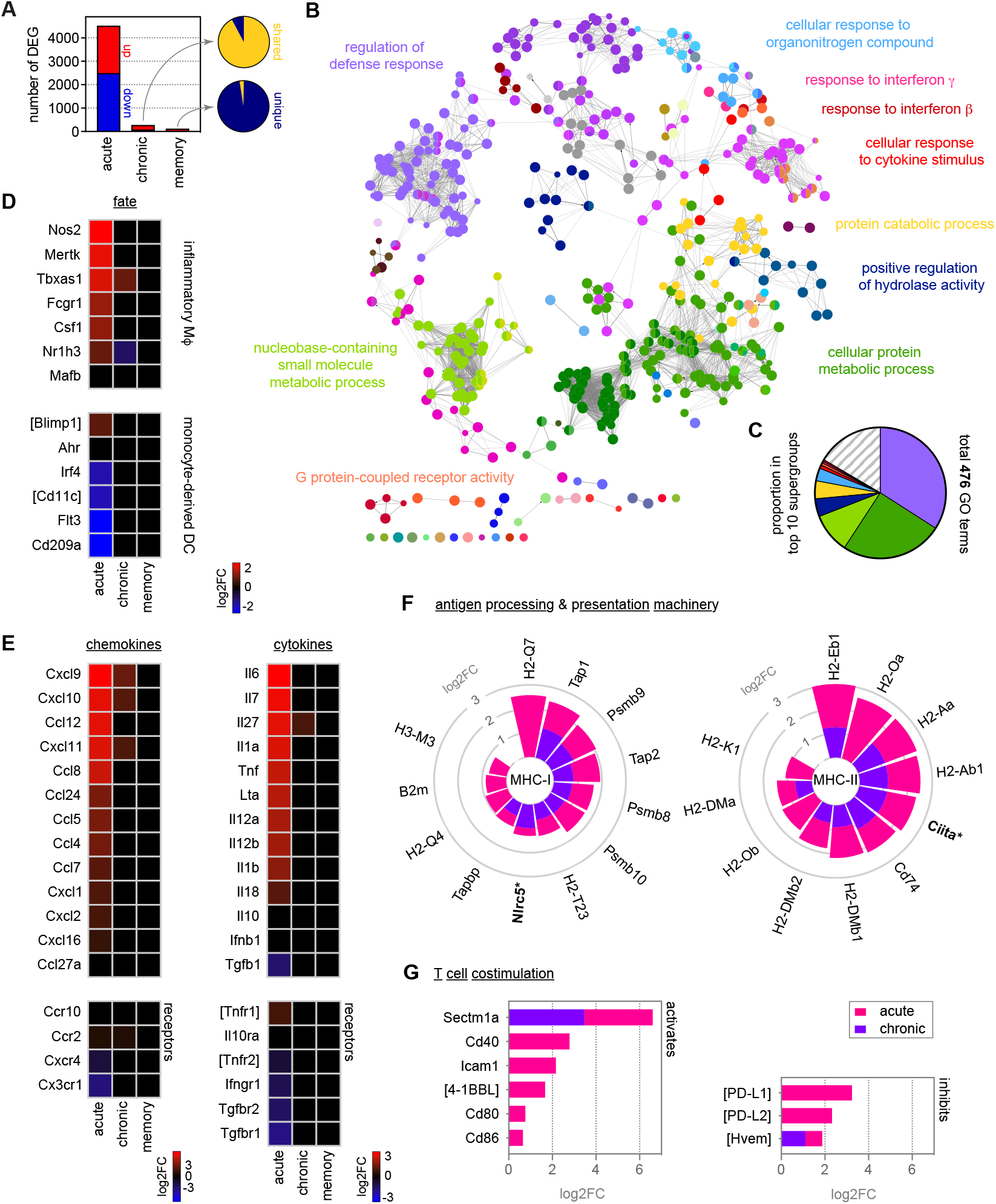
Monocytes differentiate into inflammatory macrophages in naive hosts but become quiescent in chronic infection. (A) RNA sequencing of spleen monocytes flow-sorted from AJ infected mice (7 and 40-days p.i. for acute and chronic, respectively) and once-infected mice (memory, 70-days p.i.). At each time point the number of differentially expressed genes (DEG, p_adj_ < 0.01, > 1.5 fold change) was assessed relative to uninfected controls. Pies show the proportion of shared or unique DEG between chronic & acute infection (top) or memory & chronic infection (bottom). (B and C) ClueGO network of DEG in spleen monocytes during acute infection. Each node represents a significantly enriched gene ontology (GO) term and node size is determined by p_adj_. Related GO terms that share > 40 % of genes are connected by a line and organised into functional groups (each given a unique colour). Supergroups are formed when GO terms are shared between more than one group. The names of the top 10 (super)groups (lowest p_adj_) are displayed and (C) shows their proportion of total GO terms. (D - G) Log2FC of (D) signature genes used to predict monocyte differentiation into inflammatory macrophages (MΦ) or monocyte-derived dendritic cells (DC); (E) chemokines, cytokines and their receptors; (F) class I and class II antigen processing and presentation pathways (inc. MHC transactivators*); and (G) T cell costimulation and inhibitory ligands. At each time point, log2FC is shown relative to uninfected controls. Square brackets indicate that common gene names were used. At each time point in (A - G) n = 5-6 for infected mice and n = 6-7 for uninfected controls. *see also Figures S3, S4 and S5*

We next used core lineage signatures to predict the likely outcome of monocyte differentiation in the spleen, where they can be instructed to become either inflammatory macrophages or monocyte-derived dendritic cells (Menezes et al., 2016). This revealed that recruited monocytes initiate a transcriptional programme that is typical of terminally differentiated inflammatory macrophages (Figure 2D). They upregulate their ability to sense parasite- and host-derived danger signals by increasing transcription of diverse pattern recognition receptors (Figure S3B), and upregulate expression of the hallmark cytokines and chemokines associated with a type 1 inflammatory response (Figure 2E). Furthermore, they enhance their capacity to engage T cell immunity by upregulating all major components of the antigen processing and presentation machinery (for class I and class II MHC), and attempt to fine-tune T cell activation by increasing their expression of costimulatory molecules and checkpoint inhibitors (Figures 2F and 2G). Notably, a clear Warburg effect - the metabolic switch from oxidative phosphorylation to glycolysis described when monocytes are stimulated with LPS *in vitro* (Cheng et al., 2014) - was not observed *in vivo* in response to malaria. Instead, the key enzymes involved in both metabolic pathways were transcriptionally induced (Figures S3C and S3D).

We next looked at all of these parameters of monocyte and macrophage biology in our mild model of malaria. Surprisingly, despite substantial differences in parasite density and patterns of sequestration (Figures 1B and S1D), the response of all progenitor and myeloid cells assessed in bone marrow, blood and spleen was remarkably similar between AS and AJ (Figures S4A, S4B and S4C). Moreover, the fate and function of monocytes recruited to the spleen was essentially identical - a direct pairwise comparison identified only a single differentially expressed gene (*Kelch34*) between the two models. Functional gene enrichment analysis using clueGO then revealed that the four largest supergroups identified in monocytes isolated from AJ-infected mice (Figure 2B and 2C) also dominated the response to AS (Figures S4D and S4E). It therefore appears that parasite genotype, pathogen load and the sequestration of infected red cells in immune tissues does not fundamentally alter the myeloid response to acute infection. Instead, the haematopoietic switch that promotes myelopoiesis in the bone marrow and relocates erythropoiesis to the spleen, the disappearance of tissue-resident macrophages and the differentiation of monocytes into inflammatory macrophages may all be part of an emergency response that is unavoidable in a naive host.

### Hosts learn to tolerate chronic infection

Malaria parasites can persist for many months (or even years) in humans (Felger et al., 2012) and the fitness costs of maintaining emergency myelopoiesis over these time frames would be extremely high. We therefore asked how the host adapts to an ongoing infection that can not be cleared. In the chronic phase of *P. chabaudi* the pathogen load can reach up to 1000 parasites per μl blood (Figure 1B) and insoluble malaria pigment accumulates throughout the red pulp of the spleen (Figure 1F). Despite this abundance of parasite-derived signals, the spleen stops extramedullary erythropoiesis and creates new structural demarkations between red and white pulp (Figures 1E and 1F). Furthermore, the bone marrow reduces the production of myeloid progenitors, which in turn reduces the number of inflammatory monocytes and neutrophils trafficking into the blood and spleen (Figures 1D and S2B). At the same time, resident macrophages begin to repopulate their tissue niches (Figure 1G) and ferric iron stores are re-established in the spleen (Figure S2E). These data provide compelling evidence that hosts quickly learn to tolerate chronic infection and return the myeloid compartment towards a healthy uninfected baseline.

In agreement, the transcriptome of recruited monocytes during chronic infection is almost indistinguishable from uninfected controls (Figure 2A). Monocytes are no longer programmed to differentiate into inflammatory macrophages upon their arrival in the spleen (Figure 2D) and they silence the transcriptional signatures of the acute phase response (Figure 2E, 2F, 2G, S3A and S3B). Notably, we find no evidence that they engage mechanisms to suppress inflammation, such as regulatory cytokines (IL-10 and TGFβ) or inhibitors of T cell activation (PD-L1) (Figures 2E and 2G). And furthermore, we find no evidence that they have an alternative fate such as inflammatory hemophagocytes (Figure S3E), which have been implicated in chronic anaemia (Akilesh et al., 2019). Instead, monocytes simply adopt a state of quiescence during chronic infection despite persisting parasitaemia.

Importantly, we can show that quiescence is reversible - when monocytes are removed from the spleen of chronically infected mice and stimulated *in vitro* with LPS their inflammatory response is comparable to monocytes from uninfected mice (Figures S5A and S5B). In both cases, we clearly observe the Warburg effect and transcription of a plethora of inflammatory cytokines, chemokines and co-stimulatory molecules, together with the induction of *Nos2* - the best biomarker of an inflammatory macrophage fate (Figure S5C). Clearly, monocytes are not in a permanent refractory state during chronic infection; instead, their activation and differentiation in response to parasites and their pyrogenic products must somehow be silenced in the remodelled spleen. This likely represents just one of many mechanisms through which the host quickly learns to control inflammation and resolve collateral tissue damage.

### Tolerance persists to protect tissues during reinfection

We next asked whether tolerance could persist in the absence of live replicating parasites to provide long-lived protection. To answer this question, we developed a novel reinfection model that allowed us to exactly match parasite densities between first and second infection. To this end, mice were first infected with the avirulent parasite genotype AS to induce chronic recrudescing parasitaemia and then drug-treated after 40-days of infection to clear circulating and sequestered parasites. One month later, mice were infected for a second time but now with the virulent genotype AJ (Figure 3A). In this model, parasite density (Figure 3B) and the dynamics of red cell loss (Figure 3C) were both matched between first and second infection - this eliminates pathogen load as a confounding factor when analysing acquired mechanisms of disease tolerance.

**Figure 3.**
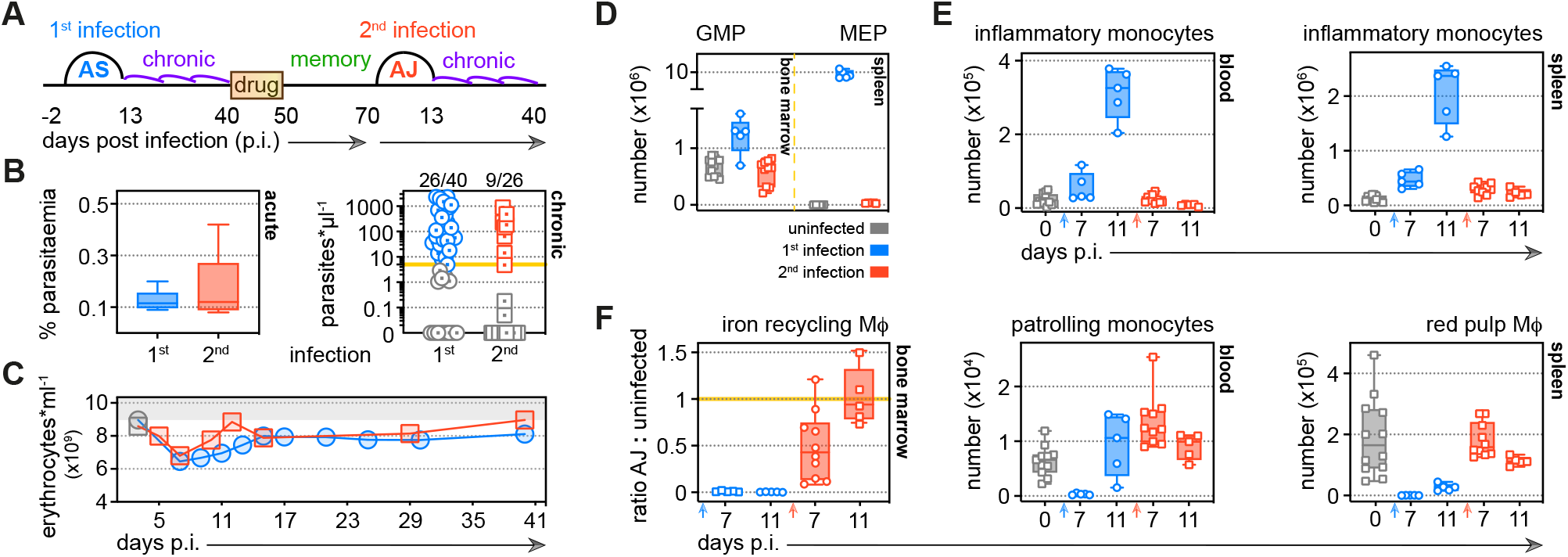
Tolerance persists to protect tissues during reinfection. (A) Malaria reinfection model: C57Bl/6 mice were first infected with *P. chabaudi* AS sporozoites. Chronic infection was confirmed by qPCR at 40-days p.i. and drug-treated with the antimalarial drug chloroquine. 30-days thereafter mice were infected for a second time with *P. chabaudi* AJ. (B) During first and second infection acute parasitaemia was monitored daily using Giemsa stained thin blood films (day 6 p.i. is shown, box-plots display median and IQR) and chronic infection was verified 40-days p.i. by qPCR (n = 40 in 1st infection and n = 26 in 2nd infection). Symbols below the limit of detection (5 parasites*μl^−1^) are coloured grey and these mice were excluded from the study. (C) The mean number of erythrocytes*ml^−1^ is shown before (grey symbols) and during first and second infection (n = 14 per group). (D - F) Inflammatory monocytes, progenitors and tissue-resident macrophages (MΦ) from mice experiencing their first (n = 5 per time point) or second (n = 5-10 per time point) infection were analysed by flow cytometry (box-plots show median and IQR). Uninfected age-matched controls were analysed at each time point and pooled for graphing (n = 12); absolute counts are shown except for tissue-resident MΦ in the bone marrow. In this case, data are presented as a ratio of infected:uninfected at each time point. In (D) granulocyte monocyte progenitors (GMP) and megakaryocytic erythroid progenitors (MEP) are shown 11-days p.i. *see also Figure S6*

In contrast to a first malaria episode, the bone marrow does not prioritise the production of myeloid cells upon reinfection (Figure 3D) and preserves its cellularity and structural integrity (Figure S6A). Consequently, the number of inflammatory monocytes and neutrophils released into circulation does not increase and nor does their accumulation in the spleen (Figures 3E and S6B). Furthermore, the spleen maintains its boundaries between red and white pulp and does not promote extramedullary erythropoiesis (Figures 3D and S6C). And whilst first infection obliterates tissue-resident macrophages in bone marrow, blood and spleen these cells are resistant to malaria-induced ablation second time around (Figure 3F); this allows the host to retain ferric iron stores (Figure S6D). A single malaria episode is therefore sufficient to disarm emergency myelopoiesis in the bone marrow and protect terminally differentiated macrophages in the spleen. And what’s more, tissue architecture is preserved and key homeostatic processes are maintained in tolerised hosts.

### Monocytes minimise inflammation and reduce stress in tolerised hosts

Nevertheless, we reasoned that the myeloid compartment cannot be entirely quiescent during reinfection since mice are able to control replication of the virulent parasite genotype AJ. We therefore isolated spleen monocytes and examined their transcriptional programme in the acute phase of second infection. We identified more than 3000 differentially expressed genes and found that most of these were unique to reinfection (not shared with first infection) (Figure 4A). Remarkably, monocytes did not differentiate into inflammatory macrophages and all functions associated with this fate were silenced (Figures 4B, 4C, S7A, S7B and S7C). Control of inflammation extended beyond the spleen and could be detected systemically with circulating levels of CXCL10 and IFNγ also attenuated at the peak of second infection (Figure 4D). Mammalian hosts can therefore quickly learn to reduce their inflammatory response to malaria parasites.

**Figure 4.**
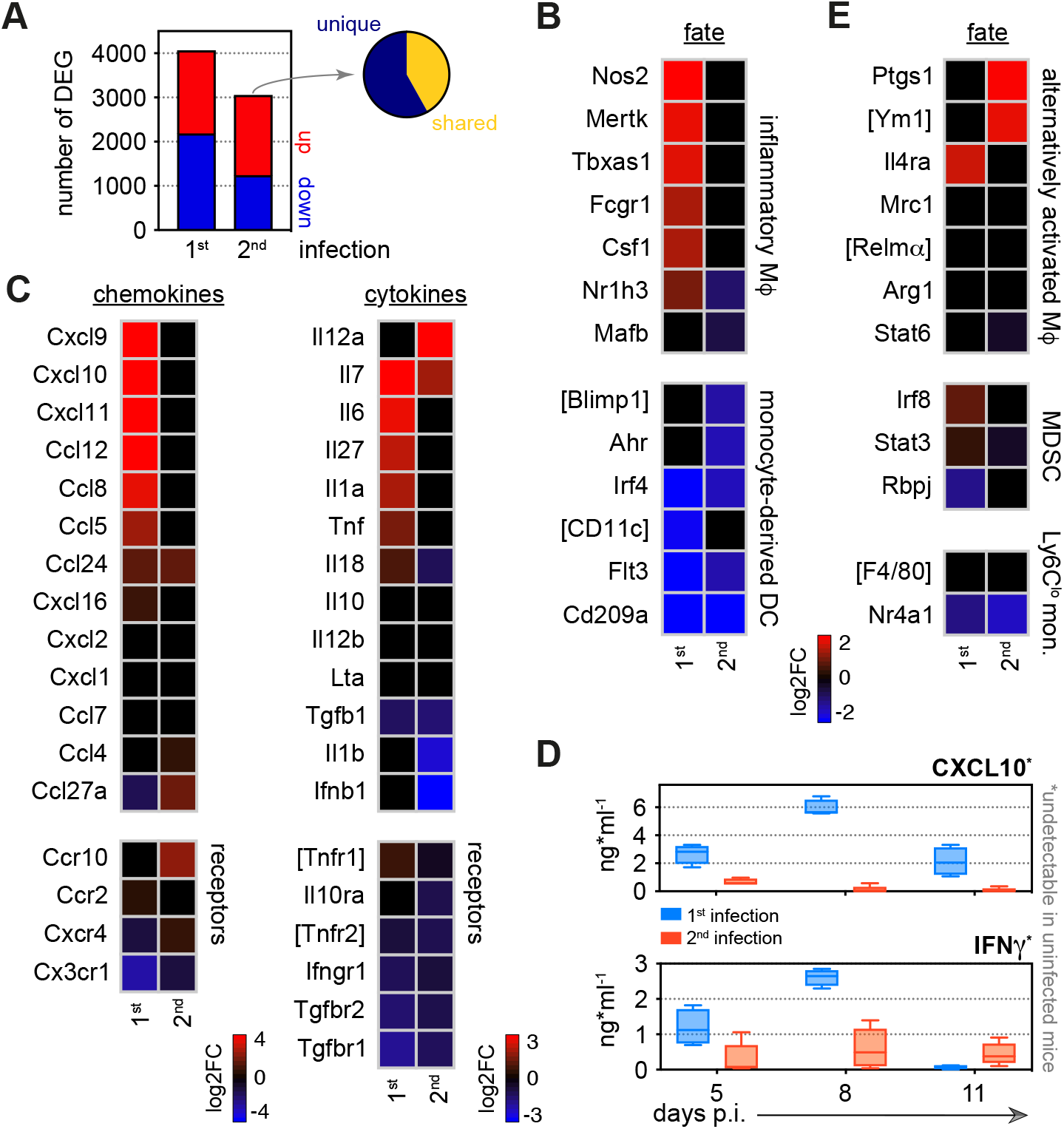
Monocytes minimise inflammation in tolerised hosts. (A) RNA sequencing of spleen monocytes flow-sorted during the acute phase of first or second infection (7-days p.i.). In each case the number of differentially expressed genes (DEG, p_adj_ < 0.01, > 1.5 fold change) was assessed relative to uninfected controls. Pie shows the proportion of DEG unique to second infection. (B - C) Log2 fold change (FC) of (B) signature genes used to predict monocyte differentiation into inflammatory macrophages (MΦ) or monocyte-derived dendritic cells (DC); and (C) chemokines, cytokines and their receptors. Log2FC is shown relative to uninfected controls. (D) Plasma concentration of CXCL10 and IFNγ during the acute phase of first (n = 4) or second (n = 5) infection (box-plots show median and IQR). Note that both plasma proteins were below the limit of detection in uninfected controls. (E) Log2FC of signature genes used to predict monocyte differentiation towards alternative fates (log2FC is shown relative to uninfected controls). In (A - C and E) n = 5 for infected mice and n = 6-7 for uninfected controls. Square brackets indicate that common gene names were used. *see also Figure S7*

We therefore explored alternative monocyte fates, such as those associated with immune regulation, wound healing and tissue repair. However, spleen monocytes were not polarised towards alternatively activated macrophages in second infection and nor did they induce signature genes associated with myeloid-derived suppressor cells (Gabrilovich, 2017) or reparative Ly6C^lo^ monocytes (Jung et al., 2017) (Figure 4E). Furthermore, they did not upregulate anti-inflammatory cytokines or checkpoint inhibitors (Figures 4C and S7C). To make sense of their complex transcriptional profile we therefore turned once more to clueGO. In contrast to first infection, we found minimal enrichment of GO terms linked to host defence; instead, all major supergroups related to regulation of cell cycle and nuclear division, and these terms were unique to second infection (Figures 5A, 5B and 5C). We suggest that proliferation of monocytes in the spleen allows the host to maintain an essential response to reinfection without having to engage increased production and recruitment from the bone marrow.

**Figure 5.**
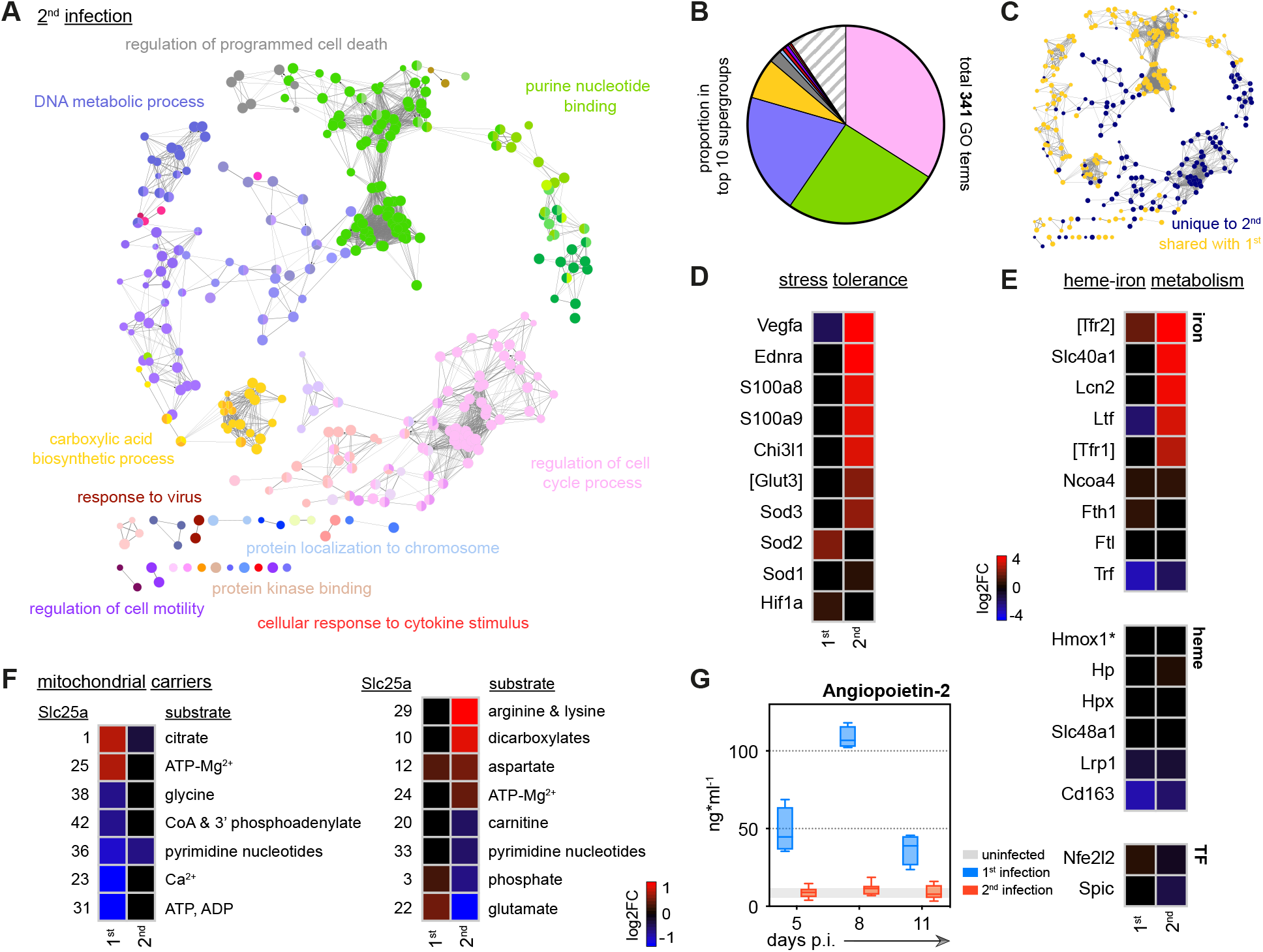
Disease tolerance can be acquired after a single malaria episode. (A and B) ClueGO network of DEG in spleen monocytes during the acute phase of second infection. Each node represents a significantly enriched gene ontology (GO) term and node size is determined by p_adj_. Related GO terms that share > 40 % of genes are connected by a line and organised into functional groups (each given a unique colour). Supergroups are formed when GO terms are shared between more than one group. The names of the top 10 (super)groups (lowest p_adj_) are displayed and (B) shows their proportion of total GO terms. (C) Recoloured clueGO network indicating the GO terms that are unique to second infection (or shared with first infection). (D - F) Log2 fold change (FC) of genes that (D) promote stress tolerance; (E) regulate heme-iron metabolism; and (F) encode mitochondrial carrier proteins. Log2FC is shown relative to uninfected controls. (G) Plasma concentration of Angiopoietin-2 during the acute phase of first (n = 4) or second (n = 8) infection (box-plots show median and IQR). The grey shaded area represents min. to max. measurements in uninfected controls (n = 3). In (A - F) n = 5 for infected mice and n = 6-7 for uninfected controls. Square brackets indicate that common gene names were used.

So what is the function of spleen monocytes in tolerised hosts? First off, they upregulate the expression of two alarmins (*S100a8* and *S100a9*) (Figure 5D) that form the heterodimer calprotectin, which acts as an endogenous TLR4 ligand to silence the inflammatory response to pathogen-derived or damage-associated products, such as free heme. Notably, increased expression of these alarmins in early life prevents hyper-inflammation to protect neonates from sepsis (Ulas et al., 2017). Similarly, *Chi3l1* (encoding the prototypical chitinase-like protein) is also upregulated in second infection and can attenuate inflammasome activation to minimise collateral tissue damage (Dela Cruz et al., 2012). In both cases, these mechanisms promote host control of inflammation without impairing pathogen clearance.

Monocytes also induce transcription of genes that can minimise the effects of hypoxia and oxidative stress on their tissue environment. For example, in first infection monocytes upregulate *Sod2*, which encodes a mitochondrial protein required to safeguard inflammatory macrophages from the reactive oxygen species that they produce. But in second infection they instead increase expression of *Sod3*, which can be secreted to inactivate extracellular reactive oxygen species in the surrounding tissue (Yao et al., 2010) (Figure 5D). In much the same way, monocytes minimise host stress by upregulating *Ednra*, which can scavenge endothelin 1 to prevent vasoconstriction and hypertension (Czopek et al., 2019), and they increase expression of *Vegfa* to promote angiogenesis. Evidently, the transcriptional profile of spleen monocytes suggests that they take on a tissue protective role during second infection.

In support of this argument, monocytes differentially regulate heme-iron metabolism to deal with the release of toxic free heme and reactive iron (Fe^2+^) from ruptured red cells (Figure 5E). In brief, they increase their transcription of a core set of genes required to sequester extracellular iron (*Ltf* encoding lactotransferrin), import sequestered iron for detoxification (*Tfr1/Tfr2* encoding the transferrin receptors) and then export detoxified iron for use in the production of new red cells (*Slc40a1* encoding ferroportin). Notably, monocytes do not appear to increase their iron storage capacity, which has been associated with tissue damage in malaria (Gozzelino et al., 2012). And nor do they upregulate expression of heme oxygenase 1, which is required to detoxify free heme. These data therefore indicate that monocytes specifically enhance iron recycling to promote tolerance to haemolysis - a major source of stress in malaria. Crucially, this only occurs during reinfection, despite identical parasite densities and red cell loss in first and second infection (Figures 3B and 3C).

Taken together, these data clearly show that spleen monocytes initiate a transcriptional programme designed to promote tolerance to malaria parasites upon reinfection. This is achieved in two ways - first by minimising inflammation to reduce collateral tissue damage and second by engaging pathways that can impart stress tolerance on their environment. Significantly, mice were first infected with the avirulent parasite genotype AS, suggesting that parasite-derived signals may be sufficient to redirect monocyte fate.

### Metabolic rewiring underpins monocyte fate

We moved on to ask whether the functional specialisation of monocytes could be underpinned by metabolic reprogramming. Cellular metabolism has emerged as a key determinant of monocyte-to-macrophage differentiation (O’Neill et al., 2016) and so we looked again at transcriptional control of the key enzymes involved in glycolysis and oxidative phosphorylation. We found that both pathways were comparably induced during first and second infection with one notable exception: monocytes switched from upregulating the glucose transporter *Slc2a1* in first infection to *Slc2a3* in second infection (Figures S3C and S7D). *Slc2a3* encodes the facilitative GLUT3 transporter, which unlike most other transporters can continue to import glucose under hypoglycaemic conditions (Simpson et al., 2008). Switching to GLUT3 may therefore constitute a cell-intrinsic adaptation that allows monocytes to tolerate infection-induced stress (low glucose). When we looked deeper into the transcriptional control of cell metabolism, we found that genes encoding mitochondrial carrier proteins was also differentially expressed during reinfection (Figure 5F). These membrane-embedded proteins provide the cellular wiring that connects metabolic reactions in the cytosol with the mitochondrial matrix by transporting metabolites, nucleotides and co-enzymes across the inner mitochondrial membrane (Palmieri, 2013). Mitochondrial carrier proteins thus facilitate the complex crosstalk between the six major metabolic pathways that operate inside immune cells. By re-wiring their mitochondria spleen monocytes may be enabling their specialised tissue protective functions.

### Disease tolerance can be acquired after a single malaria episode

Host control of inflammation may provide a quick and effective way to establish disease tolerance (Medzhitov et al., 2012); thus far, we have shown that inflammation is minimised in the bone marrow (preventing emergency myelopoiesis), blood (attenuating plasma cytokines and chemokines) and spleen (diverting monocyte fate). To directly show that this coincides with a reduction in pathology we measured circulating levels of Angiopoietin-2. This vascular growth factor is the best biomarker of systemic endothelium activation and dysfunction in human malaria and the most accurate prognostic marker of mortality in children (Yeo et al., 2008). We found that in first infection Angiopoietin-2 levels increased by more than an order of magnitude but in second infection - with an identical parasite burden - levels did not deviate from a healthy uninfected baseline (Figure 5G). A single malaria episode can therefore induce host adaptations that promote disease tolerance and generate clinical immunity.

### Specialised monocyte functions are imprinted in the remodelled spleen

So how do hosts learn to control inflammation independently of pathogen load? To begin to answer this question, we looked once again at the functional specialisation of monocytes. Our data are consistent with a model of innate memory, whereby myeloid progenitors in the bone marrow are epigenetically reprogrammed during first infection to intrinsically modify the response of monocytes to reinfection. To test this hypothesis, we asked if malaria induces heritable histone modifications that alter the epigenetic landscape of inflammatory monocytes before their release from the bone marrow. And since tolerance can persist in the absence of parasitaemia, we isolated monocytes from once-infected mice one month after drug cure (day 70, see Figure 1A). Crucially, this was exactly the same time-point at which we had performed all of our reinfection studies (Figure 3A). In this experiment, we interrogated the distribution of histone modifications genome-wide using ChIPseq and asked whether i) transcription start sites were marked with H3K27ac to activate transcription ii) enhancers or superenhancers were marked with H3K4me1 to promote gene expression or iii) DNA was condensed into heterochromatin by H3K9me3 to silence gene expression.

We used the motif discovery software HOMER (Heinz et al., 2010) to identify peaks and visualised them with the genomics exploration tool Integrative Genomics Viewer (Thorvaldsdottir et al., 2013). In the first instance, we looked at the histone modification profiles of genes that define monocyte function in first and second infection; for example, genes associated with inflammation versus stress tolerance. Subscribing to the notion that ChIPseq reveals qualitative (not quantitative) differences (Ma et al., 2018; Orlando et al., 2014) we simply asked whether these genes were marked or not marked. Our prediction was that genes that were upregulated during first infection but silenced during reinfection (tolerised genes) would lose marks associated with active transcription or be condensed into inactive heterochromatin. Conversely, specialised genes (those upregulated exclusively during second infection) would gain marks to promote transcription. Remarkably however, we found that in almost every case the histone modification profiles of tolerised and specialised genes were identical between monocytes isolated from once-infected mice and uninfected controls (Figures 6A, 6B and S8).

**Figure 6.**
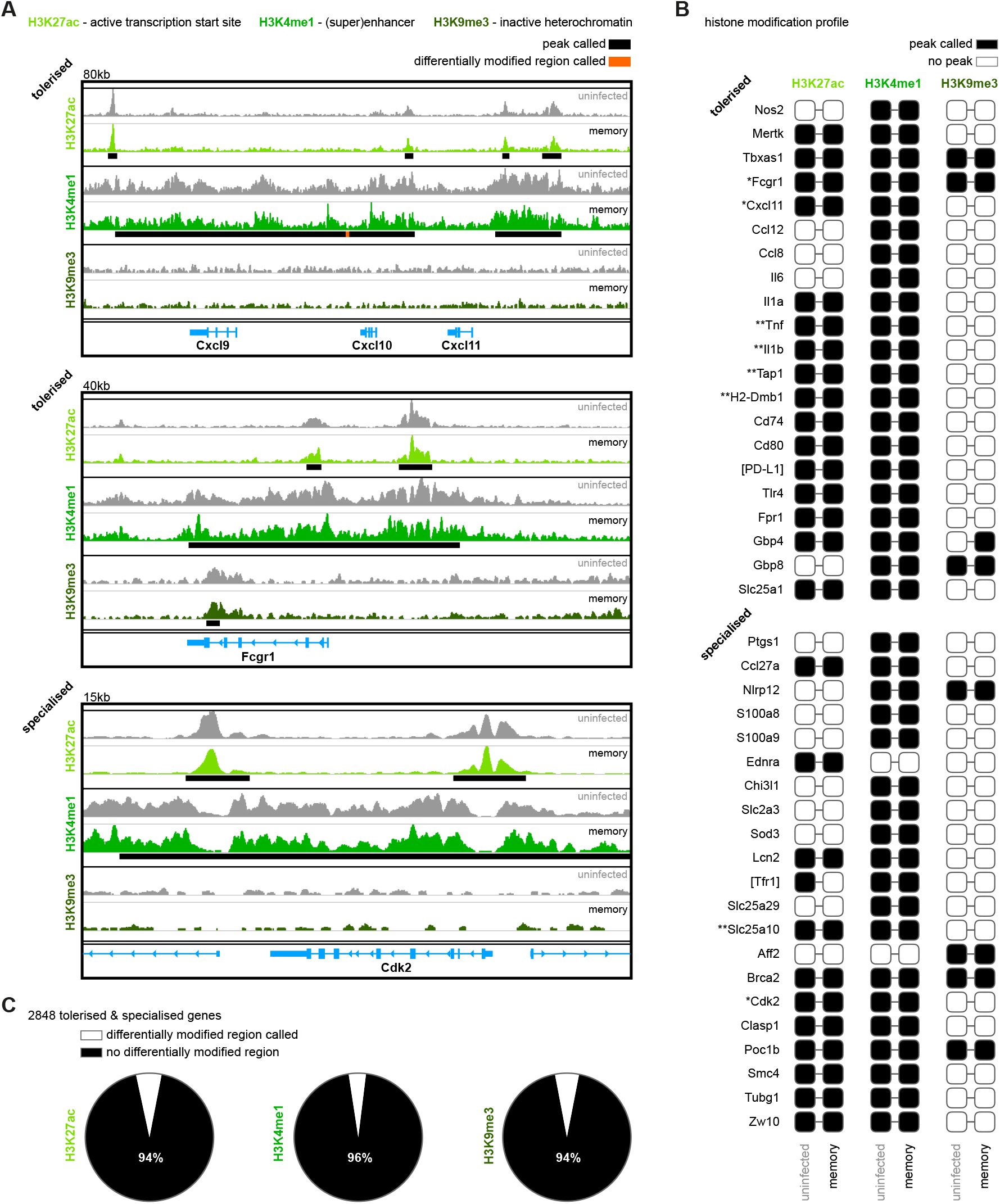
Specialised monocyte functions are imprinted in the remodelled spleen. (A) Chromatin immunoprecipitation (ChIP)seq of bone marrow monocytes flow-sorted from once-infected mice (AJ, memory, 70-days p.i.) and uninfected controls. Shown are the Integrative Genomics Viewer (IGV) traces (autoscaled) of three loci encoding genes that are transcriptionally tolerised (upregulated in first but not second infection) or specialised (upregulated only in second infection). Peaks were called relative to non-immunoprecipitated input DNA and are shown for uninfected controls (black line). Regions of the genome that were differentially marked between once-infected mice and uninfected controls are underlined in orange (differentially modified regions). (B) Histone modification profiles of bone marrow monocytes from once-infected mice and uninfected controls. Black squares indicate that a peak was called within 10 kb (H3K27ac & H3K9me3) or 100 kb (H3K4me1) of the transcription start site. Square brackets indicate that common gene names were used. IGV traces are shown in Figures 6a and S8 for genes marked with one or two asterisks, respectively. (C) Pies show the proportion of tolerised/specialised genes (n = 2848) annotated with or without a differentially modified region (DMR, annotated to the nearest gene). In (A - C) the data shown are pooled from independent biological replicates (see methods). *see also Figure S8 and Table S1*

Even when we used HOMER to call differentially modified regions (DMR) and quantify differences between once-infected and control mice we found little evidence to support epigenetic reprogramming of monocytes - of the 2848 tolerised/specialised genes identified by RNAseq 95 % had no detectable histone modifications (Figure 6C). And in those rare cases where a DMR was called, HOMER assigned a low confidence peak score (Figures 6A and S8, table S1). Innate memory can not therefore easily explain the widespread transcriptional changes that lead to the functional specialisation of monocytes during reinfection. Instead, monocyte fate in tolerised hosts must be imprinted within the remodelled spleen.

## Discussion

In this study we show that mechanisms of disease tolerance can persist after pathogen clearance to minimise tissue damage, stress and pathology during subsequent infections. These inducible mechanisms of tolerance therefore provide memory and constitute an alternative strategy of acquired immunity. Crucially, these adaptations function to preserve key homeostatic processes and protect life, and are therefore likely to be the primary form of host defense when sterile immunity can not be generated. This is particularly relevant in malaria, where even partial control of parasite densities is not usually demonstrable until adolescence (Marsh and Kinyanjui, 2006). It seems likely that many complementary strategies of tolerance will need to cooperate to provide protection and we identify three mechanisms in the myeloid compartment alone - emergency myelopoiesis is disarmed; tissue-resident macrophages become resistant to malaria-induced ablation; and spleen monocytes eschew an inflammatory macrophage fate to take on a tissue protective role. These adaptations operate independently of pathogen load and coincide with a reduction in systemic inflammation; what’s more, tissue integrity is preserved, ferric iron stores are maintained and endothelium activation/dysfunction is avoided during reinfection. Acquired mechanisms of disease tolerance can therefore avert hallmark features of severe malaria.

Host control of inflammation thus appears to provide an effective route to disease tolerance and monocyte activation is clearly not hardwired. However, their functional specialisation in tolerised hosts is not underpinned by epigenetic reprogramming of progenitor cells in the bone marrow but instead is imprinted in the remodelled spleen. Tissue printing of immune cell function is well recognised, and allows monocytes to take on a remarkably diverse range of organ-specific roles. This has been best characterised during the differentiation of monocytes into long-lived tissue resident macrophages, in which local signals imprint specialised functions that are unique to every tissue (Scott 2016, van de Laar 2018). In this way, tissue printing maximises the plasticity of myeloid cells. One of many challenges when resolving the acute phase response to malaria is to refill the red pulp macrophage niche, which was obliterated by acute infection; this is achieved through the recruitment and differentiation of bone marrow-derived monocytes (Lai et al., 2018). We find that red pulp macrophages isolated from once-infected mice are transcriptionally identical to those from uninfected controls (Figure S9), which shows that tissue printing can precisely match the transcriptional programme of recruited monocytes with the functional requirements of the tissue - in this case, to re-establish ferric iron stores. It is clear then that the tissue niche imprints organ-specific identity onto monocytes that take residence to maintain homeostatic processes. But similar mechanisms must also operate to control the fate of monocytes that do not take residence and instead are recruited to deal with injury or infection (Guilliams et al., 2018). In this scenario, how might the tissue niche be remodelled by malaria to promote tolerance?

A niche is often considered to be a small self-contained tissue scaffold that provides the physical structure required to target specialised signals to resident cells or cells transiting the niche. Nevertheless, a niche can also be a dynamic and expansive space, such as the open circulatory system created by the pulp cords and sinuses of the spleen (Guilliams et al., 2020). Malaria causes pronounced splenomegaly and disruption of architecture, including a complete loss of boundaries between red and white pulp. During chronic infection, splenomegaly begins to resolve and these boundaries are re-established but the spleen may not be put back exactly as it was before - it may acquire new and altered features. For example, malaria pigment accumulates throughout the red pulp, and these persistent insoluble deposits may regulate the activation and differentiation of monocytes arriving in the remodelled spleen. In support of this idea, malaria pigment can suppress the oxidative burst (Schwarzer et al., 1992) and interferon-induced upregulation of class II MHC (Schwarzer et al., 1998) in human monocytes *in vitro*. Alternatively, re-assembly of the stromal cell networks that compartmentalise the spleen may alter cell patterning and thereby change the distribution and density of key signals that control monocyte fate (Bonnardel et al., 2019; Koliaraki et al., 2020). Or perhaps the tissue niche required to promote tolerance is already present in the spleen (even in a naive host) but is lost when the spleen becomes enlarged and disorganised in first infection. In this example, it is the preservation of spleen architecture that facilitates the functional specialisation of monocytes in tolerised hosts. In every scenario, monocytes arriving in the remodelled spleen receive a new combination of signals that can minimise inflammation and induce their tissue protective functions. Importantly, tissue remodelling and the accumulation of malaria pigment is observed in the human spleen (Buffet et al., 2011), and we therefore suggest that this is a conserved strategy through which malaria imprints tolerance. This will almost certainly benefit the parasite as well as the host; after all, if you can minimise host sickness behaviour you will likely maximise the probability of onward transmission.

The spleen is not the only lymphoid tissue that promotes tolerance during reinfection; we find that emergency myelopoiesis is disarmed in the bone marrow. How is this avoided second time around despite an identical pathogen load? It could be that changes to the tissue niche promote tolerance, in much the same way as described in the spleen, as first infection leads to a loss of cellularity and tissue integrity in the bone marrow, which inevitably requires repair. But it is also possible that cell-intrinsic changes establish tolerance in this case. Current evidence indicates that systemic inflammation, which can push haematopoietic stem cells (HSC) down the myeloid path (Mitroulis et al., 2018), is triggered in the bone marrow by plasmacytoid dendritic cells (pDC), which produce type I interferon (Spaulding et al., 2016). Can you intrinsically modify pDC to attenuate their function (e.g. by increasing their activation threshold)? Or limit their access to parasites and their pyrogenic products? Or perhaps tolerance operates further upstream - the expression of pattern recognition receptors on HSC could be restricted to prevent their direct activation by malaria parasites. Alternatively, tolerance may require changes outwith the myeloid compartment, for example the generation of memory T cells that can actively suppress interferon signaling. Whatever the mechanism we believe that learning to disarm emergency myelopoiesis in the bone marrow is just as important as learning to imprint specialised functions on recruited monocytes in the spleen. Moreover, we argue that tolerance in the bone marrow is actually supported by tissue printing in the spleen; proliferation of monocytes is unique to second infection and could allow the host to maintain a critical tissue-specific response without having to engage increased production in the bone marrow.

The final mechanism of tolerance that we identify is the resistance of tissue-resident macrophages to malaria-induced ablation. Tissue-resident macrophages were often considered to be immune sentinels that provide a first line of defence against invading pathogens but actually they have been shown time and again to be sensitive to stress, and their disappearance in response to injury or infection is well recognised (Aegerter et al., 2020; Bleriot et al., 2015; Machiels et al., 2017; Scott et al., 2016). Their functions instead mainly relate to development, tissue repair and homeostasis. So how does malaria cause the death of tissue-resident macrophages in naive hosts and why are they protected in tolerised hosts? We suggest that the release and catabolism of free heme from ruptured red cells can explain this phenomenon. Free heme is toxic to tissue-resident macrophages (Cambos and Scorza, 2011; Ferreira et al., 2008; Haldar et al., 2014; Theurl et al., 2016) and the detoxification of free heme is known to confer disease tolerance (Seixas et al., 2009). In our model, heme accumulation is likely to be similar between first and second infection as the number of circulating parasites and red cell loss is comparable. The preservation of tissue-resident macrophages could therefore be a consequence of learning to more efficiently detoxify free heme to reduce its bioavailability and pathogenicity. Since we find no evidence that monocytes transcriptionally regulate their capacity to metabolise heme during reinfection (they exclusively upregulate iron recycling) other cell types would need to carry out this role, including hepatocytes and renal proximal tubule epithelial cells (Ramos et al., 2019; Theurl et al., 2016). In this scenario, metabolic adaptations that operate in non-lymphoid organs must also be inducible, and would persist after pathogen clearance to provide memory and clinical immunity.

Acquired immunity to malaria is often considered to be slow and ineffective - but this is only true if we consider resistance (the ability to eliminate parasites) in isolation. Despite being an equally important part of host defense, the central role of disease tolerance in acquired immunity to malaria has not been fully appreciated. We demonstrate that disease tolerance can be attained after a single malaria episode and can persist in the absence of parasitaemia to allow the host to tolerate subsequent infections. We therefore argue that mechanisms of disease tolerance provide an alternative strategy of acquired immunity that functions not to kill parasites but to limit the damage that they can cause. Adaptations in the myeloid compartment minimise inflammation and promote stress tolerance, and it is likely that these will need to be complemented by adaptations in other immune compartments such as memory T cell reservoirs, which orchestrate innate and adaptive immunity. Furthermore, changes to the immune response will need to be complemented by long-lived metabolic adaptations that maintain homeostasis and preserve organ function. Together, all of these adaptations can work in concert to control inflammation, protect host tissues and minimise pathology. The mechanisms of disease tolerance uncovered in this study are therefore likely just the tip of the iceberg but they can begin to explain how children in endemic areas acquire immunity to severe malaria so quickly and without the need for improved pathogen control (Goncalves et al., 2014; Gupta et al., 1999; Marsh and Snow, 1999).

## Acknowledgements

We would like to thank Ronnie Mooney for the production of Anopheles stephensi mosquitoes and for assistance with in vivo experiments. Flow cytometry data were generated within the cell sorting facility in Ashworth laboratories (University of Edinburgh), which is supported by funding from the Wellcome Trust. RNA sequencing libraries were prepared and sequenced by the Edinburgh Clinical Research Facility at the University of Edinburgh, which receives financial support from NHS Research Scotland (NRS). ChIPseq libraries were sequenced by Edinburgh Genomics, which is supported through core grants from NERC (R8/H10/56), MRC UK (MR/K001744/1) and BBSRC (BB/J004243/1). We thank the Shared University Research Facilities at Little France for histology and Hologic Ltd. for microarray services. This project was supported by the Wellcome Trust-University of Edinburgh Institutional Strategic Support Fund, and PJS is the recipient of a Sir Henry Dale Fellowship jointly funded by the Wellcome Trust and the Royal Society (grant no. 107668/Z/15/Z). We are grateful to Steve Jenkins, Eleanor Riley and Joanne Thompson for their thoughtful comments on our manuscript.

## Author contributions

**WN** & **PJS** conceptualisation, methodology, investigation, writing (draft and review); **WN** & **AI** & **PJS** formal analysis; **WN** visualisation; **AI** software, data curation, writing (review); **PJS** supervision, funding acquisition.

## Supplementary table legend

**Table S1. Quantitative changes in the histone modification profiles of once-infected mice**

Bone marrow monocytes were flow-sorted from once-infected mice (AJ, memory, 70-days p.i.) and uninfected controls for chromatin immunoprecipitation (ChIP)seq; differences in their histone modification profiles were then quantified by calling differentially modified regions (DMR, annotated to the nearest gene). Shown is a list of all tolerised/specialised genes annotated with a DMR, ordered by peak score.

*related to Figure 6*

## Supplementary Figures

**Figure S1.**
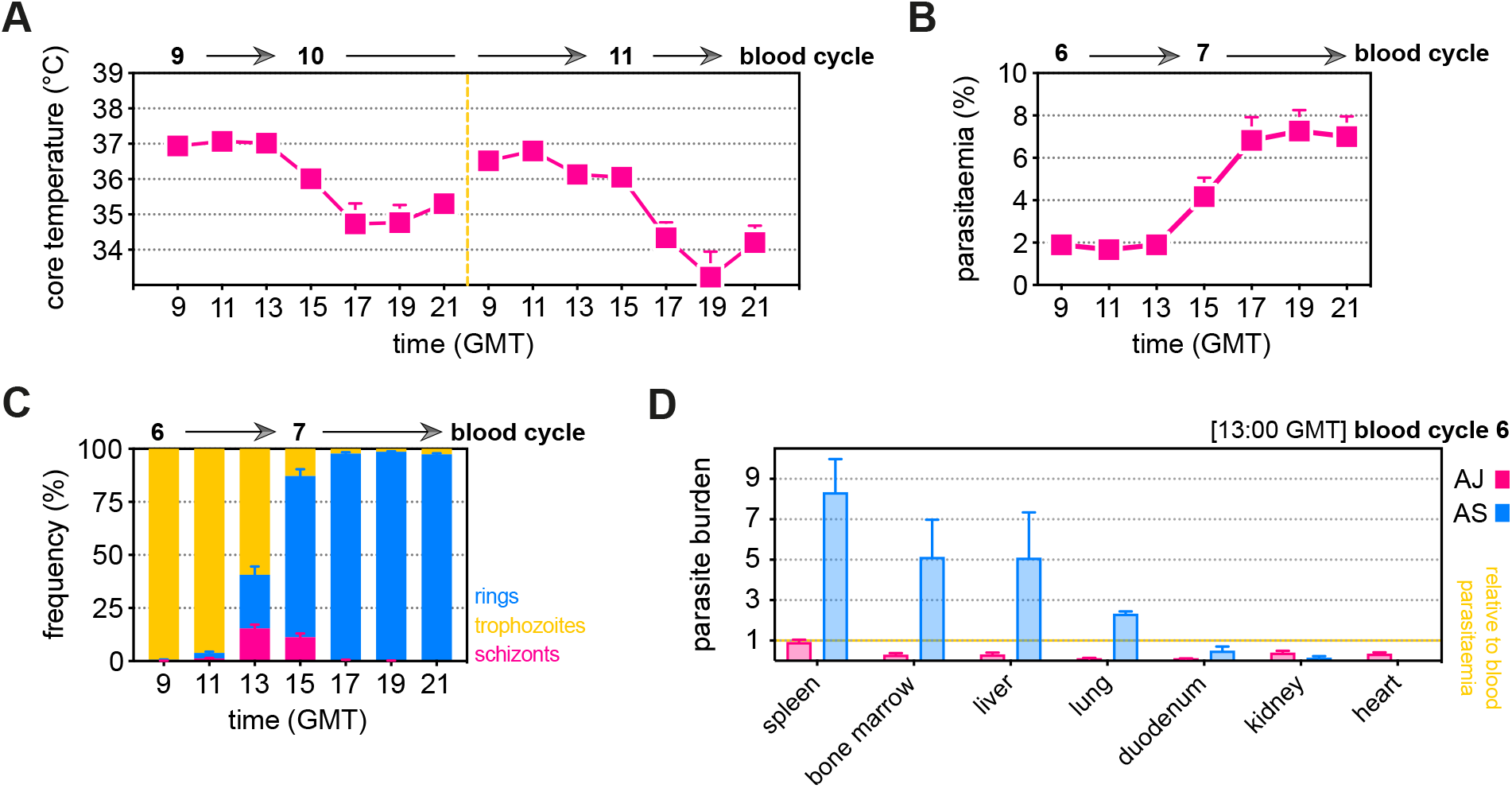
*P. chabaudi* AJ causes severe disease without sequestering in host tissues. (A - C) C57Bl/6 mice were infected with *P. chabaudi* AJ sporozoites; the blood-stage of infection started 2-days later after the release of merozoites from the liver. We refer to the emergence of parasites from the liver as the start of blood cycle 1, after which parasites undergo schizogony approximately every 24-hours to start the next blood cycle. Mice were housed under reverse light conditions (lights OFF 07:00, lights ON 19:00 GMT) so that schizogony would peak at 13:00 GMT. (A) Core body temperature was measured every 2-hours (09:00 - 21:00 GMT) as parasites transitioned from blood cycle 9 to 10 and again from blood cycle 10 to 11 (n = 20, mean+SEM). (B and C) Parasitaemia was monitored every 2-hours by Giemsa stained thin blood films (09:00 - 21:00 GMT) as parasites transitioned from blood cycle 6 to 7 (n = 9, mean+SEM). The percentage of infected red cells is shown in (B) whereas the proportion of parasites at the ring, trophozoite and schizont stages is shown in (C). Note that hypothermia is most severe after the peak of schizogony when all schizonts have ruptured. (D) C57Bl/6 mice were infected with mosquito transmitted blood-stage parasites (*P. chabaudi* AS or AJ). Circulating parasitaemia was measured in blood at the peak of schizogony (13:00 GMT) as parasites transitioned from blood cycle 6 to 7 (n = 3 per group, mean+SEM). At the same time, mice were euthanised and organs were fixed in neutral buffered formalin for histology. Sequestration rates in each organ were assessed by counting the percentage of infected red cells inside blood vessels and normalising this number to circulating parasitaemia. *related to Figure 1*

**Figure S2.**
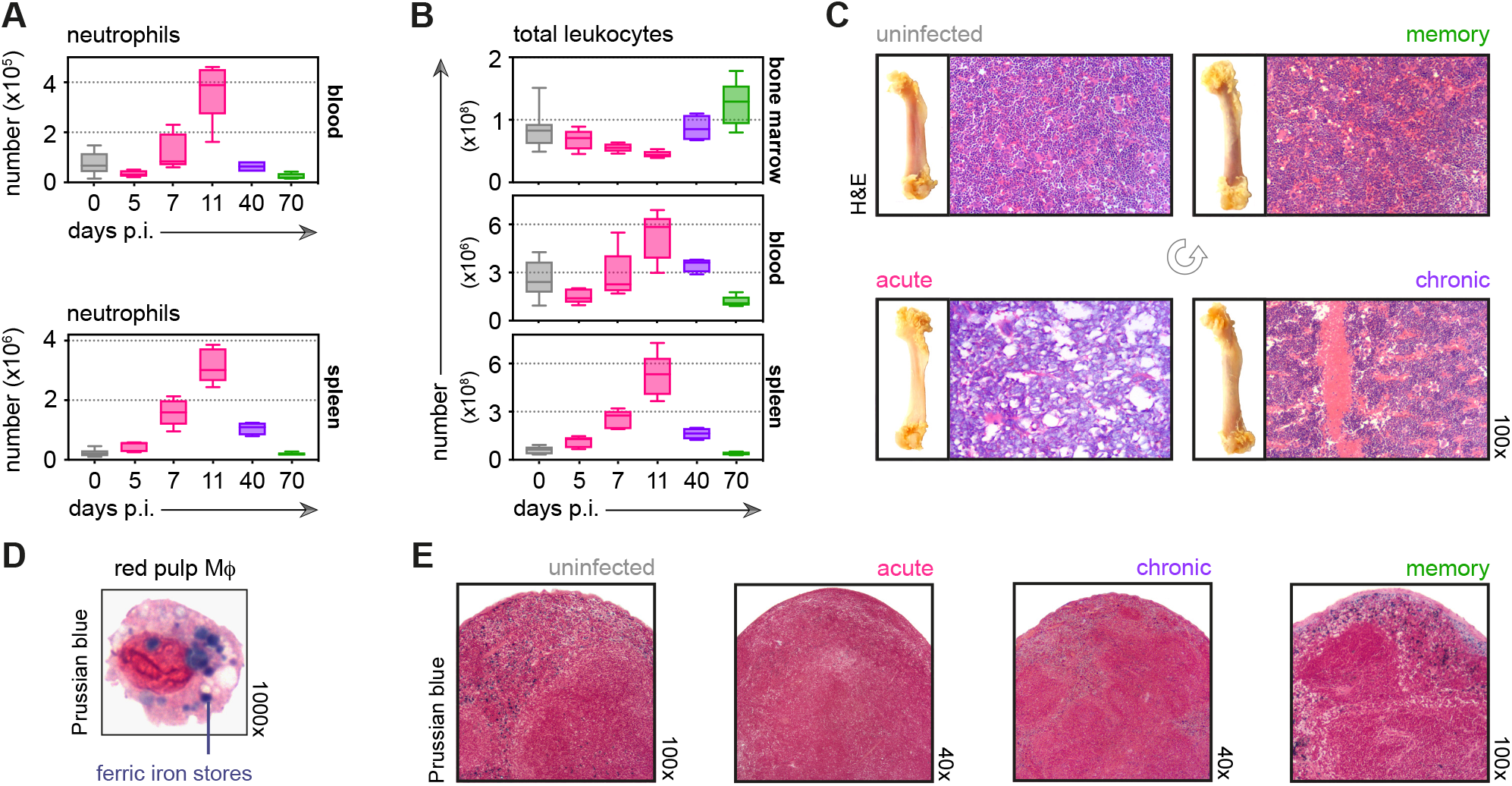
Malaria causes major disturbances in tissue structure and integrity. (A and B) Neutrophils (A) and total leukocytes (B) from uninfected mice (0-days p.i.), AJ infected mice (5, 7, 11 and 40-days p.i.) and once-infected mice (memory, 70 days p.i.) were analysed by flow cytometry (n = 4-5 per time point, box-plots show median and IQR). Uninfected age-matched controls were analysed at each time point and pooled for graphing (n = 10). (C) Paraffin embedded femur sections were H&E stained (11-days p.i. for acute AJ infection) - note that during acute infection bones appear translucent, as cellularity drops. (D) Red pulp macrophages (MΦ) were flow-sorted from uninfected control mice and stained with Prussian Blue (intracellular ferric iron stores) and Neutral Red (nuclei). (E) Paraffin embedded spleen sections were stained with Prussian Blue and Neutral Red (11-days p.i. for acute AJ infection). *related to Figure 1*

**Figure S3.**
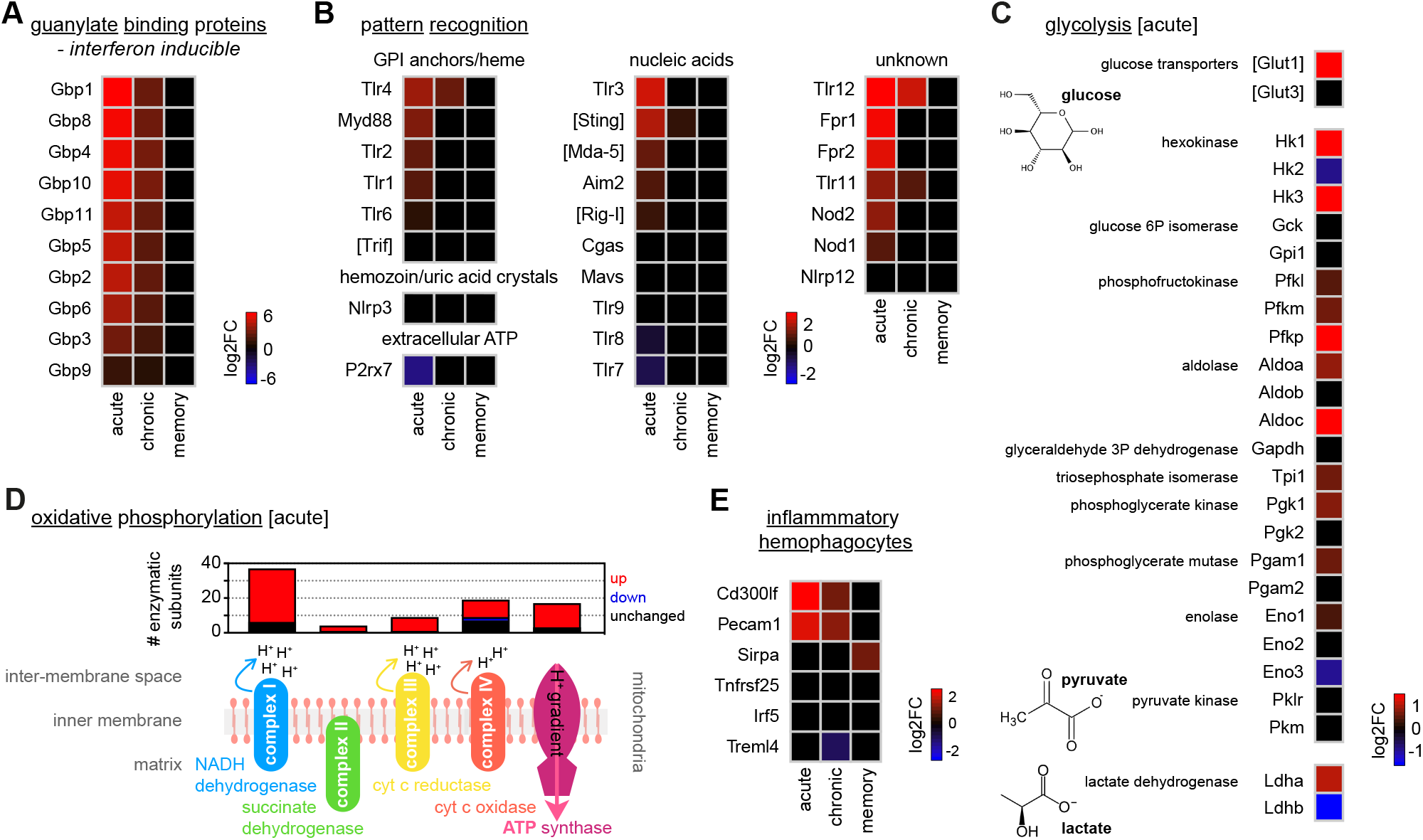
Monocytes upregulate glycolysis and oxidative phosphorylation as they differentiate into inflammatory macrophages. (A and B) RNA sequencing of spleen monocytes flow-sorted from AJ infected mice (7 and 40-days p.i. for acute and chronic, respectively) and once-infected mice (memory, 70-days p.i.). Shown is the log2 fold change (FC) of (A) interferon-inducible guanylate binding proteins and (B) pattern recognition receptors (PRR). At each time point, log2FC is shown relative to uninfected controls. If parasite- or host-derived ligands have been identified for PRR in malaria these ligands are labelled. (C) Log2FC of the major glucose transporters and glycolytic enzymes in spleen monocytes flow-sorted from AJ infected mice (acute, 7-days p.i.) - data are shown relative to uninfected controls. Note that under anaerobic conditions pyruvate can be further converted to lactate by lactate dehydrogenase, which is also shown. (D) Transcriptional regulation of oxidative phosphorylation in spleen monocytes flow-sorted from AJ infected mice (acute, 7-days p.i.). Data show the number of enzymatic subunits that are transcriptionally up- or downregulated compared to uninfected controls (p_adj_ < 0.01, > 1.5 fold change) - all subunits that are required to form complex I to IV in the electron transport chain and ATP synthase are shown. (E) Log2FC of signature genes used to predict monocyte differentiation into inflammatory hemophagocytes; data are shown relative to uninfected controls. At each time point in (A - E) n = 5-6 for infected mice and n = 6-7 for uninfected controls. Square brackets indicate that common gene names were used. *related to Figure 2*

**Figure S4.**
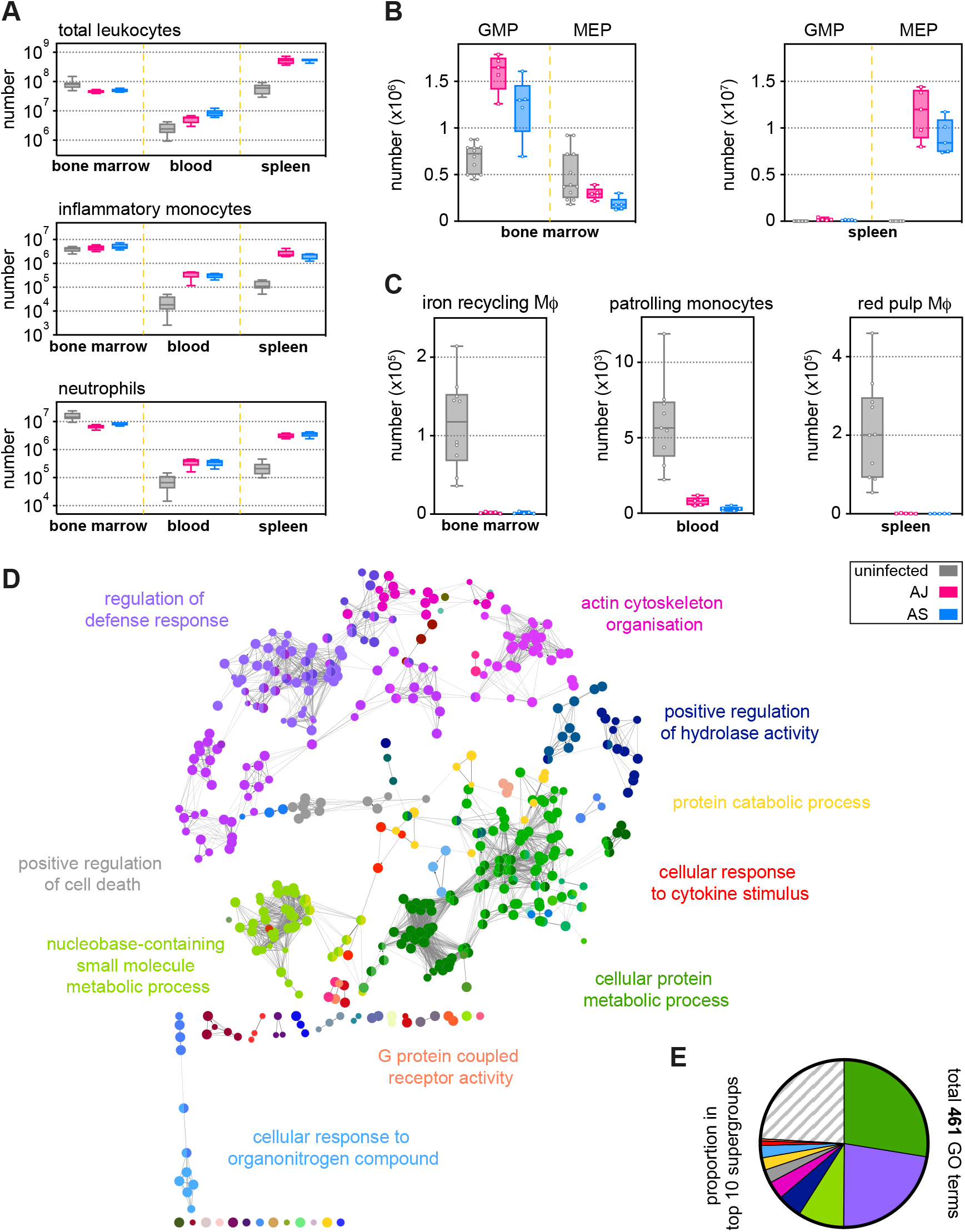
Parasite genotype does not influence the emergency myeloid response to malaria. (A - C) Inflammatory monocytes (A), progenitors (B) and tissue-resident macrophages (MΦ) (C) from uninfected mice (n = 10), AJ infected mice (n = 5) and AS infected mice (n = 5) were analysed by flow cytometry (box-plots show median and IQR). Data shown represent the peak of the acute phase response, which is (A) 11-days p.i. for total leukocytes, inflammatory monocytes and neutrophils in bone marrow, blood and spleen; (B) 7-days p.i. for granulocyte monocyte progenitors (GMP) and megakaryocyte erythroid progenitors (MEP) in the bone marrow and 11-days p.i. for progenitors in the spleen; and (C) 7-days p.i. for tissue-resident MΦ. (D and E) ClueGO network of differentially expressed genes (DEG) in spleen monocytes flow-sorted from AS infected mice (acute, 7-days p.i., n = 5). DEG were called relative to uninfected controls (n = 7). Each node represents a significantly enriched gene ontology (GO) term and node size is determined by p_adj_. Related GO terms that share > 40 % of genes are connected by a line and organised into functional groups (each given a unique colour). Supergroups are formed when GO terms are shared between more than one group. The names of the top 10 (super)groups (lowest p_adj_) are displayed and (E) shows their proportion of total GO terms. *related to Figure 2*

**Figure S5.**
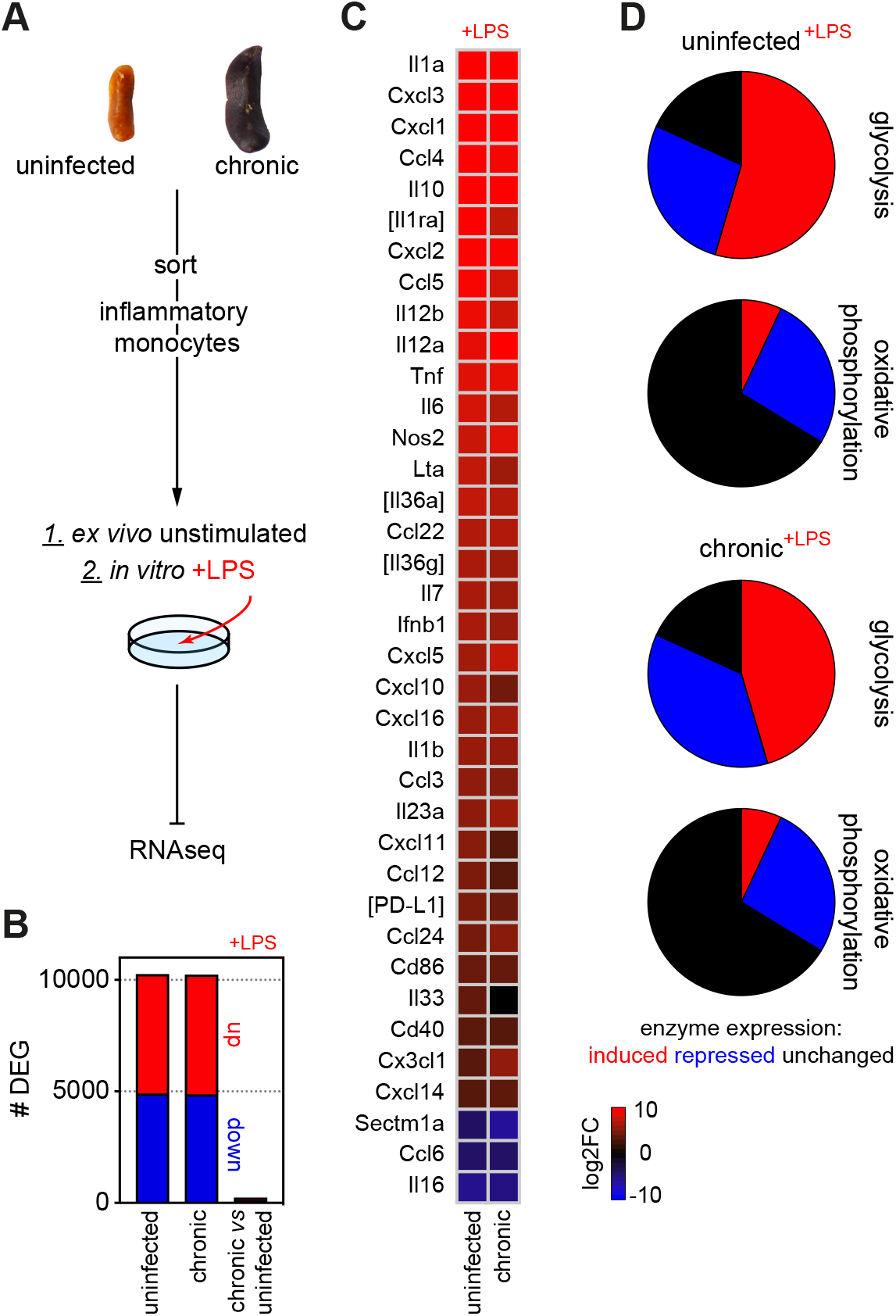
Quiescent monocytes can differentiate into inflammatory macrophages when removed from the spleen. (A) Spleen monocytes were flow-sorted from uninfected mice (n = 4) or mice chronically infected with *P. chabaudi* AJ (40-days p.i., n = 5). Sorted cells were then prepared immediately for RNA sequencing (*ex vivo*) or stimulated *in vitro* with LPS for 4-hours before RNA sequencing. (B) The number of differentially expressed genes (DEG, p_adj_ < 0.01, > 1.5 fold change) between LPS stimulated and unstimulated (*ex vivo*) monocytes is shown for uninfected mice (left bar) and chronically infected mice (centre bar). Note that a direct comparison between LPS stimulated monocytes from chronically infected mice and uninfected controls yielded only 229 DEG (right bar). (C) Log2 fold change (FC) of inflammatory markers showing the differentiation of monocytes *in vitro* after LPS stimulation (data shown relative to *ex vivo* samples). Square brackets indicate that common gene names were used. (D) Transcriptional regulation of glycolysis and oxidative phosphorylation after LPS stimulation (relative to *ex vivo* samples). Data include the major glucose transporters and glycolytic enzymes, and all enzymatic subunits that form complex I to IV in the electron transport chain and ATP synthase. *related to Figure 2*

**Figure S6.**
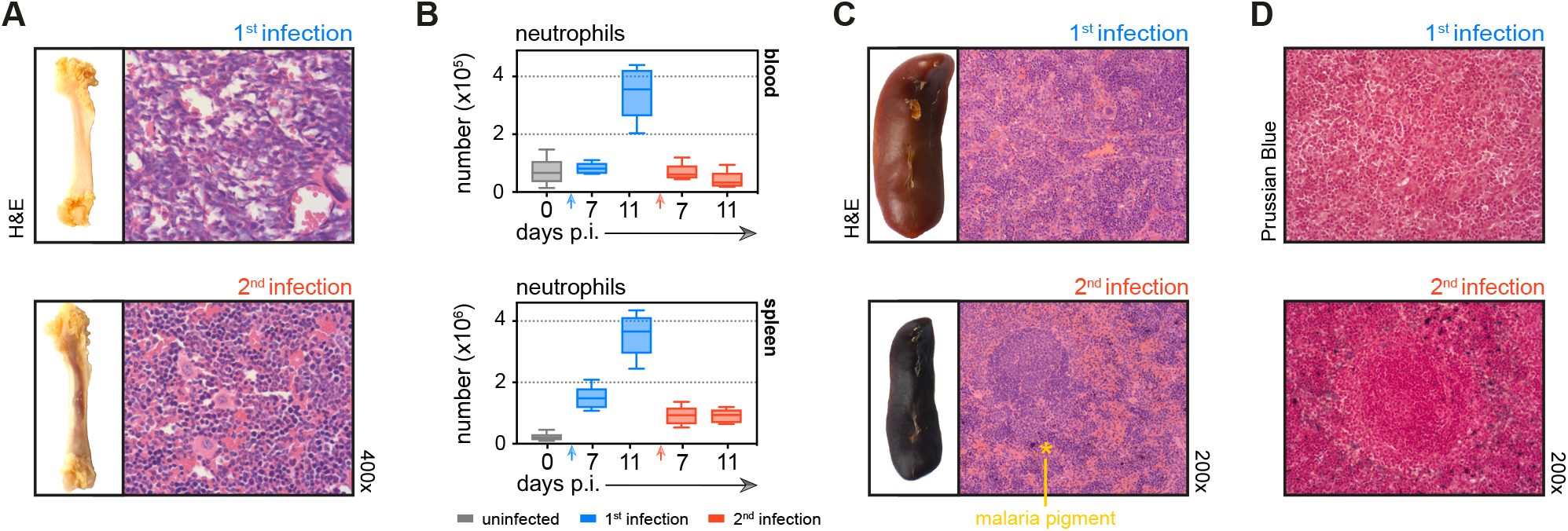
Tissue architecture is preserved and key homeostatic processes are maintained in tolerised hosts. (A) Paraffin embedded femur sections were H&E stained during the acute phase of first or second infection (11-days p.i.). (B) Neutrophils from mice experiencing their first (n = 5 per time point) or second (n = 5-10 per time point) infection were analysed by flow cytometry (box-plots show median and IQR). Uninfected age-matched controls were analysed at each time point and pooled for graphing (n = 12). (C and D) Paraffin embedded spleen sections were H&E (C) or Prussian Blue/Neutral Red (D) stained during the acute phase of first or second infection (11-days p.i.). Note that malaria pigment deposited during first infection persists and can still be observed during reinfection (marked with an asterisk). *related to Figure 3*

**Figure S7.**
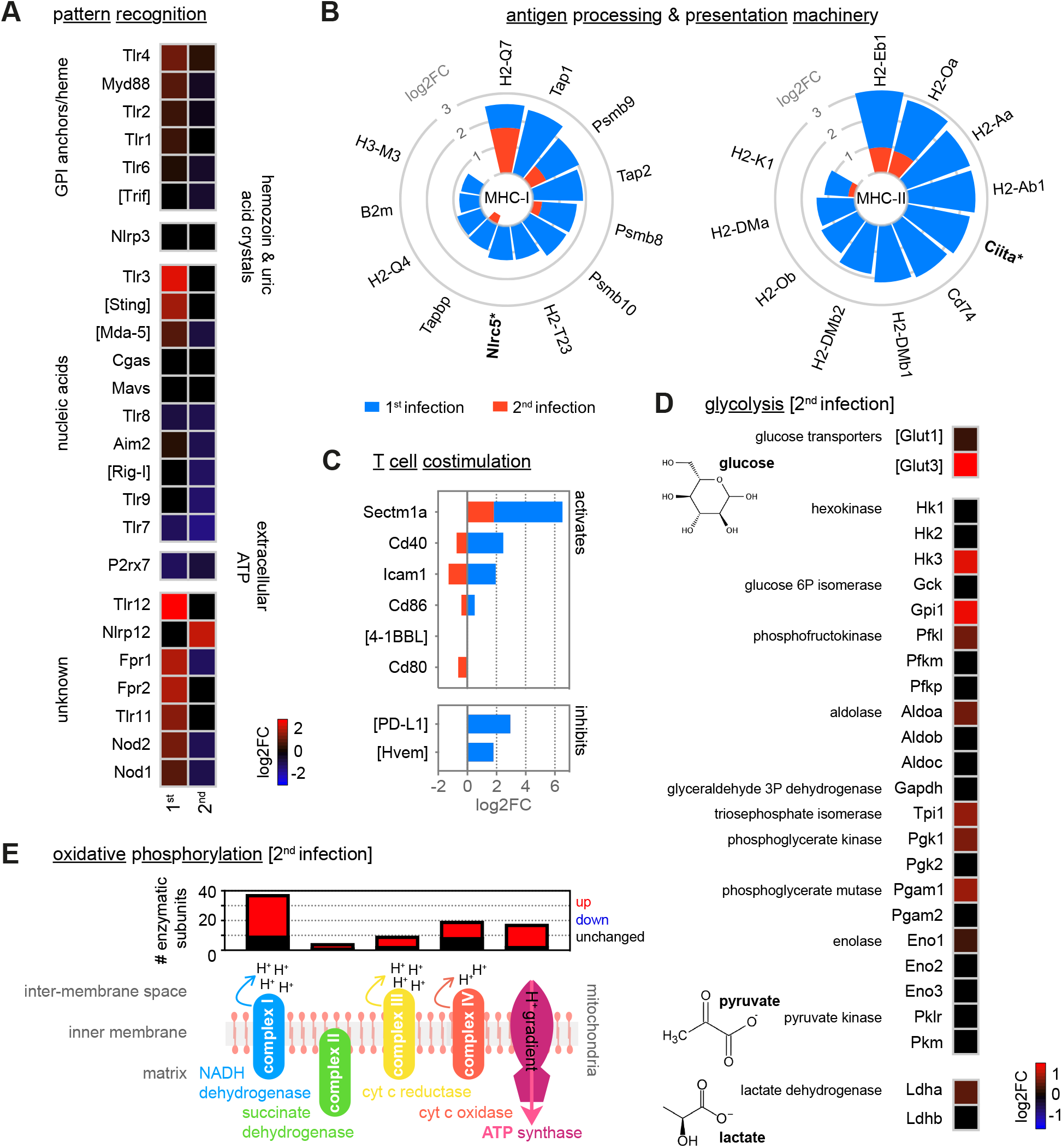
Monocytes minimise inflammation in second infection. (A - C) RNA sequencing of spleen monocytes flow-sorted from mice experiencing their first or second infection (acute, 7-days p.i.). Shown is the log2 fold change (FC) of (A) pattern recognition receptors (PRR); (B) class I and class II antigen processing and presentation pathways (inc. MHC transactivators*); and (C) T cell costimulation and inhibitory ligands. Data are shown relative to uninfected controls. If parasite- or host-derived ligands have been identified for PRR in malaria these ligands are labelled. (D) Log2FC of the major glucose transporters and glycolytic enzymes in spleen monocytes during the acute phase of second infection (7-days p.i.) - data are shown relative to uninfected controls. Note that under anaerobic conditions pyruvate can be further converted to lactate by lactate dehydrogenase, which is also shown. (E) Transcriptional regulation of oxidative phosphorylation in spleen monocytes during the acute phase of second infection (7-days p.i.). Data show the number of enzymatic subunits that are transcriptionally up- or downregulated compared to uninfected controls (p_adj_ < 0.01, > 1.5 fold change) - all subunits that are required to form complex I to IV in the electron transport chain and ATP synthase are shown. In (A - E) n = 5 for infected mice and n = 6-7 for uninfected controls. Square brackets indicate that common gene names were used. *related to Figure 4*

**Figure S8.**
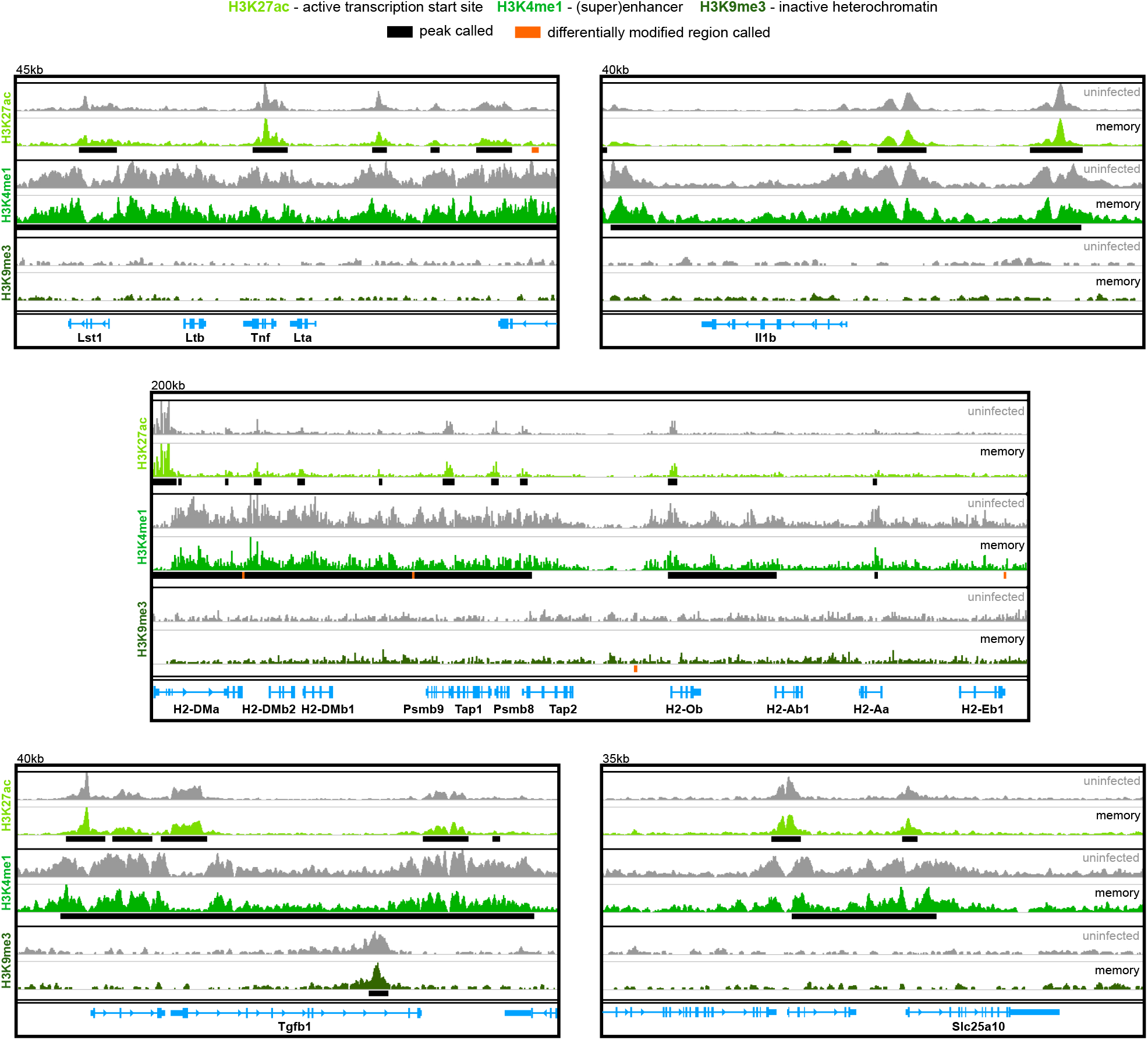
Epigenetic reprogramming does not explain the specialised functions of monocytes in tolerised hosts. Chromatin immunoprecipitation (ChIP)seq of bone marrow monocytes flow-sorted from once-infected mice (AJ, memory, 70-days p.i.) and uninfected controls. Shown are the Integrative Genomics Viewer (IGV) traces (autoscaled) of five loci encoding immune and metabolic genes. Peaks were called relative to non-immunoprecipitated input DNA and are shown for uninfected controls (black line). Regions of the genome that were differentially marked between once-infected mice and uninfected controls are underlined in orange (differentially modified regions). The data shown are pooled from independent biological replicates (see methods). *related to Figure 6*

**Figure S9.**
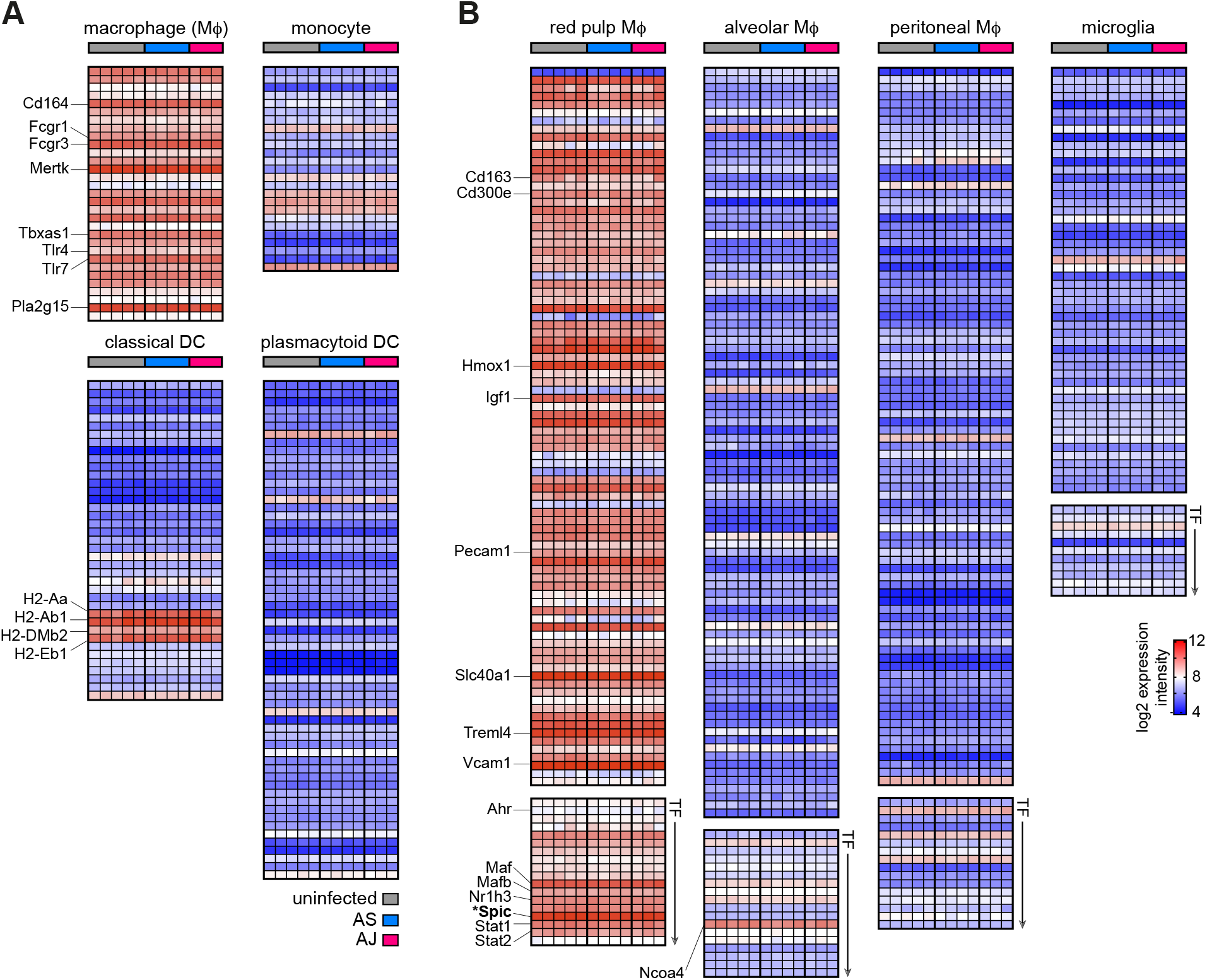
Tissue printing shapes the transcriptional programme of recruited monocytes to meet the needs of the niche. (A and B) Microarray of red pulp macrophages (MΦ) flow-sorted from the spleens of once-infected mice (memory, 100-days p.i.) and age-matched uninfected controls. Data are displayed as the RMA (robust multi-array average) normalised log2 expression intensity for each gene. Details of how we curated signature genelists for (A) mononuclear phagocytes (inc. dendritic cells, DC) and (B) tissue resident MΦ are provided in the methods. *Spic* is the master transcription factor (TF) for red pulp MΦ fate (Kohyama et al., 2009) and is marked with an asterisk. Note that pairwise comparisons between each of the three groups (uninfected, AS and AJ) revealed zero differentially expressed genes (p_adj_ < 0.05) (n = 5 for uninfected mice and n = 3-4 for once-infected mice). *related to discussion*

## Methods

### Mice

All animal experiments were conducted in accordance with UK Home Office regulations (Animals Scientific Procedures Act 1986; project licence number 70/8546) and approved by veterinarian services at the University of Edinburgh. C57Bl/6J mice, originally obtained from the *Jackson Laboratory*, were bred and housed in individually ventilated cages under specific pathogen free conditions. Mice had access to water and rat & mouse no. 3 breeding diet (*Special Diets Services*) at all times. Experimental procedures were initiated when mice were 8 - 10 weeks of age, following acclimatisation to a reversed 12-hour dark/light cycle (lights OFF at 07:00 GMT and lights ON at 19:00 GMT). Mice were culled either by cervical dislocation or by pentobarbital overdose followed by exsanguination.

### Mosquito transmission of malaria parasites

We transmitted two serially blood passaged *Plasmodium chabaudi chabaudi* clones (*P. chabaudi* AS (clone 28AS11) and *P. chabaudi* AJ (clone 96AJ15)), which were originally obtained from the University of Edinburgh. *Anopheles stephensi* mosquitoes (strain SD500) were reared in-house and infected with *P. chabaudi* according to our previously published protocol (Spence et al., 2012). In brief, donor mice were inoculated with serially blood passaged *P. chabaudi* by intraperitoneal injection of infected red cells. Gametocytes were quantified on day 14 of infection and mice with > 0.1 % gametocytaemia were anaesthetised and exposed to female mosquitoes at 11:00 GMT. To ensure optimal parasite development, mosquitoes were kept at 26.0°C (± 0.5°C) in an ultrasonic humidity cabinet and provided with 8 % Fructose and 0.05 % 4-Aminobenzoic acid (both *Sigma*) feeding solution from this point forward. Successful oocyst development was verified in mosquito midguts 8-days later and sporozoites were isolated from mosquito salivary glands on day 15 post-feed. Salivary glands were dissected under a stereomicroscope and transferred to a glass mortar; to maintain sporozoite viability, salivary glands were kept on ice in RPMI supplemented with 0.2 % Glucose, 0.2 % Sodium bicarbonate (both *Sigma*), 2 mM L-Glutamine (*Gibco*) and 10 % fetal bovine serum (*Gibco*, FBS Performance Plus, heat inactivated & filtered 0.22 μm) for a maximum of 2-hours. Sporozoites were released from salivary glands by gentle homogenisation and washed three times before enumeration. To initiate infection in experimental mice 200 *P. chabaudi* sporozoites were intravenously injected into the tail vein.

### Monitoring the course and outcome of infection

Following sporozoite injection, *P. chabaudi* develops in the liver for 52-hours before the release of merozoites to kick-start asexual blood-stage replication - each blood cycle takes approximately 24-hours to complete. Mice were closely monitored for the first 14-days of acute blood-stage infection with parasitaemia quantified daily using Giemsa stained thin blood films (counting at least 5000 red cells). Sickness behaviour and core body temperature (digital rectal thermometer, *TmeElectronics*) were also recorded daily. To assess anaemia, erythrocytes in 2 μl blood (collected from the tail tip) were counted using a Z2 Coulter Particle count and size analyser (*Beckman Coulter*). We determined the healthy range in uninfected C57Bl/6J mice housed in our facility to be 8.8 - 10.5*10^9^ erythrocytes*ml^−1^. Anaemia was classified as severe when red cell loss exceeded 50 %.

Chronic infection was verified after 40-days of blood-stage parasitaemia by quantitative PCR of parasite 18S ribosomal DNA. DNA was extracted from 20 μl blood (collected from the tail tip) using Quick DNA Universal Kit (*Zymo Research*), and amplified using TaqMan Universal PCR Mastermix (*ThermoFisher*) with 9 μM of forward primer (5-GCGAGAAAGTTAAAAGAATTGA-3), 9 μM of reverse primer (5-CTAGTGAGTTTCCCCGTGTT-3) and 2.5 μM of probe ([6FAM] - AAATTAAGCCGCAAGCTCCACG - [TAM]) on a Roche Lightcycler 480 (40 cycles of amplification). A standard curve of red cells spiked with known numbers of parasites allowed accurate quantification; mice with > 5 parasites*μl ^−1^ (limit of detection) were considered chronically infected with *P. chabaudi*, and all other mice were excluded from the study. Chronic infection was cleared using 100 mg/kg chloroquine diphosphate salt (*Sigma*, dissolved in water) administered by oral gavage daily for 10-days. Memory responses were assessed 30-days after the start of chloroquine treatment, and at this time-point once-infected mice were compared to uninfected age-matched controls that received the same schedule of chloroquine treatment.

### Malaria reinfection model

Mice were first infected with *P. chabaudi* AS by intravenous (iv) injection of sporozoites, and those with qPCR-confirmed chronic parasitaemia were chloroquine treated on day 40 of blood-stage infection. Thirty days after the start of chloroquine treatment mice were infected for a second time but now by iv injection of 5 x 10^5^ mosquito-transmitted *P. chabaudi* AJ blood-stage parasites. Reinfection was initiated by this route to avoid confounding factors that may arise as a result of liver-stage immunity, which can be observed in C57Bl/6 mice after a single infection with *P. chabaudi* AS (Nahrendorf et al., 2015). Note that uninfected age-matched controls received the same schedule of chloroquine treatment as reinfected mice.

### Quantification of plasma proteins by ELISA

Platelet-depleted plasma was prepared from heparinised (*Wockhardt*) blood using two consecutive centrifugation steps (1000 xg for 10 minutes followed by 2000 xg for 15 minutes). Plasma was kept cold throughout and aliquots were stored at - 80°C. We used commercially available ELISA kits to quantify plasma IFNγ (mouse IFNγ platinum ELISA extra sensitive, *ebioscience*), CXCL10 (mouse IP-10 platinum ELISA, *ebioscience*) and Angiopoietin-2 (mouse/rat Angiopoietin-2 quantine ELISA kit, *R&D Systems*). Absorbance was measured using a Multiskan Ascent (*MTX Lab systems*) or FluoSTAR Omega (*BMG Labtech*) plate reader.

### Tissue preparation for histology

Spleens and femurs from *P. chabaudi* infected and once-infected mice (and uninfected age-matched controls) were fixed in 10 % neutral buffered Formalin (*Sigma*) for 24 or 48-hours, respectively. Bones were then decalcified for 48-hours using 10 % EDTA (pH 7.2) with gentle shaking at 55°C. After these steps, tissues were stored in 70 % Ethanol and photographed to visualise macroscopic changes resulting from malaria. The spleen and both femurs from each mouse were paraffin-embedded in a single block and 5 μm sections were prepared (cross section for spleen and longitudinal section for bone). Sections were stained with Hematoxylin and Eosin (H&E, *ThermoFisher*) or Prussian Blue and Neutral Red (*Scientific Laboratory Supplies* and *VWR*) at the Shared University Research Facilities, University of Edinburgh. Stained slides were assessed using a Nikon A400a bright-field microscope and images were taken with a Zeiss 503 high-resolution colour camera. Images were cropped, white balance was adjusted and the brightness/contrast standardised between samples using Adobe Photoshop CS6.

### Parasite sequestration

Mice were infected by intraperitoneal injection of 10^5^ mosquito-transmitted *P. chabaudi* blood-stage parasites and circulating parasitaemia was measured at the peak of schizogony (13:00 GMT) by Giemsa stained thin blood films (counting at least 5000 red cells). Immediately thereafter, mice were euthanised and exsanguinated - spleen and both femurs were prepared for histology as described above. In addition, the left lobe of the liver, the left lung and kidney, the duodenum and the heart were fixed in 10 % neutral buffered Formalin for 24-hours and then stored in 70 % Ethanol. All organs from each mouse were paraffin-embedded in a single block and 5 μm sections were prepared (longitudinal section for bone and cross section for all other organs) and H&E stained. The number of infected red cells contained within the vessels of each organ was quantified (counting at least 1000 red cells) and displayed relative to peripheral parasitaemia.

### Flow cytometry and cell sorting

Sodium heparin (*Wockhardt*) was used as anticoagulant for whole blood samples; spleens were dissociated in C-tubes using a gentleMACS Octo Dissociator (*Miltenyi Biotec*); and bone marrow was flushed from femurs using a 27½G needle/syringe loaded with IMDM. Single cell suspensions were filtered through a 70 μm cell strainer and after red cell lysis leukocytes were counted on a haemocytometer; up to 2 x 10^6^ cells per well were placed into a 96 well V bottom plate for staining. A Zombie Aqua Fixable Viability Dye (*Biolegend*) was used to identify dead cells, after which Fc receptors were blocked using TruStain FcX (anti-mouse CD16/32, *Biolegend*). Cell surface staining was performed at room temperature (for details of antibodies and panels refer to Additional File 1) and for ChIPseq experiments cells were subsequently fixed in PBS with 1 % paraformaldehyde (*Alpha Aesar*) and 10 % FBS for 10 minutes at room temperature (reaction was quenched with 125 mM Glycine (*Sigma*)). Note that across experiments the viability of leukocytes always exceeded 93.8 % (no viability stain for cell sorting), and to confirm the identity of patrolling monocytes we performed an intracellular stain for the transcription factor Nr4a1 (clone 12.14, *eBioscience*) using the FoxP3/Transcription Factor Buffer Staining Set (*eBioscience*).

Cells were acquired on an LSR Fortessa (*BD Biosciences*) or sorted using an Aria II (*BD Biosciences*, 85 μm nozzle, sort setting “purity”). Both cytometers used BD FACS Diva v8 software and data were subsequently analysed using FlowJo v9 (for details of gating strategies see Additional File 1). Note that CD115 (*Csf1ŕ*) was not used to identify spleen monocytes when sorting since engagement of the CSF1 receptor has been shown to induce transcriptional changes (Jung et al., 2000). Samples with a sort purity < 95 % were excluded from the study.

The absolute number of cells in each tissue was calculated from leukocyte counts of single cell suspensions. For bone marrow we estimated that one femur contains approximately 11 % of total mouse bone marrow (Colvin et al., 2004) and extrapolated accordingly. For whole blood we recorded the volume collected during exsanguination and then extrapolated to total circulating blood volume, according to body weight. This approach allows a direct comparison of cell numbers across tissues.

### Cytospin of red pulp macrophages

Red pulp macrophages (Lineage-F4/80+ B220- CD11b^int^ CD11c^int^ MHC-II+) were flow-sorted from the spleens of uninfected mice and collected into polypropylene tubes containing IMDM supplemented with 20 % FBS and 8 mM L-Glutamine. Sorted cells were then spun (1000 xg for 5 minutes) onto glass slides using a Shandon Cytospin 3 Cytocentrifuge (*ThermoScientific*) and stained with Prussian Blue and Neutral Red (*Sigma*). Red pulp macrophages were visualised and photographed using a Leica DM1000 light microscope (100x oil objective).

### *In vitro* stimulation of monocytes with LPS

30,000 inflammatory monocytes (Lineage-Ly6G- CD11b+ CD11c- Ly6C^hi^) were flow-sorted from the spleens of chronically infected mice (*P. chabaudi* AJ) or uninfected controls and collected into polypropylene tubes containing IMDM supplemented with 5 % FBS and 8 mM L-Glutamine. Following a gentle spin (450 xg for 10 minutes, slow brake) monocytes were resuspended in 90 μl pre-warmed IMDM containing 10 % FBS and 8 mM L-Glutamine, and transferred to an ultra-low attachment 96 well flat bottom cell culture plate (*Corning*). To stimulate cells 0.3 ng* μl^−1^ LPS (Lipopolysaccharide from Escherichia coli 0111:B4, *Sigma*) was added and cells were incubated for 4 hours at 37°C and 7 % CO2. RNA from both adherent and non-adherent cells was preserved in 1 ml TRIzol Reagent (*ThermoFisher*) for downstream steps.

### *Ex vivo* RNA sequencing of monocytes

10,000 inflammatory monocytes (Lineage-Ly6G- CD11b+ CD11c- Ly6C^hi^) were flow-sorted from the spleens of *P. chabaudi* infected mice, once-infected mice or uninfected controls and collected into 1.5 ml eppendorf tubes containing 1 ml TRIzol Reagent. Samples were inverted ten times, incubated at room temperature for 5 minutes and snap frozen on dry ice; all samples were stored at - 80°C prior to RNA extraction.

RNA was extracted using a modified phenol-chloroform protocol (Chomczynski and Sacchi, 2006) with 1-Bromo-3-chloropropane and Isopropanol (*Sigma* and *VWR*, respectively). Total RNA was quantified and assessed for quality and integrity by Bioanalyser (RNA Pico 6000 Chip, *Agilent*) - all sequenced samples had a RIN value above 8. cDNA was generated from 2 ng total RNA using the SMART-Seq v4 Ultra Low Input RNA Kit (*Clontech Laboratories*) and amplified using 11 cycles of PCR. Amplified cDNA was purified using Agencourt AMPure XP beads (*Beckman Coulter*), quantified on a Qubit 2.0 Fluorometer (dsDNA HS assay, *ThermoFisher*) and quality assessed by Bioanalyser (DNA HS Kit, *Agilent*). Libraries were then constructed from 150 pg of cDNA using the Nextera XT DNA Library Preparation Kit (*Illumina*) according to the manufacturer’s instructions. Libraries were quantified by Qubit (dsDNA HS assay) and fragment size distribution was assessed by Bioanalyser (DNA HS Kit). Using this information, samples were combined to create equimolar library pools that were sequenced on a NextSeq 550 platform (*Illumina*) to yield 75 bp paired-end (PE) reads; the median number of PE reads passing QC across all experiments was 4.79 x 10^7^.

### RNA sequencing analysis

FastQ files were downloaded from BaseSpace (*Illumina*) and raw sequence data assessed for quality and content using FastQC. We aligned paired-end sequences to the Ensembl release 96 murine transcripts set with bowtie2 v2.2.7 (parameters: —very-sensitive -p 30 —no-mixed —nodiscordant —no-unal) to obtain sorted, indexed bam files. Counts for each transcript were obtained using samtools idxstats and transcript counts were imported into the R/Bioconductor environment using the DESeq2 package (Love et al., 2014) for pairwise comparisons. Lists of differentially expressed transcripts were filtered to retain only those with an adjusted p value (p_adj_) < 0.01 and a fold change > 1.5 using R v3.6; multiple transcripts annotated to the same gene were consolidated by keeping the transcript with the highest absolute fold change. Heatmaps and stacked circular bar charts were generated using the R ggplot2 package (Wickham, 2016). All RNAseq data are publicly available (GEO accession number GSE150047).

### Functional gene enrichment analysis using ClueGO

Lists of differentially expressed genes were imported into ClueGO v2.5.4 (Bindea et al., 2009; Mlecnik et al., 2014). ClueGO identified the significantly enriched GO terms (GO Biological Process and GO Molecular Function) associated with these genes and placed them into a functionally organised non-redundant gene ontology network based on the following parameters: p_adj_ cutoff = 0.01; correction method used = Bonferroni step down; min. GO level = 5; max. GO level = 11; number of genes = 3; min. percentage = 5.0; GO fusion = true; sharing group percentage = 40.0; merge redundant groups with > 40.0% overlap; kappa score threshold = 0.4; and evidence codes used [All]. Each of the functional groups was assigned a unique colour and a network was then generated using an edge-weighted spring-embedded layout based on kappa score. We found that some GO terms were shared between multiple groups and so we manually merged these functionally connected groups to form supergroups, which we named according to the leading GO term (lowest p_adj_ with min. GO level 5).

### Chromatin immunoprecipitation for sequencing (ChIPseq)

50,000 fixed inflammatory monocytes (Lineage- Ly6G- cKit- CD135- CD11b+ Ly6C^hi^) were flow-sorted from the bone marrow of once-infected mice or uninfected controls and collected into polypropylene tubes containing IMDM supplemented with 5 % FBS and 8 mM L-Glutamine. Sorted monocytes were pelleted by centrifugation and washed in HBSS (*Gibco*) that was supplemented with protease inhibitors (complete ULTRA Tablets Protease Inhibitor Cocktail, *Roche*) and 5 mM sodium butyrate (*Alpha Aesar*); cell pellets were stored at - 80°C. Note that for each biological sample we pooled the femurs and tibias from two mice and three tubes (each containing 50,000 cells) were collected for every sample so that we could perform chromatin-immunoprecipitation (ChIP) with three different antibodies (H3K27ac, H3K4me1 and H3K9me3).

We performed ChIP using the True MicroChIP Kit (*Diagenode*). In brief, chromatin was sheared using a Bioruptor Pico Sonicator (5 cycles: 30 seconds ON - 30 seconds OFF, fragments 100 - 300 bp, *Diagenode*) and 10 % of sheared chromatin was kept as input control whilst 90 % was immunoprecipitated overnight using antibodies against H3K27ac, H3K4me1 or H3K9me3 (all ChIP-grade from *Diagenode*). Protein A coated magnetic beads were then added to samples and after a 6-hour incubation unbound chromatin fragments were removed by thorough washing. ChIP and input DNA were decrosslinked and purified using MicroChIP DiaPure columns (*Diagenode*).

Libraries were prepared using the MicroPlex v2 Library Preparation Kit (*Diagenode*) and amplification was monitored in real-time on a LightCycler 480 (*Roche*) to ensure the optimum number of cycles was used. Amplified libraries were quantified by Qubit (dsDNA HS assay) and fragment size distribution assessed by Bioanalyser (DNA HS Kit). After equimolar pooling of samples we purified libraries with AMPure XP beads and sequenced on a HiSeq 4000 (75 bp PE reads) or NovaSeq S1 (100 bp PE reads) (both *Illumina*). A step-by-step guide to our optimised ChIPseq protocol is available at <u>protocols.io</u>.

### ChIPseq analysis

ChIPseq data quality and content were assessed using FastQC; all samples passed initial QC and were aligned to the mm10 genome using bowtie2 v2.2.7 (parameters: —very-sensitive -p 30 —nomixed —no-discordant —no-unal). We then used the motif discovery software HOMER (v4.10 (Heinz et al., 2010)) to turn indexed bam files (generated using samtools idxstats) into tag directories of individual ChIP and input samples. Alignments were converted to bedgraph format using the HOMER script makeUCSCfile. Wig format outputs were converted to tdf files to view data in the Integrative Genomics Viewer (IGV v.2.7.2 (Thorvaldsdottir et al., 2013)) using igvtools v2.3.93 (parameters: toTDF -z 7 -f p98). In HOMER, we identified areas of the genome where ChIP read counts were significantly enriched over fragmented, non-immunoprecipitated input DNA (which indicates the presence of a histone mark) by calling peaks in ChIP relative to sample-matched input DNA using predefined parameters (H3K27ac and H3K9me3 used “regions” and H3K4me1 used “typical” and “supertypical”). Default settings were used in every case with the exception of fold change over input, which was set to > 3-fold for H3K9me3.

Individual samples were then combined in HOMER to create pooled ChIP tag directories. We selected only samples with a high IP efficiency (> 5 % for H3K27ac and H3K9me3, and > 10 % for H3K4me1) to generate pooled tag directories that comprised: 4 biological replicates for H3K27ac; 2 (once-infected) or 3 (uninfected) biological replicates for H3K4me1; and 1 replicate for H3K9me3. Bedgraphs of pooled ChIP tag directories were generated and converted to tdf format as above using more than 2.2 x 10^8^ tags for H3K27ac, 1.1 x 10^8^ tags for H3K4me1 and 7 x 10^7^ tags for H3K9me3. Peaks were called on pooled ChIP samples relative to pool-matched input DNA using the parameters described above. We created tdf files that indicated the position of peaks across the genome with a fixed height bar and visualised the histone modification profile alongside peak location in IGV. To ask whether genes were marked or not marked we looked for the presence or absence of peaks within 10 kb (H3K27ac and H3K9me3) or 100 kb (H3K4me1) of the transcription start site. To identify differentially modified regions (DMR) across the genome we again called peaks on pooled ChIP samples but this time instead of using non-immunoprecipitated input DNA to correct for background we called peaks in once-infected mice relative to uninfected controls (and vice versa). In this way, we identified areas of the genome where read counts were significantly enriched in one or the other experimental group. Low confidence peaks were removed by applying a peak score cut off > 3 and DMR were annotated to the nearest gene using the script annotatePeaks in HOMER. We then asked how many of the 2848 tolerised/specialised genes identified by RNAseq were annotated with a differentially modified region (if a gene was annotated with more than one DMR then the region with the highest peak score was retained). All ChIPseq data (inc. individual biological replicates and pooled tag directories) are publicly available (GEO accession number GSE150478).

### *Ex vivo* transcriptional profiling of red pulp macrophages

10,000 red pulp macrophages (Lineage- F4/80+ B220- CD11b^int^ CD11c^int^ autofluorescent cells) were flow-sorted from mice 100 days after self-resolving *P. chabaudi* infection or from uninfected age-matched controls. Sorted cells were collected into 1.5 ml eppendorf tubes containing 1 ml TRIzol Reagent (*ThermoFisher*) and samples were stored at - 80°C prior to RNA extraction. RNA was extracted using a modified phenol-chloroform protocol (Chomczynski and Sacchi, 2006) and treated with Baseline-ZERO DNase to remove genomic DNA (*Illumina*). DNase-treated RNA was then purified using the RNA Clean and Concentrator Kit (*Zymo Research*) and total RNA was quantified and assessed for quality and integrity by Bioanalyser (RNA Pico 6000 Chip, *Agilent*). RNA samples were processed for gene expression analysis using the GeneChip WT Pico Kit and Mouse Gene 1.0 ST Array (*Affymetrix*) according to the manufacturer’s instructions.

Microarray data were processed in R/Bioconductor making use of the oligo, pd.mta.1.0 and mta10sttranscriptcluster packages. Data quality was assessed using the arrayQualityMetrics package (Kauffmann and Huber, 2010); all samples passed QC and were normalised using robust multi-array average (RMA), which results in log2 expression intensities. Limma (linear models for microarray data) and eBayes packages were used for pairwise comparisons to find differentially expressed genes (DEG) between groups (AS vs uninfected, AJ vs uninfected and AJ vs AS) but yielded zero DEG in all comparisons (p_adj_ < 0.05). Log2 expression intensities of signature genes for monocytes, tissue resident macrophages and dendritic cells were plotted using the heatmap.2() function in R; these genelists were manually compiled from published studies that set out to identify the gene expression profiles that underpin identity in myeloid cells (Gautier et al., 2012; Haldar et al., 2014; Miller et al., 2012; Okabe and Medzhitov, 2014). Microarray data are publicly available (GEO accession number GSE149894).

### Data access and in-depth protocols

Step-by-step protocols of key experimental procedures are available at protocols.io. All RNAseq, ChIPseq and microarray data have been deposited in NCBI’s Gene Expression Omnibus (Edgar et al., 2002) and are accessible through GEO SuperSeries accession number GSE150479 (https://www.ncbi.nlm.nih.gov/geo/query/acc.cgi?acc=GSE150479).

## References

Ademolue, T.W., Aniweh, Y., Kusi, K.A., and Awandare, G.A. (2017). Patterns of inflammatory responses and parasite tolerance vary with malaria transmission intensity. Malaria journal 16, 145.

Aegerter, H., Kulikauskaite, J., Crotta, S., Patel, H., Kelly, G., Hessel, E.M., Mack, M., Beinke, S., and Wack, A. (2020). Influenza-induced monocyte-derived alveolar macrophages confer prolonged antibacterial protection. Nature immunology 21, 145–157.

Akilesh, H.M., Buechler, M.B., Duggan, J.M., Hahn, W.O., Matta, B., Sun, X., Gessay, G., Whalen, E., Mason, M., Presnell, S.R., et al. (2019). Chronic TLR7 and TLR9 signaling drives anemia via differentiation of specialized hemophagocytes. Science 363.

Belyaev, N.N., Brown, D.E., Diaz, A.I., Rae, A., Jarra, W., Thompson, J., Langhorne, J., and Potocnik, A.J. (2010). Induction of an IL7-R(+)c-Kit(hi) myelolymphoid progenitor critically dependent on IFN-gamma signaling during acute malaria. Nature immunology 11, 477–485.

Bindea, G., Mlecnik, B., Hackl, H., Charoentong, P., Tosolini, M., Kirilovsky, A., Fridman, W.H., Pages, F., Trajanoski, Z., and Galon, J. (2009). ClueGO: a Cytoscape plug-in to decipher functionally grouped gene ontology and pathway annotation networks. Bioinformatics 25, 1091–1093.

Bleriot, C., Dupuis, T., Jouvion, G., Eberl, G., Disson, O., and Lecuit, M. (2015). Liver-resident macrophage necroptosis orchestrates type 1 microbicidal inflammation and type-2-mediated tissue repair during bacterial infection. Immunity 42, 145–158.

Bonnardel, J., T’Jonck, W., Gaublomme, D., Browaeys, R., Scott, C.L., Martens, L., Vanneste, B., De Prijck, S., Nedospasov, S.A., Kremer, A., et al. (2019). Stellate Cells, Hepatocytes, and Endothelial Cells Imprint the Kupffer Cell Identity on Monocytes Colonizing the Liver Macrophage Niche. Immunity 51, 638–654 e639.

Brugat, T., Cunningham, D., Sodenkamp, J., Coomes, S., Wilson, M., Spence, P.J., Jarra, W., Thompson, J., Scudamore, C., and Langhorne, J. (2014). Sequestration and histopathology in Plasmodium chabaudi malaria are influenced by the immune response in an organ-specific manner. Cellular microbiology 16, 687–700.

Buffet, P.A., Safeukui, I., Deplaine, G., Brousse, V., Prendki, V., Thellier, M., Turner, G.D., and Mercereau-Puijalon, O. (2011). The pathogenesis of Plasmodium falciparum malaria in humans: insights from splenic physiology. Blood 117, 381–392.

Cambos, M., and Scorza, T. (2011). Robust erythrophagocytosis leads to macrophage apoptosis via a hemin-mediated redox imbalance: role in hemolytic disorders. Journal of leukocyte biology 89, 159–171.

Carlin, L.M., Stamatiades, E.G., Auffray, C., Hanna, R.N., Glover, L., Vizcay-Barrena, G., Hedrick, C.C., Cook, H.T., Diebold, S., and Geissmann, F. (2013). Nr4a1-dependent Ly6C(low) monocytes monitor endothelial cells and orchestrate their disposal. Cell 153, 362–375.

Cheng, S.C., Quintin, J., Cramer, R.A., Shepardson, K.M., Saeed, S., Kumar, V., Giamarellos-Bourboulis, E.J., Martens, J.H., Rao, N.A., Aghajanirefah, A., et al. (2014). mTOR- and HIF-1alpha-mediated aerobic glycolysis as metabolic basis for trained immunity. Science 345, 1250684.

Chomczynski, P., and Sacchi, N. (2006). The single-step method of RNA isolation by acid guanidinium thiocyanate-phenol-chloroform extraction: twenty-something years on. Nature protocols 1, 581–585.

Colvin, G.A., Lambert, J.F., Abedi, M., Hsieh, C.C., Carlson, J.E., Stewart, F.M., and Quesenberry, P.J. (2004). Murine marrow cellularity and the concept of stem cell competition: geographic and quantitative determinants in stem cell biology. Leukemia 18, 575–583.

Cumnock, K., Gupta, A.S., Lissner, M., Chevee, V., Davis, N.M., and Schneider, D.S. (2018). Host Energy Source Is Important for Disease Tolerance to Malaria. Curr Biol 28, 1635–1642 e1633.

Czopek, A., Moorhouse, R., Guyonnet, L., Farrah, T., Lenoir, O., Owen, E., van Bragt, J., Costello, H.M., Menolascina, F., Baudrie, V., et al. (2019). A novel role for myeloid endothelin-B receptors in hypertension. Eur Heart J 40, 768–784.

Dela Cruz, C.S., Liu, W., He, C.H., Jacoby, A., Gornitzky, A., Ma, B., Flavell, R., Lee, C.G., and Elias, J.A. (2012). Chitinase 3-like-1 promotes Streptococcus pneumoniae killing and augments host tolerance to lung antibacterial responses. Cell host & microbe 12, 34–46.

Dominguez-Andres, J., and Netea, M.G. (2018). Long-term reprogramming of the innate immune system. Journal of leukocyte biology.

Edgar, R., Domrachev, M., and Lash, A.E. (2002). Gene Expression Omnibus: NCBI gene expression and hybridization array data repository. Nucleic acids research 30, 207–210.

Felger, I., Maire, M., Bretscher, M.T., Falk, N., Tiaden, A., Sama, W., Beck, H.P., Owusu-Agyei, S., and Smith, T.A. (2012). The dynamics of natural Plasmodium falciparum infections. PloS one 7, e45542.

Ferreira, A., Balla, J., Jeney, V., Balla, G., and Soares, M.P. (2008). A central role for free heme in the pathogenesis of severe malaria: the missing link? J Mol Med (Berl) 86, 1097–1111.

Foster, S.L., Hargreaves, D.C., and Medzhitov, R. (2007). Gene-specific control of inflammation by TLR-induced chromatin modifications. Nature 447, 972–978.

Gabrilovich, D.I. (2017). Myeloid-Derived Suppressor Cells. Cancer Immunol Res 5, 3–8.

Gautier, E.L., Shay, T., Miller, J., Greter, M., Jakubzick, C., Ivanov, S., Helft, J., Chow, A., Elpek, K.G., Gordonov, S., et al. (2012). Gene-expression profiles and transcriptional regulatory pathways that underlie the identity and diversity of mouse tissue macrophages. Nature immunology 13, 1118–1128.

Goncalves, B.P., Huang, C.Y., Morrison, R., Holte, S., Kabyemela, E., Prevots, D.R., Fried, M., and Duffy, P.E. (2014). Parasite burden and severity of malaria in Tanzanian children. The New England journal of medicine 370, 1799–1808.

Gozzelino, R., Andrade, B.B., Larsen, R., Luz, N.F., Vanoaica, L., Seixas, E., Coutinho, A., Cardoso, S., Rebelo, S., Poli, M., et al. (2012). Metabolic adaptation to tissue iron overload confers tolerance to malaria. Cell host & microbe 12, 693–704.

Guilliams, M., Mildner, A., and Yona, S. (2018). Developmental and Functional Heterogeneity of Monocytes. Immunity 49, 595–613.

Guilliams, M., Thierry, G.R., Bonnardel, J., and Bajenoff, M. (2020). Establishment and Maintenance of the Macrophage Niche. Immunity 52, 434–451.

Gupta, S., Snow, R.W., Donnelly, C.A., Marsh, K., and Newbold, C. (1999). Immunity to non-cerebral severe malaria is acquired after one or two infections. Nature medicine 5, 340–343.

Haldar, M., Kohyama, M., So, A.Y., Kc, W., Wu, X., Briseno, C.G., Satpathy, A.T., Kretzer, N.M., Arase, H., Rajasekaran, N.S., et al. (2014). Heme-mediated SPI-C induction promotes monocyte differentiation into iron-recycling macrophages. Cell 156, 1223–1234.

Heinz, S., Benner, C., Spann, N., Bertolino, E., Lin, Y.C., Laslo, P., Cheng, J.X., Murre, C., Singh, H., and Glass, C.K. (2010). Simple combinations of lineage-determining transcription factors prime cis-regulatory elements required for macrophage and B cell identities. Mol Cell 38, 576–589.

Jakeman, G.N., Saul, A., Hogarth, W.L., and Collins, W.E. (1999). Anaemia of acute malaria infections in non-immune patients primarily results from destruction of uninfected erythrocytes. Parasitology 119 (Pt 2), 127–133.

Jeney, V., Ramos, S., Bergman, M.L., Bechmann, I., Tischer, J., Ferreira, A., Oliveira-Marques, V., Janse, C.J., Rebelo, S., Cardoso, S., et al. (2014). Control of disease tolerance to malaria by nitric oxide and carbon monoxide. Cell reports 8, 126–136.

Jung, K., Heishi, T., Khan, O.F., Kowalski, P.S., Incio, J., Rahbari, N.N., Chung, E., Clark, J.W., Willett, C.G., Luster, A.D., et al. (2017). Ly6Clo monocytes drive immunosuppression and confer resistance to anti-VEGFR2 cancer therapy. The Journal of clinical investigation 127, 3039–3051.

Jung, S., Aliberti, J., Graemmel, P., Sunshine, M.J., Kreutzberg, G.W., Sher, A., and Littman, D.R. (2000). Analysis of fractalkine receptor CX(3)CR1 function by targeted deletion and green fluorescent protein reporter gene insertion. Molecular and cellular biology 20, 4106–4114.

Kauffmann, A., and Huber, W. (2010). Microarray data quality control improves the detection of differentially expressed genes. Genomics 95, 138–142.

Kaufmann, E., Sanz, J., Dunn, J.L., Khan, N., Mendonca, L.E., Pacis, A., Tzelepis, F., Pernet, E., Dumaine, A., Grenier, J.C., et al. (2018). BCG Educates Hematopoietic Stem Cells to Generate Protective Innate Immunity against Tuberculosis. Cell 172, 176–190 e119.

Kohyama, M., Ise, W., Edelson, B.T., Wilker, P.R., Hildner, K., Mejia, C., Frazier, W.A., Murphy, T.L., and Murphy, K.M. (2009). Role for Spi-C in the development of red pulp macrophages and splenic iron homeostasis. Nature 457, 318–321.

Koliaraki, V., Prados, A., Armaka, M., and Kollias, G. (2020). The mesenchymal context in inflammation, immunity and cancer. Nature immunology.

Lai, S.M., Sheng, J., Gupta, P., Renia, L., Duan, K., Zolezzi, F., Karjalainen, K., Newell, E.W., and Ruedl, C. (2018). Organ-Specific Fate, Recruitment, and Refilling Dynamics of Tissue-Resident Macrophages during Blood-Stage Malaria. Cell reports 25, 3099–3109 e3093.

Love, M.I., Huber, W., and Anders, S. (2014). Moderated estimation of fold change and dispersion for RNA-seq data with DESeq2. Genome Biol 15, 550.

Ma, Z., Wang, H., Cai, Y., Wang, H., Niu, K., Wu, X., Ma, H., Yang, Y., Tong, W., Liu, F., et al. (2018). Epigenetic drift of H3K27me3 in aging links glycolysis to healthy longevity in Drosophila. eLife 7.

Machiels, B., Dourcy, M., Xiao, X., Javaux, J., Mesnil, C., Sabatel, C., Desmecht, D., Lallemand, F., Martinive, P., Hammad, H., et al. (2017). A gammaherpesvirus provides protection against allergic asthma by inducing the replacement of resident alveolar macrophages with regulatory monocytes. Nature immunology 18, 1310–1320.

Mandala, W.L., Msefula, C.L., Gondwe, E.N., Drayson, M.T., Molyneux, M.E., and MacLennan, C.A. (2017). Cytokine Profiles in Malawian Children Presenting with Uncomplicated Malaria, Severe Malarial Anemia, and Cerebral Malaria. Clinical and vaccine immunology : CVI 24.

Marsh, K., and Kinyanjui, S. (2006). Immune effector mechanisms in malaria. Parasite immunology 28, 51–60.

Marsh, K., and Snow, R.W. (1999). Malaria transmission and morbidity. Parassitologia 41, 241–246.

Martins, R., Carlos, A.R., Braza, F., Thompson, J.A., Bastos-Amador, P., Ramos, S., and Soares, M.P. (2019). Disease Tolerance as an Inherent Component of Immunity. Annual review of immunology.

Medzhitov, R., Schneider, D.S., and Soares, M.P. (2012). Disease tolerance as a defense strategy. Science 335, 936–941.

Menezes, S., Melandri, D., Anselmi, G., Perchet, T., Loschko, J., Dubrot, J., Patel, R., Gautier, E.L., Hugues, S., Longhi, M.P., et al. (2016). The Heterogeneity of Ly6Chi Monocytes Controls Their Differentiation into iNOS+ Macrophages or Monocyte-Derived Dendritic Cells. Immunity 45, 1205–1218.

Mildner, A., Schonheit, J., Giladi, A., David, E., Lara-Astiaso, D., Lorenzo-Vivas, E., Paul, F., Chappell-Maor, L., Priller, J., Leutz, A., et al. (2017). Genomic Characterization of Murine Monocytes Reveals C/EBPbeta Transcription Factor Dependence of Ly6C-Cells. Immunity 46, 849–862 e847.

Miller, J.C., Brown, B.D., Shay, T., Gautier, E.L., Jojic, V., Cohain, A., Pandey, G., Leboeuf, M., Elpek, K.G., Helft, J., et al. (2012). Deciphering the transcriptional network of the dendritic cell lineage. Nature immunology 13, 888–899.

Mitroulis, I., Kalafati, L., Hajishengallis, G., and Chavakis, T. (2018). Myelopoiesis in the Context of Innate Immunity. J Innate Immun 10, 365–372.

Mlecnik, B., Bindea, G., Angell, H.K., Sasso, M.S., Obenauf, A.C., Fredriksen, T., Lafontaine, L., Bilocq, A.M., Kirilovsky, A., Tosolini, M., et al. (2014). Functional network pipeline reveals genetic determinants associated with in situ lymphocyte proliferation and survival of cancer patients. Science translational medicine 6, 228ra237.

Nahrendorf, W., Spence, P.J., Tumwine, I., Levy, P., Jarra, W., Sauerwein, R.W., and Langhorne, J. (2015). Blood-stage immunity to Plasmodium chabaudi malaria following chemoprophylaxis and sporozoite immunization. eLife 4.

Netea, M.G., Joosten, L.A., Latz, E., Mills, K.H., Natoli, G., Stunnenberg, H.G., O’Neill, L.A., and Xavier, R.J. (2016). Trained immunity: A program of innate immune memory in health and disease. Science 352, aaf1098.

O’Neill, L.A., Kishton, R.J., and Rathmell, J. (2016). A guide to immunometabolism for immunologists. Nature reviews Immunology.

Okabe, Y., and Medzhitov, R. (2014). Tissue-specific signals control reversible program of localization and functional polarization of macrophages. Cell 157, 832–844.

Orlando, D.A., Chen, M.W., Brown, V.E., Solanki, S., Choi, Y.J., Olson, E.R., Fritz, C.C., Bradner, J.E., and Guenther, M.G. (2014). Quantitative ChIP-Seq normalization reveals global modulation of the epigenome. Cell reports 9, 1163–1170.

Otto, T.D., Bohme, U., Jackson, A.P., Hunt, M., Franke-Fayard, B., Hoeijmakers, W.A., Religa, A.A., Robertson, L., Sanders, M., Ogun, S.A., et al. (2014). A comprehensive evaluation of rodent malaria parasite genomes and gene expression. BMC biology 12, 86.

Palmieri, F. (2013). The mitochondrial transporter family SLC25: identification, properties and physiopathology. Mol Aspects Med 34, 465–484.

Pathak, V.A., and Ghosh, K. (2016). Erythropoiesis in Malaria Infections and Factors Modifying the Erythropoietic Response. Anemia 2016, 9310905.

Portugal, S., Moebius, J., Skinner, J., Doumbo, S., Doumtabe, D., Kone, Y., Dia, S., Kanakabandi, K., Sturdevant, D.E., Virtaneva, K., et al. (2014). Exposure-dependent control of malaria-induced inflammation in children. PLoS pathogens 10, e1004079.

Ramos, S., Carlos, A.R., Sundaram, B., Jeney, V., Ribeiro, A., Gozzelino, R., Bank, C., Gjini, E., Braza, F., Martins, R., et al. (2019). Renal control of disease tolerance to malaria. Proceedings of the National Academy of Sciences of the United States of America.

Schofield, L., and Grau, G.E. (2005). Immunological processes in malaria pathogenesis. Nature reviews Immunology 5, 722–735.

Schwarzer, E., Alessio, M., Ulliers, D., and Arese, P. (1998). Phagocytosis of the malarial pigment, hemozoin, impairs expression of major histocompatibility complex class II antigen, CD54, and CD11c in human monocytes. Infection and immunity 66, 1601–1606.

Schwarzer, E., Turrini, F., Ulliers, D., Giribaldi, G., Ginsburg, H., and Arese, P. (1992). Impairment of macrophage functions after ingestion of Plasmodium falciparum-infected erythrocytes or isolated malarial pigment. The Journal of experimental medicine 176, 1033–1041.

Scott, C.L., Zheng, F., De Baetselier, P., Martens, L., Saeys, Y., De Prijck, S., Lippens, S., Abels, C., Schoonooghe, S., Raes, G., et al. (2016). Bone marrow-derived monocytes give rise to self-renewing and fully differentiated Kupffer cells. Nature communications 7, 10321.

Seeley, J.J., and Ghosh, S. (2017). Molecular mechanisms of innate memory and tolerance to LPS. Journal of leukocyte biology 101, 107–119.

Seixas, E., Gozzelino, R., Chora, A., Ferreira, A., Silva, G., Larsen, R., Rebelo, S., Penido, C., Smith, N.R., Coutinho, A., et al. (2009). Heme oxygenase-1 affords protection against noncerebral forms of severe malaria. Proceedings of the National Academy of Sciences of the United States of America 106, 15837–15842.

Simpson, I.A., Dwyer, D., Malide, D., Moley, K.H., Travis, A., and Vannucci, S.J. (2008). The facilitative glucose transporter GLUT3: 20 years of distinction. American journal of physiology Endocrinology and metabolism 295, E242–253.

Spaulding, E., Fooksman, D., Moore, J.M., Saidi, A., Feintuch, C.M., Reizis, B., Chorro, L., Daily, J., and Lauvau, G. (2016). STING-Licensed Macrophages Prime Type I IFN Production by Plasmacytoid Dendritic Cells in the Bone Marrow during Severe Plasmodium yoelii Malaria. PLoS pathogens 12, e1005975.

Spence, P.J., Jarra, W., Levy, P., Nahrendorf, W., and Langhorne, J. (2012). Mosquito transmission of the rodent malaria parasite Plasmodium chabaudi. Malaria journal 11, 407.

Spence, P.J., Jarra, W., Levy, P., Reid, A.J., Chappell, L., Brugat, T., Sanders, M., Berriman, M., and Langhorne, J. (2013). Vector transmission regulates immune control of Plasmodium virulence. Nature 498, 228–231.

Theurl, I., Hilgendorf, I., Nairz, M., Tymoszuk, P., Haschka, D., Asshoff, M., He, S., Gerhardt, L.M., Holderried, T.A., Seifert, M., et al. (2016). On-demand erythrocyte disposal and iron recycling requires transient macrophages in the liver. Nature medicine 22, 945–951.

Thorvaldsdottir, H., Robinson, J.T., and Mesirov, J.P. (2013). Integrative Genomics Viewer (IGV): high-performance genomics data visualization and exploration. Brief Bioinform 14, 178–192.

Ulas, T., Pirr, S., Fehlhaber, B., Bickes, M.S., Loof, T.G., Vogl, T., Mellinger, L., Heinemann, A.S., Burgmann, J., Schoning, J., et al. (2017). S100-alarmin-induced innate immune programming protects newborn infants from sepsis. Nature immunology 18, 622–632.

von Seidlein, L., Olaosebikan, R., Hendriksen, I.C., Lee, S.J., Adedoyin, O.T., Agbenyega, T., Nguah, S.B., Bojang, K., Deen, J.L., Evans, J., et al. (2012). Predicting the clinical outcome of severe falciparum malaria in african children: findings from a large randomized trial. Clinical infectious diseases : an official publication of the Infectious Diseases Society of America 54, 1080–1090.

Weis, S., Carlos, A.R., Moita, M.R., Singh, S., Blankenhaus, B., Cardoso, S., Larsen, R., Rebelo, S., Schauble, S., Del Barrio, L., et al. (2017). Metabolic Adaptation Establishes Disease Tolerance to Sepsis. Cell 169, 1263–1275 e1214.

Weiss, D.J., Lucas, T.C.D., Nguyen, M., Nandi, A.K., Bisanzio, D., Battle, K.E., Cameron, E., Twohig, K.A., Pfeffer, D.A., Rozier, J.A., et al. (2019). Mapping the global prevalence, incidence, and mortality of Plasmodium falciparum, 2000-17: a spatial and temporal modelling study. Lancet.

Wendeln, A.C., Degenhardt, K., Kaurani, L., Gertig, M., Ulas, T., Jain, G., Wagner, J., Hasler, L.M., Wild, K., Skodras, A., et al. (2018). Innate immune memory in the brain shapes neurological disease hallmarks. Nature 556, 332–338.

Wickham, H. (2016). ggplot2: Elegant Graphics for Data Analysis (Springer-Verlag New York).

Yao, H., Arunachalam, G., Hwang, J.W., Chung, S., Sundar, I.K., Kinnula, V.L., Crapo, J.D., and Rahman, I. (2010). Extracellular superoxide dismutase protects against pulmonary emphysema by attenuating oxidative fragmentation of ECM. Proceedings of the National Academy of Sciences of the United States of America 107, 15571–15576.

Yeo, T.W., Lampah, D.A., Gitawati, R., Tjitra, E., Kenangalem, E., Piera, K., Price, R.N., Duffull, S.B., Celermajer, D.S., and Anstey, N.M. (2008). Angiopoietin-2 is associated with decreased endothelial nitric oxide and poor clinical outcome in severe falciparum malaria. Proceedings of the National Academy of Sciences of the United States of America 105, 17097–17102.

