## Supplementary material for "Inducible mechanisms of disease tolerance provide an alternative strategy of acquired immunity to malaria": Table_S1

## H3K27ac

| | Gene.Name | Chr | Start | End | Peak.Score | Focus.Ratio.Region.Size | Detailed.Annotation | Gene.Type | Distance.to.TSS | Nearest.PromoterID | Entrez.ID | Nearest.Unigene |
|---|---|---|---|---|---|---|---|---|---|---|---|---|
| 1 | Zdhhc21 | 4 | 82866346 | 82866846 | 7.25 | 500 | Intergenic | protein_coding | -6638 | ENSMUST00000107239 | ENSMUSG00000028403 | ENSMUST00000139401 |
| 2 | Abtb2 | 2 | 103603582 | 103604082 | 7.1 | 500 | protein_coding-intron (ENSMUST00000076212, intron 1 of 16) | protein_coding | 37522 | ENSMUST00000076212 | ENSMUSG00000032724 | ENSMUST00000076212 |
| 3 | Vegfa | 17 | 46012913 | 46013539 | 6.82 | 626 | Intergenic | protein_coding | 8146 | ENSMUST00000143985 | ENSMUSG00000023951 | ENSMUST00000150327 |
| 4 | Pola1 | X | 93416292 | 93417239 | 6.67 | 947 | protein_coding-intron (ENSMUST00000006856, intron 35 of 36) | protein_coding | 74181 | ENSMUST00000142479 | ENSMUSG00000006678 | ENSMUST00000006856 |
| 5 | Csl | 10 | 99761549 | 99762369 | 6.1 | 820 | lncRNA-exon (ENSMUST00000217945, exon 2 of 5) | protein_coding | -2301 | ENSMUST00000056085 | ENSMUSG00000046934 | ENSMUST00000056085 |
| 6 | Neto2 | 8 | 85694382 | 85695009 | 6.04 | 627 | protein_coding-intron (ENSMUST00000209479, intron 1 of 8) | protein_coding | -3686 | ENSMUST00000209259 | ENSMUSG00000036902 | ENSMUST00000209479 |
| 7 | Flot1 | 17 | 35824471 | 35824971 | 6.01 | 500 | protein_coding-TTS (ENSMUST00000174227) | protein_coding | 725 | ENSMUST00000173337 | ENSMUSG00000059714 | ENSMUST00000174297 |
| 8 | Ldlr | 9 | 21732865 | 21733499 | 5.86 | 634 | protein_coding-intron (ENSMUST00000034713, intron 3 of 17) | protein_coding | -2925 | ENSMUST00000217613 | ENSMUSG00000032193 | ENSMUST00000215917 |
| 9 | Pprc1 | 19 | 46070209 | 46070860 | 5.84 | 651 | protein_coding-intron (ENSMUST00000062322, intron 9 of 13) | protein_coding | -1220 | ENSMUST00000146882 | ENSMUSG00000055491 | ENSMUST00000099392 |
| 10 | Epha7 | 4 | 28770147 | 28770774 | 5.83 | 627 | Intergenic | protein_coding | -42671 | ENSMUST00000029964 | ENSMUSG00000028289 | ENSMUST00000080934 |
| 11 | Plcb2 | 2 | 118729876 | 118730376 | 5.67 | 500 | Intergenic | protein_coding | -1688 | ENSMUST00000102524 | ENSMUSG00000040061 | ENSMUST00000006415 |
| 12 | Ube2f | 1 | 91252511 | 91253011 | 5.67 | 500 | protein_coding-intron (ENSMUST00000059743, intron 1 of 10) | protein_coding | 2105 | ENSMUST00000191533 | ENSMUSG00000034343 | ENSMUST00000187494 |
| 13 | Rasal3 | 17 | 32395232 | 32395944 | 5.52 | 712 | protein_coding-promoter-TSS (ENSMUST00000143808) | protein_coding | -217 | ENSMUST00000143808 | ENSMUSG00000052142 | ENSMUST00000135968 |
| 14 | Ptch2 | 4 | 117106680 | 117107306 | 5.5 | 626 | protein_coding-TTS (ENSMUST00000156989) | protein_coding | 1755 | ENSMUST00000137209 | ENSMUSG00000028681 | ENSMUST00000030443 |
| 15 | Cd81 | 7 | 143043476 | 143043976 | 5.48 | 500 | lncRNA-intron (ENSMUST00000131129, intron 2 of 4) | protein_coding | -9013 | ENSMUST00000037941 | ENSMUSG00000037706 | ENSMUST00000208278 |
| 16 | B230118H07Rik | 2 | 101613445 | 101614071 | 5.45 | 626 | protein_coding-intron (ENSMUST00000090513, intron 1 of 6) | protein_coding | -3202 | ENSMUST00000136601 | ENSMUSG00000027165 | ENSMUST00000132640 |
| 17 | Trpm4 | 7 | 45300223 | 45300723 | 5.44 | 500 | Intergenic | protein_coding | 4731 | ENSMUST00000210639 | ENSMUSG00000038260 | ENSMUST00000042194 |
| 18 | Pcyox1l | 18 | 61710828 | 61711454 | 5.38 | 626 | Intergenic | protein_coding | -3506 | ENSMUST00000154763 | ENSMUSG00000024579 | ENSMUST00000195229 |
| 19 | Dapk2 | 9 | 66230602 | 66231102 | 5.27 | 500 | protein_coding-intron (ENSMUST00000034944, intron 3 of 11) | protein_coding | 10105 | ENSMUST00000132987 | ENSMUSG00000032380 | ENSMUST00000129442 |
| 20 | Ankib1 | 5 | 3712077 | 3712741 | 5.2 | 664 | protein_coding-intron (ENSMUST00000043551, intron 11 of 19) | protein_coding | -10277 | ENSMUST00000199043 | ENSMUSG00000040351 | ENSMUST00000200052 |
| 21 | Pnpt1 | 11 | 29117103 | 29117750 | 5.2 | 647 | Intergenic | protein_coding | -13318 | ENSMUST00000154924 | ENSMUSG00000020464 | ENSMUST00000148173 |
| 22 | Cenpe | 3 | 135213664 | 135214164 | 5.2 | 500 | protein_coding-intron (ENSMUST00000062893, intron 1 of 45) | protein_coding | 1377 | ENSMUST00000062893 | ENSMUSG00000045328 | ENSMUST00000199497 |
| 23 | Dnase1l3 | 14 | 7982115 | 7982743 | 5.18 | 628 | protein_coding-intron (ENSMUST00000026315, intron 4 of 7) | protein_coding | -5207 | ENSMUST00000145234 | ENSMUSG00000025279 | ENSMUST00000145234 |
| 24 | Adam3 | 8 | 24686528 | 24687231 | 5.17 | 703 | protein_coding-intron (ENSMUST00000033958, intron 18 of 21) | protein_coding | 1084 | ENSMUST00000167431 | ENSMUSG00000031553 | ENSMUST00000166986 |
| 25 | Eif2s2 | 2 | 154868784 | 154869423 | 5.17 | 639 | protein_coding-intron (ENSMUST00000137333, intron 1 of 4) | protein_coding | 23679 | ENSMUST00000166171 | ENSMUSG00000074656 | ENSMUST00000099173 |
| 26 | Cp | 3 | 19967887 | 19968387 | 5.15 | 500 | protein_coding-exon (ENSMUST00000003714, exon 4 of 20) | protein_coding | -1705 | ENSMUST00000150264 | ENSMUSG00000003617 | ENSMUST00000129135 |
| 27 | Myoz1 | 14 | 20647993 | 20648493 | 5.1 | 500 | protein_coding-TTS (ENSMUST00000090469) | protein_coding | 2475 | ENSMUST00000224472 | ENSMUSG00000068697 | ENSMUST00000225231 |
| 28 | Cux2 | 5 | 121988617 | 121989326 | 4.97 | 709 | protein_coding-intron (ENSMUST00000086317, intron 2 of 22) | protein_coding | 13373 | ENSMUST00000134326 | ENSMUSG00000042589 | ENSMUST00000197179 |
| 29 | Fmnl3 | 15 | 99370582 | 99371082 | 4.92 | 500 | protein_coding-promoter-TSS (ENSMUST00000081224) | protein_coding | -350 | ENSMUST00000088233 | ENSMUSG00000023008 | ENSMUST00000088233 |
| 30 | Zdhhc2 | 8 | 40459368 | 40460006 | 4.88 | 638 | protein_coding-intron (ENSMUST00000049389, intron 6 of 12) | protein_coding | 35583 | ENSMUST00000168799 | ENSMUSG00000039470 | ENSMUST00000128166 |
| 31 | Tada2a | 11 | 84098616 | 84099116 | 4.82 | 500 | protein_coding-intron (ENSMUST00000018795, intron 8 of 15) | protein_coding | -5012 | ENSMUST00000141852 | ENSMUSG00000018651 | ENSMUST00000134994 |
| 32 | Terf1 | 1 | 15818573 | 15819223 | 4.81 | 650 | protein_coding-intron (ENSMUST00000027057, intron 3 of 8) | protein_coding | 11257 | ENSMUST00000188684 | ENSMUSG00000025925 | ENSMUST00000188684 |
| 33 | Tnip3 | 6 | 65533129 | 65533629 | 4.81 | 500 | protein_coding-intron (ENSMUST00000212375, intron 1 of 10) | protein_coding | 7994 | ENSMUST00000212375 | ENSMUSG00000044162 | ENSMUST00000204241 |
| 34 | Rttn | 18 | 89109756 | 89110398 | 4.79 | 642 | protein_coding-intron (ENSMUST00000023828, intron 42 of 48) | protein_coding | 1970 | ENSMUST00000237196 | ENSMUSG00000023066 | ENSMUST00000236313 |
| 35 | Uchl5 | 1 | 143797873 | 143798500 | 4.77 | 627 | protein_coding-intron (ENSMUST00000018333, intron 7 of 10) | protein_coding | 3469 | ENSMUST00000185539 | ENSMUSG00000018189 | ENSMUST00000185742 |
| 36 | Evl | 12 | 108627950 | 108628450 | 4.71 | 500 | protein_coding-intron (ENSMUST00000021689, intron 1 of 13) | protein_coding | -7510 | ENSMUST00000223109 | ENSMUSG00000021262 | ENSMUST00000223109 |
| 37 | BC049762 | 11 | 51247835 | 51248335 | 4.65 | 500 | Intergenic | protein_coding | 8179 | ENSMUST00000231433 | ENSMUSG00000045942 | ENSMUST00000054226 |
| 38 | Trim37 | 11 | 87213626 | 87214126 | 4.65 | 500 | protein_coding-intron (ENSMUST00000041282, intron 22 of 23) | protein_coding | 23802 | ENSMUST00000154138 | ENSMUSG00000018548 | ENSMUST00000041282 |
| 39 | Esr1 | 10 | 4983449 | 4983949 | 4.63 | 500 | protein_coding-intron (ENSMUST00000067086, intron 7 of 8) | protein_coding | 47173 | ENSMUST00000137012 | ENSMUSG00000019768 | ENSMUST00000105590 |
| 40 | Cdk5rap2 | 4 | 70319189 | 70319826 | 4.58 | 637 | protein_coding-intron (ENSMUST00000076541, intron 9 of 33) | protein_coding | -54827 | ENSMUST00000138561 | ENSMUSG00000039298 | ENSMUST00000126416 |
| 41 | Naa10 | X | 73918003 | 73918503 | 4.58 | 500 | protein_coding-exon (ENSMUST00000143076, exon 5 of 5) | protein_coding | 1609 | ENSMUST00000139105 | ENSMUSG00000031388 | ENSMUST00000096316 |
| 42 | Nampt | 12 | 32833758 | 32834258 | 4.58 | 500 | protein_coding-intron (ENSMUST00000020886, intron 3 of 10) | protein_coding | 13207 | ENSMUST00000218491 | ENSMUSG00000020572 | ENSMUST00000020886 |
| 43 | Rnase10 | 14 | 51015187 | 51015687 | 4.56 | 500 | Intergenic | protein_coding | 7508 | ENSMUST00000022424 | ENSMUSG00000021872 | ENSMUST00000164632 |
| 44 | Hipk2 | 6 | 38827283 | 38827783 | 4.56 | 500 | protein_coding-intron (ENSMUST00000114855, intron 1 of 1) | protein_coding | -9187 | ENSMUST00000159894 | ENSMUSG00000061436 | ENSMUST00000114855 |
| 45 | Rfc1 | 5 | 65286954 | 65287454 | 4.54 | 500 | protein_coding-intron (ENSMUST00000172732, intron 11 of 24) | protein_coding | -13537 | ENSMUST00000205063 | ENSMUSG00000029191 | ENSMUST00000172732 |
| 46 | Tmem167 | 13 | 90090531 | 90091031 | 4.54 | 500 | protein_coding-intron (ENSMUST00000012566, intron 1 of 3) | protein_coding | 1019 | ENSMUST00000012566 | ENSMUSG00000012422 | ENSMUST00000161568 |
| 47 | Arid5b | 10 | 68258281 | 68258781 | 4.48 | 500 | protein_coding-intron (ENSMUST00000020106, intron 2 of 10) | protein_coding | 20190 | ENSMUST00000020106 | ENSMUSG00000019947 | ENSMUST00000218425 |
| 48 | Tnip1 | 11 | 54970124 | 54970624 | 4.48 | 500 | Intergenic | protein_coding | -7457 | ENSMUST00000102731 | ENSMUSG00000020400 | ENSMUST00000108889 |
| 49 | Abca13 | 11 | 9344571 | 9345071 | 4.47 | 500 | protein_coding-intron (ENSMUST00000042740, intron 27 of 61) | protein_coding | 151370 | ENSMUST00000151522 | ENSMUSG00000004668 | ENSMUST00000151522 |
| 50 | Plxdc2 | 2 | 16522305 | 16522805 | 4.45 | 500 | protein_coding-intron (ENSMUST00000028081, intron 2 of 13) | protein_coding | -116141 | ENSMUST00000126173 | ENSMUSG00000026748 | ENSMUST00000114703 |
| 51 | Mast2 | 4 | 116396194 | 116396694 | 4.4 | 500 | protein_coding-intron (ENSMUST00000003908, intron 3 of 28) | protein_coding | 9542 | ENSMUST00000123072 | ENSMUSG00000003810 | ENSMUST00000123761 |
| 52 | Rapgef2 | 3 | 79134937 | 79135437 | 4.39 | 500 | protein_coding-intron (ENSMUST00000118100, intron 1 of 23) | protein_coding | 9790 | ENSMUST00000118100 | ENSMUSG00000062232 | ENSMUST00000152275 |
| 53 | Cenpk | 13 | 104260555 | 104261055 | 4.37 | 500 | Intergenic | protein_coding | 22536 | ENSMUST00000224098 | ENSMUSG00000021714 | ENSMUST00000224098 |
| 54 | Nav2 | 7 | 49000998 | 49001498 | 4.37 | 500 | protein_coding-intron (ENSMUST00000183659, intron 2 of 39) | protein_coding | 42151 | ENSMUST00000183659 | ENSMUSG00000052512 | ENSMUST00000184336 |
| 55 | Gpld1 | 13 | 24945467 | 24945967 | 4.36 | 500 | protein_coding-intron (ENSMUST00000021773, intron 2 of 25) | protein_coding | 2565 | ENSMUST00000021773 | ENSMUSG00000021340 | ENSMUST00000223873 |
| 56 | Gtf2ird1 | 5 | 134453355 | 134453855 | 4.35 | 500 | protein_coding-intron (ENSMUST00000073161, intron 1 of 30) | protein_coding | 2346 | ENSMUST00000201647 | ENSMUSG00000023079 | ENSMUST00000111245 |
| 57 | Cntn3 | 6 | 102608760 | 102609260 | 4.33 | 500 | Intergenic | protein_coding | -35909 | ENSMUST00000203619 | ENSMUSG00000030075 | ENSMUST00000203619 |
| 58 | Lin9 | 1 | 180667281 | 180667781 | 4.33 | 500 | protein_coding-intron (ENSMUST00000085804, intron 8 of 11) | protein_coding | 6054 | ENSMUST00000194638 | ENSMUSG00000058729 | ENSMUST00000192561 |
| 59 | Rbmx2 | X | 48777269 | 48777769 | 4.29 | 500 | Intergenic | protein_coding | 71339 | ENSMUST00000139150 | ENSMUSG00000031107 | ENSMUST00000033433 |
| 60 | Bcl2l15 | 3 | 103842161 | 103842661 | 4.26 | 500 | protein_coding-intron (ENSMUST00000106822, intron 3 of 3) | protein_coding | -4413 | ENSMUST00000199772 | ENSMUSG00000044165 | ENSMUST00000062945 |
| 61 | Ogn | 13 | 49604827 | 49605327 | 4.26 | 500 | protein_coding-intron (ENSMUST00000021818, intron 5 of 7) | protein_coding | -2969 | ENSMUST00000021822 | ENSMUSG00000021390 | ENSMUST00000021822 |
| 62 | Atg10 | 13 | 91025135 | 91025635 | 4.24 | 500 | protein_coding-intron (ENSMUST00000022119, intron 4 of 7) | protein_coding | 100225 | ENSMUST00000223729 | ENSMUSG00000021619 | ENSMUST00000022119 |
| 63 | Casz1 | 4 | 148854696 | 148855196 | 4.24 | 500 | protein_coding-intron (ENSMUST00000094464, intron 2 of 15) | protein_coding | -34435 | ENSMUST00000140617 | ENSMUSG00000028977 | ENSMUST00000239035 |
| 64 | Picalm | 7 | 90172515 | 90173015 | 4.22 | 500 | protein_coding-intron (ENSMUST00000049537, intron 7 of 20) | protein_coding | 515 | ENSMUST00000208089 | ENSMUSG00000039361 | ENSMUST00000208089 |
| 65 | Snx7 | 3 | 117769321 | 117769821 | 4.22 | 500 | Intergenic | protein_coding | 61703 | ENSMUST00000196421 | ENSMUSG00000028007 | ENSMUST00000169812 |
| 66 | Nphp4 | 4 | 152466293 | 152466793 | 4.22 | 500 | protein_coding-intron (ENSMUST00000030768, intron 1 of 14) | protein_coding | -10163 | ENSMUST00000142027 | ENSMUSG00000039577 | ENSMUST00000123635 |
| 67 | Gfod1 | 13 | 43314088 | 43314588 | 4.2 | 500 | Intergenic | protein_coding | -10166 | ENSMUST00000055341 | ENSMUSG00000051335 | ENSMUST00000055341 |
| 68 | Rhbdf2 | 11 | 116625654 | 116626154 | 4.2 | 500 | protein_coding-intron (ENSMUST00000103028, intron 1 of 18) | protein_coding | 1115 | ENSMUST00000103028 | ENSMUSG00000020806 | ENSMUST00000103028 |
| 69 | Dthd1 | 5 | 63189464 | 63189964 | 4.18 | 500 | Intergenic | protein_coding | 375891 | ENSMUST00000170704 | ENSMUSG00000090326 | ENSMUST00000170704 |
| 70 | Mertk | 2 | 128711999 | 128712499 | 4.18 | 500 | protein_coding-intron (ENSMUST00000014505, intron 1 of 18) | protein_coding | 13293 | ENSMUST00000014505 | ENSMUSG00000014361 | ENSMUST00000014505 |
| 71 | Nudt3 | 17 | 27596437 | 27596937 | 4.16 | 500 | protein_coding-exon (ENSMUST00000146321, exon 2 of 2) | protein_coding | 528 | ENSMUST00000231742 | ENSMUSG00000024213 | ENSMUST00000176876 |
| 72 | Smc6 | 12 | 11290974 | 11291474 | 4.15 | 500 | protein_coding-promoter-TSS (ENSMUST00000217930) | protein_coding | 455 | ENSMUST00000220289 | ENSMUSG00000020608 | ENSMUST00000219041 |
| 73 | Anxa4 | 6 | 86818816 | 86819316 | 4.11 | 500 | Intergenic | protein_coding | -25482 | ENSMUST00000001187 | ENSMUSG00000029994 | ENSMUST00000155456 |
| 74 | Tsen2 | 6 | 115578973 | 115579473 | 4.11 | 500 | protein_coding-TTS (ENSMUST00000040234) | protein_coding | 4775 | ENSMUST00000143520 | ENSMUSG00000042389 | ENSMUST00000040234 |
| 75 | Filip1 | 9 | 79949576 | 79950076 | 4.09 | 500 | protein_coding-intron (ENSMUST00000093811, intron 1 of 5) | protein_coding | 12207 | ENSMUST00000172973 | ENSMUSG00000034898 | ENSMUST00000172973 |
| 76 | Erc1 | 6 | 119790667 | 119791167 | 4.09 | 500 | protein_coding-intron (ENSMUST00000032279, intron 2 of 18) | protein_coding | 5997 | ENSMUST00000183872 | ENSMUSG00000030172 | ENSMUST00000184838 |
| 77 | Tcf7l2 | 19 | 55905021 | 55905521 | 4.09 | 500 | protein_coding-intron (ENSMUST00000041717, intron 4 of 11) | protein_coding | 6962 | ENSMUST00000148666 | ENSMUSG00000024985 | ENSMUST00000041717 |
| 78 | Ccdc34 | 2 | 110044224 | 110044724 | 4.06 | 500 | protein_coding-exon (ENSMUST00000028580, exon 6 of 6) | protein_coding | 3579 | ENSMUST00000155110 | ENSMUSG00000027160 | ENSMUST00000155110 |
| 79 | Ubtd1 | 19 | 42010322 | 42010822 | 4.05 | 500 | protein_coding-intron (ENSMUST00000026170, intron 1 of 2) | protein_coding | -6577 | ENSMUST00000237817 | ENSMUSG00000025171 | ENSMUST00000026170 |
| 80 | Crem | 18 | 3348761 | 3349261 | 4.04 | 500 | lncRNA-intron (ENSMUST00000181558, intron 1 of 1) | protein_coding | -11263 | ENSMUST00000154470 | ENSMUSG00000063889 | ENSMUST00000142690 |
| 81 | Pde4d | 13 | 108983642 | 108984142 | 4.04 | 500 | protein_coding-intron (ENSMUST00000122041, intron 2 of 16) | protein_coding | 123726 | ENSMUST00000177907 | ENSMUSG00000021699 | ENSMUST00000129374 |
| 82 | Arhgef6 | X | 57355804 | 57356304 | 4.04 | 500 | Intergenic | protein_coding | -17325 | ENSMUST00000033468 | ENSMUSG00000031133 | ENSMUST00000175900 |
| 83 | Camk2d | 3 | 126806947 | 126807447 | 4.04 | 500 | protein_coding-promoter-TSS (ENSMUST00000198637) | protein_coding | 35 | ENSMUST00000198637 | ENSMUSG00000053819 | ENSMUST00000131156 |
| 84 | Gbe1 | 16 | 70551367 | 70551867 | 4.04 | 500 | protein_coding-intron (ENSMUST00000023393, intron 14 of 16) | protein_coding | -9377 | ENSMUST00000169432 | ENSMUSG00000022707 | ENSMUST00000169432 |
| 85 | Mctp2 | 7 | 72073624 | 72074124 | 4.04 | 500 | Intergenic | protein_coding | 36709 | ENSMUST00000206466 | ENSMUSG00000032776 | ENSMUST00000205410 |
| 86 | Rasgef1c | 11 | 49923087 | 49923587 | 4.04 | 500 | protein_coding-intron (ENSMUST00000063444, intron 1 of 13) | protein_coding | 21176 | ENSMUST00000063444 | ENSMUSG00000020374 | ENSMUST00000109172 |
| 87 | Snx2 | 18 | 53170140 | 53170640 | 4.04 | 500 | Intergenic | protein_coding | -5926 | ENSMUST00000037850 | ENSMUSG00000034484 | ENSMUST00000236911 |
| 88 | Nasp | 4 | 116613165 | 116613665 | 4.02 | 500 | protein_coding-intron (ENSMUST00000030456, intron 5 of 15) | protein_coding | -7167 | ENSMUST00000155398 | ENSMUSG00000028693 | ENSMUST00000155398 |
| 89 | Rgl1 | 1 | 152588813 | 152589313 | 4.02 | 500 | protein_coding-intron (ENSMUST00000027760, intron 2 of 17) | protein_coding | 3696 | ENSMUST00000128251 | ENSMUSG00000026482 | ENSMUST00000128251 |
| 90 | Apool | X | 112350887 | 112351387 | 3.99 | 500 | protein_coding-intron (ENSMUST00000026599, intron 4 of 8) | protein_coding | 39771 | ENSMUST00000113415 | ENSMUSG00000025525 | ENSMUST00000154378 |
| 91 | Fbxw17 | 13 | 50395224 | 50395724 | 3.99 | 500 | lncRNA-intron (ENSMUST00000185962, intron 1 of 1) | protein_coding | -22390 | ENSMUST00000046974 | ENSMUSG00000037816 | ENSMUST00000176772 |
| 92 | Haus3 | 5 | 34163121 | 34163621 | 3.99 | 500 | snRNA-TTS (ENSMUST00000104266) | protein_coding | 341 | ENSMUST00000201042 | ENSMUSG00000079555 | ENSMUST00000201042 |
| 93 | Mpp7 | 18 | 7614238 | 7614738 | 3.99 | 500 | protein_coding-intron (ENSMUST00000115869, intron 1 of 16) | protein_coding | -5506 | ENSMUST00000234510 | ENSMUSG00000057440 | ENSMUST00000235093 |
| 94 | Ppp1cb | 5 | 32490516 | 32491016 | 3.99 | 500 | protein_coding-intron (ENSMUST00000015100, intron 7 of 7) | protein_coding | 4796 | ENSMUST00000201880 | ENSMUSG00000014956 | ENSMUST00000201880 |
| 95 | Arap2 | 5 | 62573281 | 62573781 | 3.97 | 500 | Intergenic | protein_coding | 38843 | ENSMUST00000160620 | ENSMUSG00000037999 | ENSMUST00000159470 |
| 96 | BC052040 | 2 | 115608213 | 115608713 | 3.95 | 500 | protein_coding-intron (ENSMUST00000110918, intron 1 of 10) | protein_coding | 26662 | ENSMUST00000147968 | ENSMUSG00000040282 | ENSMUST00000166472 |
| 97 | Prune2 | 19 | 17191023 | 17191523 | 3.95 | 500 | protein_coding-intron (ENSMUST00000087689, intron 11 of 19) | protein_coding | -16978 | ENSMUST00000225838 | ENSMUSG00000039126 | ENSMUST00000087689 |
| 98 | Fmnl2 | 2 | 52957542 | 52958042 | 3.94 | 500 | protein_coding-intron (ENSMUST00000049483, intron 1 of 26) | protein_coding | 99924 | ENSMUST00000050719 | ENSMUSG00000036053 | ENSMUST00000090952 |
| 99 | Lrp12 | 15 | 39922807 | 39923307 | 3.94 | 500 | protein_coding-intron (ENSMUST00000022916, intron 1 of 6) | protein_coding | 20633 | ENSMUST00000228575 | ENSMUSG00000022305 | ENSMUST00000110305 |
| 100 | Zfp281 | 1 | 136645472 | 136645972 | 3.94 | 500 | Intergenic | protein_coding | 20821 | ENSMUST00000112046 | ENSMUSG00000041483 | ENSMUST00000047734 |
| 101 | Phf11 | 14 | 59247955 | 59248634 | 3.93 | 679 | unprocessed_pseudogene-intron (ENSMUST00000224042, intron 3 of 5) | unprocessed_pseudogene | 11886 | ENSMUST00000224042 | ENSMUSG00000090881 | ENSMUST00000224042 |
| 102 | Taf4b | 18 | 14798426 | 14798926 | 3.9 | 500 | protein_coding-intron (ENSMUST00000169862, intron 2 of 14) | protein_coding | 15431 | ENSMUST00000169862 | ENSMUSG00000054321 | ENSMUST00000234789 |
| 103 | Gtse1 | 15 | 85847436 | 85847936 | 3.9 | 500 | protein_coding-intron (ENSMUST00000133158, intron 8 of 12) | protein_coding | -12059 | ENSMUST00000170629 | ENSMUSG00000022385 | ENSMUST00000170629 |
| 104 | Jam3 | 9 | 27154738 | 27155238 | 3.9 | 500 | protein_coding-intron (ENSMUST00000034472, intron 1 of 8) | protein_coding | 405 | ENSMUST00000034472 | ENSMUSG00000031990 | ENSMUST00000034472 |
| 105 | Lpin2 | 17 | 71238237 | 71238737 | 3.88 | 500 | protein_coding-promoter-TSS (ENSMUST00000140150) | protein_coding | -175 | ENSMUST00000140150 | ENSMUSG00000024052 | ENSMUST00000126681 |
| 106 | Mllt3 | 4 | 87921309 | 87921809 | 3.86 | 500 | protein_coding-intron (ENSMUST00000078090, intron 2 of 10) | protein_coding | -47017 | ENSMUST00000134555 | ENSMUSG00000028496 | ENSMUST00000128930 |
| 107 | Tnr | 1 | 159837255 | 159837755 | 3.86 | 500 | protein_coding-intron (ENSMUST00000111669, intron 1 of 21) | protein_coding | -12892 | ENSMUST00000192457 | ENSMUSG00000015829 | ENSMUST00000111669 |
| 108 | Dab2ip | 2 | 35577342 | 35577842 | 3.86 | 500 | protein_coding-intron (ENSMUST00000112986, intron 1 of 16) | protein_coding | -3468 | ENSMUST00000112987 | ENSMUSG00000026883 | ENSMUST00000145698 |
| 109 | Krt18 | 15 | 102032516 | 102033016 | 3.86 | 500 | protein_coding-TTS (ENSMUST00000023803) | protein_coding | 4586 | ENSMUST00000023803 | ENSMUSG00000023043 | ENSMUST00000023803 |
| 110 | Tubg1 | 11 | 101123359 | 101123859 | 3.84 | 500 | protein_coding-promoter-TSS (ENSMUST00000137844) | protein_coding | -891 | ENSMUST00000137844 | ENSMUSG00000035198 | ENSMUST00000137844 |
| 111 | Ccdc90b | 7 | 92514663 | 92515163 | 3.81 | 500 | Intergenic | protein_coding | -46236 | ENSMUST00000032842 | ENSMUSG00000030613 | ENSMUST00000153776 |
| 112 | Haus5 | 7 | 30666597 | 30667097 | 3.81 | 500 | Intergenic | protein_coding | -1853 | ENSMUST00000019697 | ENSMUSG00000078762 | ENSMUST00000132982 |
| 113 | Stx3 | 19 | 11821755 | 11822255 | 3.81 | 500 | Intergenic | protein_coding | -2602 | ENSMUST00000047698 | ENSMUSG00000041488 | ENSMUST00000209304 |
| 114 | Acsm3 | 7 | 119772261 | 119772761 | 3.79 | 500 | protein_coding-intron (ENSMUST00000063770, intron 4 of 14) | protein_coding | -2033 | ENSMUST00000149766 | ENSMUSG00000030935 | ENSMUST00000154828 |
| 115 | Sfmbt2 | 2 | 10372835 | 10373335 | 3.79 | 500 | protein_coding-intron (ENSMUST00000041105, intron 1 of 22) | protein_coding | 635 | ENSMUST00000041105 | ENSMUSG00000061186 | ENSMUST00000145398 |
| 116 | Cdan1 | 2 | 120778376 | 120778876 | 3.77 | 500 | protein_coding-intron (ENSMUST00000028740, intron 7 of 15) | protein_coding | -47108 | ENSMUST00000110700 | ENSMUSG00000027284 | ENSMUST00000150860 |
| 117 | Ica1 | 6 | 8823570 | 8824070 | 3.77 | 500 | Intergenic | protein_coding | -45332 | ENSMUST00000135113 | ENSMUSG00000062995 | ENSMUST00000126430 |
| 118 | Prkcq | 2 | 11152672 | 11153172 | 3.77 | 500 | Intergenic | protein_coding | -19186 | ENSMUST00000028118 | ENSMUSG00000026778 | ENSMUST00000114853 |
| 119 | Stard4 | 18 | 33284170 | 33284670 | 3.77 | 500 | Intergenic | protein_coding | -70558 | ENSMUST00000025236 | ENSMUSG00000024378 | ENSMUST00000141617 |
| 120 | Ift172 | 5 | 31264606 | 31265106 | 3.77 | 500 | protein_coding-promoter-TSS (ENSMUST00000201953) | protein_coding | -935 | ENSMUST00000201953 | ENSMUSG00000038564 | ENSMUST00000202410 |
| 121 | Oas1g | 5 | 120879024 | 120879524 | 3.77 | 500 | protein_coding-exon (ENSMUST00000086368, exon 4 of 7) | protein_coding | 329 | ENSMUST00000161809 | ENSMUSG00000066861 | ENSMUST00000086368 |
| 122 | Gramd1b | 9 | 40317260 | 40317760 | 3.75 | 500 | protein_coding-TTS (ENSMUST00000137189) | protein_coding | 3410 | ENSMUST00000239052 | ENSMUSG00000040111 | ENSMUST00000239052 |
| 123 | Top2a | 11 | 98990695 | 98991195 | 3.75 | 500 | Intergenic | protein_coding | 4425 | ENSMUST00000139460 | ENSMUSG00000020914 | ENSMUST00000129777 |
| 124 | Bub1b | 2 | 118633226 | 118633726 | 3.73 | 500 | protein_coding-intron (ENSMUST00000038341, intron 18 of 22) | protein_coding | 1936 | ENSMUST00000126013 | ENSMUSG00000040084 | ENSMUST00000130512 |
| 125 | Tet1 | 10 | 62812882 | 62813382 | 3.73 | 500 | protein_coding-TTS (ENSMUST00000173087) | protein_coding | 1407 | ENSMUST00000173087 | ENSMUSG00000047146 | ENSMUST00000218438 |
| 126 | Chl1 | 6 | 103489392 | 103489892 | 3.72 | 500 | Intergenic | protein_coding | -20944 | ENSMUST00000203912 | ENSMUSG00000030077 | ENSMUST00000203912 |
| 127 | Fancb | X | 164983262 | 164983762 | 3.72 | 500 | protein_coding-intron (ENSMUST00000057150, intron 2 of 8) | protein_coding | 2800 | ENSMUST00000057150 | ENSMUSG00000047757 | ENSMUST00000057150 |
| 128 | Ltbp1 | 17 | 75018209 | 75018709 | 3.72 | 500 | protein_coding-intron (ENSMUST00000001927, intron 2 of 33) | protein_coding | 12165 | ENSMUST00000146839 | ENSMUSG00000001870 | ENSMUST00000234008 |
| 129 | Neil3 | 8 | 53678903 | 53679403 | 3.72 | 500 | Intergenic | protein_coding | -40088 | ENSMUST00000047768 | ENSMUSG00000039396 | ENSMUST00000047768 |
| 130 | Dbf4 | 5 | 8408470 | 8408970 | 3.7 | 500 | protein_coding-intron (ENSMUST00000002368, intron 4 of 11) | protein_coding | -3480 | ENSMUST00000197442 | ENSMUSG00000002297 | ENSMUST00000166364 |
| 131 | Gbp9 | 5 | 105083953 | 105084453 | 3.7 | 500 | protein_coding-intron (ENSMUST00000031235, intron 1 of 9) | protein_coding | -1440 | ENSMUST00000199453 | ENSMUSG00000029298 | ENSMUST00000031238 |
| 132 | Mttp | 3 | 138159766 | 138160266 | 3.7 | 500 | protein_coding-TTS (ENSMUST00000040321) | protein_coding | -15048 | ENSMUST00000197680 | ENSMUSG00000028158 | ENSMUST00000029805 |
| 133 | Dusp10 | 1 | 184050531 | 184051031 | 3.69 | 500 | protein_coding-intron (ENSMUST00000048655, intron 2 of 3) | protein_coding | 16400 | ENSMUST00000048655 | ENSMUSG00000039384 | ENSMUST00000048655 |
| 134 | Slco3a1 | 7 | 74558845 | 74559345 | 3.68 | 500 | Intergenic | protein_coding | -4315 | ENSMUST00000107453 | ENSMUSG00000025790 | ENSMUST00000129292 |
| 135 | Macrod2 | 2 | 141463136 | 141463636 | 3.68 | 500 | protein_coding-intron (ENSMUST00000078027, intron 5 of 17) | protein_coding | -92290 | ENSMUST00000138786 | ENSMUSG00000068205 | ENSMUST00000124403 |
| 136 | Zbtb32 | 7 | 30602403 | 30602903 | 3.68 | 500 | protein_coding-TTS (ENSMUST00000006476) | protein_coding | -3744 | ENSMUST00000144532 | ENSMUSG00000006310 | ENSMUST00000108150 |
| 137 | Chka | 19 | 3884432 | 3884932 | 3.67 | 500 | protein_coding-TTS (ENSMUST00000146463) | protein_coding | -7049 | ENSMUST00000125013 | ENSMUSG00000024843 | ENSMUST00000128694 |
| 138 | Kcnab1 | 3 | 65264421 | 65264921 | 3.67 | 500 | protein_coding-intron (ENSMUST00000049230, intron 1 of 13) | protein_coding | 11243 | ENSMUST00000161979 | ENSMUSG00000027827 | ENSMUST00000160136 |
| 139 | Rasgrp1 | 2 | 117320493 | 117320993 | 3.67 | 500 | protein_coding-intron (ENSMUST00000102534, intron 2 of 16) | protein_coding | 21966 | ENSMUST00000174770 | ENSMUSG00000027347 | ENSMUST00000110898 |
| 140 | Adamts3 | 5 | 89947302 | 89947802 | 3.65 | 500 | Intergenic | protein_coding | -64218 | ENSMUST00000198151 | ENSMUSG00000043635 | ENSMUST00000198151 |
| 141 | Stag3 | 5 | 138289028 | 138289528 | 3.65 | 500 | protein_coding-exon (ENSMUST00000048028, exon 5 of 34) | protein_coding | 8722 | ENSMUST00000161691 | ENSMUSG00000036928 | ENSMUST00000161113 |
| 142 | Tes | 6 | 17116084 | 17116584 | 3.65 | 500 | lncRNA-intron (ENSMUST00000150251, intron 1 of 1) | protein_coding | 51110 | ENSMUST00000076654 | ENSMUSG00000029552 | ENSMUST00000154266 |
| 143 | Capg | 6 | 72542240 | 72542740 | 3.65 | 500 | Intergenic | protein_coding | -1901 | ENSMUST00000071044 | ENSMUSG00000056737 | ENSMUST00000114071 |
| 144 | Ifi203 | 1 | 173959740 | 173960240 | 3.65 | 500 | Intergenic | protein_coding | -17318 | ENSMUST00000156895 | ENSMUSG00000039997 | ENSMUST00000156895 |
| 145 | Mad1l1 | 5 | 140241481 | 140241981 | 3.65 | 500 | protein_coding-intron (ENSMUST00000031534, intron 10 of 17) | protein_coding | 20834 | ENSMUST00000132832 | ENSMUSG00000029554 | ENSMUST00000110829 |
| 146 | Tns3 | 11 | 8426384 | 8426884 | 3.63 | 500 | Intergenic | protein_coding | 42041 | ENSMUST00000129074 | ENSMUSG00000020422 | ENSMUST00000134823 |
| 147 | Arl6 | 16 | 59737795 | 59738295 | 3.63 | 500 | protein_coding-intron (ENSMUST00000068860, intron 14 of 17) | protein_coding | -98654 | ENSMUST00000099646 | ENSMUSG00000022722 | ENSMUST00000149797 |
| 148 | Cndp2 | 18 | 84691472 | 84691972 | 3.63 | 500 | Intergenic | protein_coding | -6020 | ENSMUST00000025546 | ENSMUSG00000024644 | ENSMUST00000236415 |
| 149 | Kif2c | 4 | 117171652 | 117172152 | 3.62 | 500 | protein_coding-intron (ENSMUST00000065896, intron 7 of 19) | protein_coding | -6492 | ENSMUST00000132868 | ENSMUSG00000028678 | ENSMUST00000142205 |
| 150 | Pwwp2b | 7 | 139243991 | 139244491 | 3.62 | 500 | Intergenic | protein_coding | -4241 | ENSMUST00000093993 | ENSMUSG00000060260 | ENSMUST00000209721 |
| 151 | Tnk2 | 16 | 32635401 | 32635901 | 3.62 | 500 | Intergenic | protein_coding | -8223 | ENSMUST00000145627 | ENSMUSG00000022791 | ENSMUST00000150383 |
| 152 | Dusp16 | 6 | 134771852 | 134772352 | 3.61 | 500 | protein_coding-intron (ENSMUST00000100857, intron 1 of 6) | protein_coding | -11002 | ENSMUST00000204083 | ENSMUSG00000030203 | ENSMUST00000204083 |
| 153 | Klf5 | 14 | 99313784 | 99314284 | 3.61 | 500 | protein_coding-exon (ENSMUST00000005279, exon 4 of 4) | protein_coding | 14880 | ENSMUST00000226784 | ENSMUSG00000005148 | ENSMUST00000226784 |
| 154 | Bcan | 3 | 87996287 | 87996787 | 3.6 | 500 | protein_coding-promoter-TSS (ENSMUST00000193276) | protein_coding | -172 | ENSMUST00000193276 | ENSMUSG00000004892 | ENSMUST00000191627 |
| 155 | Ocln | 13 | 100547610 | 100548110 | 3.6 | 500 | protein_coding-intron (ENSMUST00000022140, intron 1 of 8) | protein_coding | 4600 | ENSMUST00000159515 | ENSMUSG00000021638 | ENSMUST00000159459 |
| 156 | Akr1d1 | 6 | 37537882 | 37538382 | 3.59 | 500 | protein_coding-intron (ENSMUST00000040987, intron 3 of 8) | protein_coding | -1549 | ENSMUST00000123648 | ENSMUSG00000038641 | ENSMUST00000123648 |
| 157 | Dach1 | 14 | 98160264 | 98160764 | 3.59 | 500 | protein_coding-intron (ENSMUST00000069334, intron 1 of 10) | protein_coding | 9029 | ENSMUST00000069334 | ENSMUSG00000055639 | ENSMUST00000069334 |
| 158 | Parp14 | 16 | 35877544 | 35878044 | 3.59 | 500 | Intergenic | protein_coding | -6250 | ENSMUST00000042665 | ENSMUSG00000034422 | ENSMUST00000042665 |
| 159 | Cenpi | X | 134350195 | 134350695 | 3.56 | 500 | protein_coding-intron (ENSMUST00000081064, intron 19 of 22) | protein_coding | 3622 | ENSMUST00000125604 | ENSMUSG00000031262 | ENSMUST00000125604 |
| 160 | Fst | 13 | 114481461 | 114481961 | 3.56 | 500 | Intergenic | protein_coding | -22760 | ENSMUST00000223640 | ENSMUSG00000021765 | ENSMUST00000223865 |
| 161 | Gstcd | 3 | 133071907 | 133072407 | 3.56 | 500 | protein_coding-intron (ENSMUST00000029651, intron 3 of 11) | protein_coding | 10971 | ENSMUST00000197381 | ENSMUSG00000028018 | ENSMUST00000161577 |
| 162 | Hmgcr | 13 | 96673658 | 96674158 | 3.56 | 500 | Intergenic | protein_coding | -2972 | ENSMUST00000022176 | ENSMUSG00000021670 | ENSMUST00000168855 |
| 163 | Pi4k2b | 5 | 52709806 | 52710306 | 3.56 | 500 | Intergenic | protein_coding | -31518 | ENSMUST00000031081 | ENSMUSG00000029186 | ENSMUST00000031082 |
| 164 | Gys1 | 7 | 45436208 | 45436708 | 3.56 | 500 | protein_coding-intron (ENSMUST00000003964, intron 1 of 15) | protein_coding | 1581 | ENSMUST00000211150 | ENSMUSG00000003865 | ENSMUST00000210563 |
| 165 | Limk2 | 11 | 3378242 | 3378742 | 3.56 | 500 | protein_coding-intron (ENSMUST00000101638, intron 2 of 15) | protein_coding | -7116 | ENSMUST00000101642 | ENSMUSG00000020451 | ENSMUST00000101642 |
| 166 | Alms1 | 6 | 85570460 | 85570960 | 3.54 | 500 | Intergenic | protein_coding | -16821 | ENSMUST00000072018 | ENSMUSG00000063810 | ENSMUST00000072018 |
| 167 | C6 | 15 | 4703046 | 4703546 | 3.54 | 500 | Intergenic | protein_coding | -23879 | ENSMUST00000162585 | ENSMUSG00000022181 | ENSMUST00000162585 |
| 168 | Prdx6 | 1 | 161245692 | 161246192 | 3.54 | 500 | protein_coding-intron (ENSMUST00000051925, intron 3 of 5) | protein_coding | 4612 | ENSMUST00000139387 | ENSMUSG00000026701 | ENSMUST00000130867 |
| 169 | Znhit3 | 11 | 84919976 | 84920476 | 3.54 | 500 | Intergenic | protein_coding | -3860 | ENSMUST00000103195 | ENSMUSG00000020526 | ENSMUST00000144552 |
| 170 | Dlg2 | 7 | 92293715 | 92294215 | 3.52 | 500 | protein_coding-intron (ENSMUST00000074273, intron 14 of 22) | protein_coding | 58838 | ENSMUST00000208377 | ENSMUSG00000052572 | ENSMUST00000129818 |
| 171 | Palld | 8 | 61669666 | 61670166 | 3.52 | 500 | protein_coding-intron (ENSMUST00000034057, intron 10 of 19) | protein_coding | -78774 | ENSMUST00000121200 | ENSMUSG00000058056 | ENSMUST00000135439 |
| 172 | Inpp5b | 4 | 124779977 | 124780477 | 3.52 | 500 | protein_coding-intron (ENSMUST00000094782, intron 8 of 23) | protein_coding | 10899 | ENSMUST00000142652 | ENSMUSG00000028894 | ENSMUST00000142652 |
| 173 | Myo1d | 11 | 80755611 | 80756111 | 3.52 | 500 | protein_coding-intron (ENSMUST00000041065, intron 1 of 21) | protein_coding | 24142 | ENSMUST00000125944 | ENSMUSG00000035441 | ENSMUST00000041065 |
| 174 | Cox7c | 13 | 86058970 | 86059470 | 3.47 | 500 | Intergenic | protein_coding | -12316 | ENSMUST00000131011 | ENSMUSG00000017778 | ENSMUST00000017922 |
| 175 | Tm4sf1 | 3 | 57293050 | 57293550 | 3.47 | 500 | protein_coding-exon (ENSMUST00000196704, exon 1 of 2) | protein_coding | 403 | ENSMUST00000196704 | ENSMUSG00000027800 | ENSMUST00000196704 |
| 176 | Megf9 | 4 | 70764609 | 70765109 | 3.45 | 500 | Intergenic | protein_coding | -229864 | ENSMUST00000037881 | ENSMUSG00000039270 | ENSMUST00000107359 |
| 177 | Syne2 | 12 | 75981850 | 75982350 | 3.45 | 500 | protein_coding-intron (ENSMUST00000044217, intron 53 of 115) | protein_coding | -40200 | ENSMUST00000148825 | ENSMUSG00000063450 | ENSMUST00000143031 |
| 178 | Ralgps2 | 1 | 156825557 | 156826057 | 3.4 | 500 | protein_coding-intron (ENSMUST00000027886, intron 13 of 18) | protein_coding | 5040 | ENSMUST00000191503 | ENSMUSG00000026594 | ENSMUST00000172057 |
| 179 | Hs3st3b1 | 11 | 63899805 | 63900305 | 3.39 | 500 | protein_coding-intron (ENSMUST00000094103, intron 1 of 1) | protein_coding | 22235 | ENSMUST00000094103 | ENSMUSG00000070407 | ENSMUST00000094103 |
| 180 | Nos2 | 11 | 78879479 | 78879979 | 3.37 | 500 | Intergenic | protein_coding | -41058 | ENSMUST00000018610 | ENSMUSG00000020826 | ENSMUST00000208783 |
| 181 | Mapkapk2 | 1 | 131098748 | 131099248 | 3.35 | 500 | Intergenic | protein_coding | -1172 | ENSMUST00000016672 | ENSMUSG00000016528 | ENSMUST00000016672 |
| 182 | Rpn1 | 6 | 88094870 | 88095370 | 3.31 | 500 | protein_coding-intron (ENSMUST00000032143, intron 4 of 9) | protein_coding | 1303 | ENSMUST00000203813 | ENSMUSG00000030062 | ENSMUST00000204164 |
| 183 | Tlr1 | 5 | 64930055 | 64930555 | 3.24 | 500 | protein_coding-intron (ENSMUST00000059349, intron 1 of 2) | protein_coding | 2456 | ENSMUST00000059349 | ENSMUSG00000044827 | ENSMUST00000059349 |

H3K9me3

| | Gene.Name | Chr | Start | End | Peak.Score | Focus.Ratio.Region.Size | Detailed.Annotation | Gene.Type | Distance.to.TSS | Nearest.PromoterID | Entrez.ID | Nearest.Unigene |
|---|---|---|---|---|---|---|---|---|---|---|---|---|
| 1 | Cntn3 | 6 | 102488141 | 102488953 | 9.18 | 812 | protein_coding-intron (ENSMUST00000203619, intron 1 of 22) | protein_coding | -23880 | ENSMUST00000032159 | ENSMUSG00000030075 | ENSMUST00000032159 |
| 2 | Macrod2 | 2 | 141566506 | 141567272 | 8.25 | 766 | protein_coding-intron (ENSMUST00000078027, intron 6 of 17) | protein_coding | 11213 | ENSMUST00000138786 | ENSMUSG00000068205 | ENSMUST00000124403 |
| 3 | Cks2 | 13 | 51638845 | 51639345 | 8.12 | 500 | Intergenic | protein_coding | -6137 | ENSMUST00000075853 | ENSMUSG00000062248 | ENSMUST00000075853 |
| 4 | Rab32 | 10 | 10521239 | 10522149 | 7.92 | 910 | Intergenic | protein_coding | 36487 | ENSMUST00000220018 | ENSMUSG00000019832 | ENSMUST00000220018 |
| 5 | Gvin1 | 7 | 106235609 | 106236406 | 7.65 | 797 | Intergenic | protein_coding | -20681 | ENSMUST00000183409 | ENSMUSG00000045868 | ENSMUST00000006667 |
| 6 | Chl1 | 6 | 103446818 | 103447554 | 7.58 | 736 | Intergenic | protein_coding | -63400 | ENSMUST00000203912 | ENSMUSG00000030077 | ENSMUST00000066905 |
| 7 | Sec63 | 10 | 42776738 | 42777517 | 7.12 | 779 | protein_coding-intron (ENSMUST00000019937, intron 1 of 20) | protein_coding | 15321 | ENSMUST00000144228 | ENSMUSG00000019802 | ENSMUST00000105496 |
| 8 | Alpk1 | 3 | 127783525 | 127784242 | 6.92 | 717 | Intergenic | protein_coding | -3356 | ENSMUST00000029662 | ENSMUSG00000028028 | ENSMUST00000161239 |
| 9 | Ica1 | 6 | 8863080 | 8863772 | 6.65 | 692 | Intergenic | protein_coding | -84938 | ENSMUST00000135113 | ENSMUSG00000062995 | ENSMUST00000151758 |
| 10 | Adamts3 | 5 | 89952098 | 89952598 | 6.63 | 500 | Intergenic | protein_coding | -69014 | ENSMUST00000198151 | ENSMUSG00000043635 | ENSMUST00000198151 |
| 11 | Basp1 | 15 | 25381804 | 25382480 | 6.56 | 676 | protein_coding-intron (ENSMUST00000058845, intron 1 of 1) | protein_coding | 31556 | ENSMUST00000228597 | ENSMUSG00000045763 | ENSMUST00000228597 |
| 12 | C6 | 15 | 4689696 | 4690196 | 6.52 | 500 | Intergenic | protein_coding | -37229 | ENSMUST00000162585 | ENSMUSG00000022181 | ENSMUST00000161997 |
| 13 | Zfp114 | 7 | 24157080 | 24157705 | 6.42 | 625 | Intergenic | protein_coding | -17668 | ENSMUST00000205309 | ENSMUSG00000068962 | ENSMUST00000086010 |
| 14 | Card11 | 5 | 140916521 | 140917021 | 6.39 | 500 | protein_coding-intron (ENSMUST00000085786, intron 2 of 24) | protein_coding | -34566 | ENSMUST00000196169 | ENSMUSG00000036526 | ENSMUST00000196169 |
| 15 | Isoc1 | 18 | 58711757 | 58712385 | 6.32 | 628 | Intergenic | protein_coding | 52591 | ENSMUST00000238139 | ENSMUSG00000024601 | ENSMUST00000025503 |
| 16 | Esyt2 | 12 | 116329900 | 116330541 | 6.25 | 641 | protein_coding-intron (ENSMUST00000100986, intron 6 of 21) | protein_coding | 6044 | ENSMUST00000220804 | ENSMUSG00000021171 | ENSMUST00000223150 |
| 17 | Snx7 | 3 | 117841212 | 117841712 | 6.25 | 500 | protein_coding-intron (ENSMUST00000029639, intron 2 of 8) | protein_coding | 5246 | ENSMUST00000198499 | ENSMUSG00000028007 | ENSMUST00000167877 |
| 18 | Ablim1 | 19 | 57056438 | 57057120 | 6.19 | 682 | protein_coding-intron (ENSMUST00000079360, intron 15 of 25) | protein_coding | -9541 | ENSMUST00000137389 | ENSMUSG00000025085 | ENSMUST00000111546 |
| 19 | Trem1 | 17 | 48221571 | 48222225 | 6.19 | 654 | Intergenic | protein_coding | -10870 | ENSMUST00000113251 | ENSMUSG00000042265 | ENSMUST00000048782 |
| 20 | Gmpr | 13 | 45499865 | 45500610 | 6.14 | 745 | Intergenic | protein_coding | -7207 | ENSMUST00000000260 | ENSMUSG00000000253 | ENSMUST00000129176 |
| 21 | Msra | 14 | 64366127 | 64366756 | 6.14 | 629 | protein_coding-intron (ENSMUST00000067927, intron 6 of 10) | protein_coding | 50507 | ENSMUST00000210428 | ENSMUSG00000054733 | ENSMUST00000209513 |
| 22 | Abtb2 | 2 | 103560069 | 103560716 | 6 | 647 | Intergenic | protein_coding | -5918 | ENSMUST00000076212 | ENSMUSG00000032724 | ENSMUST00000138580 |
| 23 | Gstcd | 3 | 133001062 | 133001700 | 6 | 638 | protein_coding-intron (ENSMUST00000029651, intron 8 of 11) | protein_coding | 4296 | ENSMUST00000160402 | ENSMUSG00000028018 | ENSMUST00000029651 |
| 24 | Schip1 | 3 | 67991681 | 67992388 | 6 | 707 | protein_coding-intron (ENSMUST00000063263, intron 1 of 3) | protein_coding | -72768 | ENSMUST00000029346 | ENSMUSG00000027777 | ENSMUST00000182532 |
| 25 | Slc27a2 | 2 | 126570826 | 126571326 | 6 | 500 | protein_coding-intron (ENSMUST00000061491, intron 4 of 9) | protein_coding | 6316 | ENSMUST00000150947 | ENSMUSG00000027359 | ENSMUST00000150947 |
| 26 | Dlg2 | 7 | 92222001 | 92222677 | 5.99 | 676 | protein_coding-intron (ENSMUST00000074273, intron 13 of 22) | protein_coding | -12584 | ENSMUST00000138389 | ENSMUSG00000052572 | ENSMUST00000138389 |
| 27 | Casp12 | 9 | 5460627 | 5461127 | 5.99 | 500 | Intergenic | protein_coding | 115373 | ENSMUST00000151332 | ENSMUSG00000025887 | ENSMUST00000149520 |
| 28 | Cp | 3 | 19990342 | 19990986 | 5.92 | 644 | protein_coding-intron (ENSMUST00000003714, intron 18 of 19) | protein_coding | 1512 | ENSMUST00000173779 | ENSMUSG00000003617 | ENSMUST00000172860 |
| 29 | Aff2 | X | 69192280 | 69192780 | 5.85 | 500 | Intergenic | protein_coding | -167764 | ENSMUST00000033532 | ENSMUSG00000031189 | ENSMUST00000033532 |
| 30 | Reln | 5 | 22146939 | 22147439 | 5.85 | 500 | protein_coding-intron (ENSMUST00000062372, intron 6 of 63) | protein_coding | -12520 | ENSMUST00000162876 | ENSMUSG00000042453 | ENSMUST00000200318 |
| 31 | Plxdc2 | 2 | 16190894 | 16191394 | 5.79 | 500 | protein_coding-intron (ENSMUST00000146205, intron 35 of 38) | protein_coding | -165160 | ENSMUST00000028081 | ENSMUSG00000026748 | ENSMUST00000114703 |
| 32 | Epha7 | 4 | 28858453 | 28858953 | 5.72 | 500 | protein_coding-intron (ENSMUST00000029964, intron 3 of 16) | protein_coding | -13138 | ENSMUST00000129029 | ENSMUSG00000028289 | ENSMUST00000149030 |
| 33 | Tfpi2 | 6 | 3991033 | 3991533 | 5.72 | 500 | Intergenic | protein_coding | -2364 | ENSMUST00000183682 | ENSMUSG00000029664 | ENSMUST00000183682 |
| 34 | Tlr4 | 4 | 66877082 | 66877719 | 5.66 | 637 | protein_coding-intron (ENSMUST00000107365, intron 2 of 3) | protein_coding | 49198 | ENSMUST00000147008 | ENSMUSG00000039005 | ENSMUST00000147008 |
| 35 | Kif5c | 2 | 49586163 | 49586663 | 5.66 | 500 | Intergenic | protein_coding | -32885 | ENSMUST00000028102 | ENSMUSG00000026764 | ENSMUST00000138834 |
| 36 | E2f7 | 10 | 110704836 | 110705336 | 5.52 | 500 | Intergenic | protein_coding | -40353 | ENSMUST00000173294 | ENSMUSG00000020185 | ENSMUST00000173471 |
| 37 | Dthd1 | 5 | 63135515 | 63136015 | 5.46 | 500 | Intergenic | protein_coding | 321942 | ENSMUST00000170704 | ENSMUSG00000090326 | ENSMUST00000170704 |
| 38 | Csl | 10 | 99796356 | 99796856 | 5.45 | 500 | lncRNA-intron (ENSMUST00000217945, intron 2 of 4) | protein_coding | -36948 | ENSMUST00000056085 | ENSMUSG00000046934 | ENSMUST00000056085 |
| 39 | Dtd1 | 2 | 144667632 | 144668301 | 5.39 | 669 | protein_coding-intron (ENSMUST00000028917, intron 4 of 5) | protein_coding | -10132 | ENSMUST00000137406 | ENSMUSG00000027430 | ENSMUST00000145814 |
| 40 | Crem | 18 | 3235116 | 3235616 | 5.38 | 500 | Intergenic | protein_coding | 45712 | ENSMUST00000049942 | ENSMUSG00000063889 | ENSMUST00000154135 |
| 41 | Mgst1 | 6 | 138281671 | 138282171 | 5.38 | 500 | Intergenic | protein_coding | 139067 | ENSMUST00000125810 | ENSMUSG00000008540 | ENSMUST00000008684 |
| 42 | Psma1 | 7 | 114345059 | 114345559 | 5.38 | 500 | lncRNA-intron (ENSMUST00000163996, intron 1 of 3) | protein_coding | -69191 | ENSMUST00000033008 | ENSMUSG00000030751 | ENSMUST00000154909 |
| 43 | Aoah | 13 | 20992075 | 20992575 | 5.31 | 500 | protein_coding-intron (ENSMUST00000021757, intron 13 of 20) | protein_coding | -18308 | ENSMUST00000222135 | ENSMUSG00000021322 | ENSMUST00000222135 |
| 44 | B230118H07Rik | 2 | 101605559 | 101606059 | 5.26 | 500 | protein_coding-intron (ENSMUST00000090513, intron 2 of 6) | protein_coding | 4747 | ENSMUST00000136601 | ENSMUSG00000027165 | ENSMUST00000160722 |
| 45 | Fhl4 | 10 | 85090488 | 85090988 | 5.26 | 500 | Intergenic | protein_coding | 11757 | ENSMUST00000059383 | ENSMUSG00000050035 | ENSMUST00000216889 |
| 46 | Map2k6 | 11 | 110487810 | 110488310 | 5.19 | 500 | protein_coding-intron (ENSMUST00000020949, intron 2 of 11) | protein_coding | 9021 | ENSMUST00000133920 | ENSMUSG00000020623 | ENSMUST00000020949 |
| 47 | Tm4sf1 | 3 | 57308244 | 57308744 | 5.19 | 500 | Intergenic | protein_coding | -6506 | ENSMUST00000029376 | ENSMUSG00000027800 | ENSMUST00000196704 |
| 48 | Lrrc63 | 14 | 75090241 | 75090741 | 5.17 | 500 | protein_coding-intron (ENSMUST00000022574, intron 8 of 9) | protein_coding | 40390 | ENSMUST00000022574 | ENSMUSG00000021997 | ENSMUST00000022574 |
| 49 | Rnf144b | 13 | 47228137 | 47228637 | 5.17 | 500 | protein_coding-intron (ENSMUST00000068891, intron 3 of 7) | protein_coding | 34562 | ENSMUST00000110111 | ENSMUSG00000038068 | ENSMUST00000110111 |
| 50 | Neil3 | 8 | 53706565 | 53707065 | 5.12 | 500 | Intergenic | protein_coding | -67750 | ENSMUST00000047768 | ENSMUSG00000039396 | ENSMUST00000149053 |
| 51 | Gpsm2 | 3 | 108725602 | 108726102 | 5.12 | 500 | Intergenic | protein_coding | -3543 | ENSMUST00000029482 | ENSMUSG00000027883 | ENSMUST00000124043 |
| 52 | Ttc32 | 12 | 9172333 | 9172833 | 5.12 | 500 | Intergenic | protein_coding | 142555 | ENSMUST00000219470 | ENSMUSG00000066637 | ENSMUST00000085741 |
| 53 | Ndufa4 | 6 | 11808568 | 11809068 | 5.1 | 500 | Intergenic | protein_coding | 98575 | ENSMUST00000204084 | ENSMUSG00000029632 | ENSMUST00000031637 |
| 54 | Nr4a2 | 2 | 57111513 | 57112013 | 5.1 | 500 | protein_coding-TTS (ENSMUST00000140165) | protein_coding | 1309 | ENSMUST00000112627 | ENSMUSG00000026826 | ENSMUST00000112627 |
| 55 | Pkib | 10 | 57655748 | 57656248 | 5.06 | 500 | protein_coding-intron (ENSMUST00000066028, intron 2 of 4) | protein_coding | 5004 | ENSMUST00000175852 | ENSMUSG00000019876 | ENSMUST00000176225 |
| 56 | Tmem168 | 6 | 13581478 | 13581978 | 5.06 | 500 | protein_coding-exon (ENSMUST00000031554, exon 5 of 5) | protein_coding | 26020 | ENSMUST00000127050 | ENSMUSG00000029569 | ENSMUST00000146139 |
| 57 | Alg6 | 4 | 99728027 | 99728527 | 5.03 | 500 | protein_coding-exon (ENSMUST00000137473, exon 3 of 3) | protein_coding | 9125 | ENSMUST00000124147 | ENSMUSG00000073792 | ENSMUST00000107004 |
| 58 | Mastl | 2 | 23151164 | 23151664 | 5.03 | 500 | protein_coding-intron (ENSMUST00000028119, intron 1 of 11) | protein_coding | 4483 | ENSMUST00000136207 | ENSMUSG00000026779 | ENSMUST00000136207 |
| 59 | Cenpw | 10 | 30281822 | 30282322 | 4.99 | 500 | Intergenic | protein_coding | -81512 | ENSMUST00000099985 | ENSMUSG00000075266 | ENSMUST00000216853 |
| 60 | Mbnl3 | X | 51178405 | 51178905 | 4.99 | 500 | protein_coding-intron (ENSMUST00000114875, intron 1 of 6) | protein_coding | -13981 | ENSMUST00000136404 | ENSMUSG00000036109 | ENSMUST00000041495 |
| 61 | N4bp1 | 8 | 86833830 | 86834330 | 4.99 | 500 | protein_coding-intron (ENSMUST00000209389, intron 1 of 1) | protein_coding | 9965 | ENSMUST00000209389 | ENSMUSG00000031652 | ENSMUST00000034074 |
| 62 | Ccdc82 | 9 | 13272794 | 13273294 | 4.96 | 500 | protein_coding-intron (ENSMUST00000110583, intron 7 of 8) | protein_coding | 713 | ENSMUST00000215778 | ENSMUSG00000079084 | ENSMUST00000216578 |
| 63 | Gbp9 | 5 | 105071856 | 105072356 | 4.96 | 500 | protein_coding-intron (ENSMUST00000031235, intron 1 of 9) | protein_coding | 10657 | ENSMUST00000199453 | ENSMUSG00000029298 | ENSMUST00000199453 |
| 64 | Hira | 16 | 18905809 | 18906309 | 4.92 | 500 | protein_coding-intron (ENSMUST00000004222, intron 5 of 24) | protein_coding | -17014 | ENSMUST00000140748 | ENSMUSG00000022702 | ENSMUST00000232419 |
| 65 | Fbxo5 | 10 | 5774038 | 5774538 | 4.92 | 500 | Intergenic | protein_coding | 30312 | ENSMUST00000142200 | ENSMUSG00000019773 | ENSMUST00000019907 |
| 66 | Kctd14 | 7 | 97440823 | 97441323 | 4.92 | 500 | Intergenic | protein_coding | -10250 | ENSMUST00000206658 | ENSMUSG00000051727 | ENSMUST00000206658 |
| 67 | Pcmtd1 | 1 | 7151617 | 7152117 | 4.92 | 500 | TEC-exon (ENSMUST00000194854, exon 1 of 1) | protein_coding | -3000 | ENSMUST00000182388 | ENSMUSG00000051285 | ENSMUST00000182306 |
| 68 | Zfp281 | 1 | 136630310 | 136630810 | 4.92 | 500 | protein_coding-TTS (ENSMUST00000047734) | protein_coding | 5659 | ENSMUST00000112046 | ENSMUSG00000041483 | ENSMUST00000112046 |
| 69 | Dars | 1 | 128377233 | 128377733 | 4.89 | 500 | protein_coding-intron (ENSMUST00000027602, intron 8 of 15) | protein_coding | 2004 | ENSMUST00000186398 | ENSMUSG00000026356 | ENSMUST00000027602 |
| 70 | Gria3 | X | 41535112 | 41535612 | 4.89 | 500 | protein_coding-intron (ENSMUST00000076349, intron 3 of 15) | protein_coding | -74835 | ENSMUST00000126843 | ENSMUSG00000001986 | ENSMUST00000124169 |
| 71 | Mpp7 | 18 | 7389753 | 7390253 | 4.89 | 500 | protein_coding-intron (ENSMUST00000115869, intron 12 of 16) | protein_coding | -35337 | ENSMUST00000234210 | ENSMUSG00000057440 | ENSMUST00000234210 |
| 72 | Tmx4 | 2 | 134638698 | 134639198 | 4.89 | 500 | protein_coding-TTS (ENSMUST00000110119) | protein_coding | 5093 | ENSMUST00000110120 | ENSMUSG00000034723 | ENSMUST00000110120 |
| 73 | Trim37 | 11 | 87173479 | 87173979 | 4.89 | 500 | protein_coding-intron (ENSMUST00000041282, intron 12 of 23) | protein_coding | -12784 | ENSMUST00000152637 | ENSMUSG00000018548 | ENSMUST00000154138 |
| 74 | Gbe1 | 16 | 70473491 | 70473991 | 4.86 | 500 | protein_coding-intron (ENSMUST00000023393, intron 7 of 16) | protein_coding | -87253 | ENSMUST00000169432 | ENSMUSG00000022707 | ENSMUST00000169432 |
| 75 | Nox1 | X | 134235803 | 134236303 | 4.86 | 500 | protein_coding-intron (ENSMUST00000087541, intron 12 of 12) | protein_coding | -14097 | ENSMUST00000162833 | ENSMUSG00000031257 | ENSMUST00000159231 |
| 76 | Syne1 | 10 | 5385553 | 5386053 | 4.86 | 500 | protein_coding-intron (ENSMUST00000041639, intron 13 of 31) | protein_coding | -17317 | ENSMUST00000215887 | ENSMUSG00000096054 | ENSMUST00000217592 |
| 77 | Fignl1 | 11 | 11793367 | 11793867 | 4.82 | 500 | protein_coding-intron (ENSMUST00000123815, intron 1 of 2) | protein_coding | 7844 | ENSMUST00000123815 | ENSMUSG00000035455 | ENSMUST00000123815 |
| 78 | Pcca | 14 | 122592536 | 122593036 | 4.82 | 500 | protein_coding-intron (ENSMUST00000038374, intron 6 of 23) | protein_coding | 10102 | ENSMUST00000177312 | ENSMUSG00000041650 | ENSMUST00000138836 |
| 79 | Pnlip | 19 | 58655285 | 58655785 | 4.82 | 500 | protein_coding-intron (ENSMUST00000237451, intron 1 of 1) | protein_coding | 3037 | ENSMUST00000237451 | ENSMUSG00000046008 | ENSMUST00000237451 |
| 80 | Adamts5 | 16 | 85945802 | 85946302 | 4.79 | 500 | Intergenic | protein_coding | -44224 | ENSMUST00000023611 | ENSMUSG00000022894 | ENSMUST00000023611 |
| 81 | Kcnab1 | 3 | 65089872 | 65090372 | 4.79 | 500 | protein_coding-intron (ENSMUST00000193778, intron 1 of 1) | protein_coding | -19262 | ENSMUST00000049230 | ENSMUSG00000027827 | ENSMUST00000238901 |
| 82 | Ugt8a | 3 | 125869671 | 125870171 | 4.79 | 500 | protein_coding-intron (ENSMUST00000057944, intron 5 of 5) | protein_coding | 45964 | ENSMUST00000198610 | ENSMUSG00000032854 | ENSMUST00000057944 |
| 83 | Abca13 | 11 | 9260166 | 9260666 | 4.79 | 500 | protein_coding-intron (ENSMUST00000042740, intron 7 of 61) | protein_coding | 66965 | ENSMUST00000151522 | ENSMUSG00000004668 | ENSMUST00000042740 |
| 84 | Casp1 | 9 | 5222903 | 5223403 | 4.79 | 500 | Intergenic | protein_coding | -75355 | ENSMUST00000027015 | ENSMUSG00000025888 | ENSMUST00000238348 |
| 85 | Cdk5rap2 | 4 | 70279009 | 70279509 | 4.79 | 500 | protein_coding-intron (ENSMUST00000076541, intron 18 of 33) | protein_coding | -14579 | ENSMUST00000138561 | ENSMUSG00000039298 | ENSMUST00000144099 |
| 86 | Mast2 | 4 | 116361144 | 116361644 | 4.79 | 500 | protein_coding-intron (ENSMUST00000003908, intron 4 of 28) | protein_coding | 10338 | ENSMUST00000239239 | ENSMUSG00000003810 | ENSMUST00000147262 |
| 87 | Mcpt8 | 14 | 56086151 | 56086651 | 4.79 | 500 | Intergenic | protein_coding | -1128 | ENSMUST00000225107 | ENSMUSG00000022157 | ENSMUST00000015594 |
| 88 | Ccdc90b | 7 | 92562532 | 92563032 | 4.75 | 500 | protein_coding-intron (ENSMUST00000032842, intron 1 of 8) | protein_coding | 1633 | ENSMUST00000153776 | ENSMUSG00000030613 | ENSMUST00000153776 |
| 89 | Dapk2 | 9 | 66309125 | 66309625 | 4.75 | 500 | Intergenic | protein_coding | 41001 | ENSMUST00000129778 | ENSMUSG00000032380 | ENSMUST00000034944 |
| 90 | Nphp4 | 4 | 152505976 | 152506476 | 4.75 | 500 | protein_coding-intron (ENSMUST00000056567, intron 6 of 28) | protein_coding | 3312 | ENSMUST00000123635 | ENSMUSG00000039577 | ENSMUST00000153759 |
| 91 | Ssx2ip | 3 | 146417525 | 146418025 | 4.75 | 500 | protein_coding-exon (ENSMUST00000039021, exon 2 of 14) | protein_coding | -12754 | ENSMUST00000141103 | ENSMUSG00000036825 | ENSMUST00000106153 |
| 92 | Rab3gap1 | 1 | 127890248 | 127890748 | 4.72 | 500 | protein_coding-intron (ENSMUST00000037649, intron 4 of 23) | protein_coding | -10415 | ENSMUST00000150945 | ENSMUSG00000036104 | ENSMUST00000142662 |
| 93 | Tshr | 12 | 91523936 | 91524436 | 4.72 | 500 | protein_coding-intron (ENSMUST00000021346, intron 8 of 9) | protein_coding | -13880 | ENSMUST00000186437 | ENSMUSG00000020963 | ENSMUST00000021346 |
| 94 | Myo6 | 9 | 80160827 | 80161327 | 4.72 | 500 | Intergenic | protein_coding | -3954 | ENSMUST00000035889 | ENSMUSG00000033577 | ENSMUST00000138650 |
| 95 | 4930438A08Rik | 11 | 58259490 | 58259990 | 4.68 | 500 | Intergenic | protein_coding | -15059 | ENSMUST00000108834 | ENSMUSG00000069873 | ENSMUST00000208022 |
| 96 | Akap9 | 5 | 4056906 | 4057406 | 4.68 | 500 | protein_coding-intron (ENSMUST00000044492, intron 35 of 47) | protein_coding | -2094 | ENSMUST00000196918 | ENSMUSG00000040407 | ENSMUST00000196350 |
| 97 | Ccdc38 | 10 | 93545080 | 93545580 | 4.68 | 500 | protein_coding-intron (ENSMUST00000092215, intron 1 of 15) | protein_coding | 4698 | ENSMUST00000092215 | ENSMUSG00000036168 | ENSMUST00000140460 |
| 98 | Dpy30 | 17 | 74323201 | 74323701 | 4.68 | 500 | protein_coding-intron (ENSMUST00000164832, intron 1 of 4) | protein_coding | 493 | ENSMUST00000164832 | ENSMUSG00000024067 | ENSMUST00000233059 |
| 99 | Far2 | 6 | 147932814 | 147933314 | 4.68 | 500 | Intergenic | protein_coding | -114195 | ENSMUST00000032443 | ENSMUSG00000030303 | ENSMUST00000032443 |
| 100 | Fundc1 | X | 17593956 | 17594456 | 4.68 | 500 | Intergenic | protein_coding | -21881 | ENSMUST00000026016 | ENSMUSG00000025040 | ENSMUST00000026016 |
| 101 | Nipa2 | 7 | 55944998 | 55945498 | 4.68 | 500 | protein_coding-promoter-TSS (ENSMUST00000205773) | protein_coding | -301 | ENSMUST00000205773 | ENSMUSG00000030452 | ENSMUST00000130239 |
| 102 | Rab30 | 7 | 92777103 | 92777603 | 4.68 | 500 | protein_coding-promoter-TSS (ENSMUST00000208945) | protein_coding | -230 | ENSMUST00000208945 | ENSMUSG00000030643 | ENSMUST00000208945 |
| 103 | Tlr1 | 5 | 64929986 | 64930486 | 4.68 | 500 | protein_coding-intron (ENSMUST00000059349, intron 1 of 2) | protein_coding | 2525 | ENSMUST00000059349 | ENSMUSG00000044827 | ENSMUST00000197886 |
| 104 | Glt1d1 | 5 | 127724302 | 127724802 | 4.66 | 500 | Intergenic | protein_coding | 67519 | ENSMUST00000134529 | ENSMUSG00000049971 | ENSMUST00000137157 |
| 105 | Hcfc2 | 10 | 82705967 | 82706467 | 4.66 | 500 | protein_coding-intron (ENSMUST00000020478, intron 3 of 14) | protein_coding | 7006 | ENSMUST00000162422 | ENSMUSG00000020246 | ENSMUST00000160845 |
| 106 | Myo1b | 1 | 51818137 | 51818637 | 4.66 | 500 | protein_coding-intron (ENSMUST00000018561, intron 6 of 30) | protein_coding | 2159 | ENSMUST00000135125 | ENSMUSG00000018417 | ENSMUST00000114541 |
| 107 | Ola1 | 2 | 73219358 | 73219858 | 4.66 | 500 | protein_coding-promoter-TSS (ENSMUST00000133228) | protein_coding | -684 | ENSMUST00000133228 | ENSMUSG00000027108 | ENSMUST00000133228 |
| 108 | Alas2 | X | 150506782 | 150507282 | 4.61 | 500 | Intergenic | protein_coding | -40343 | ENSMUST00000066337 | ENSMUSG00000025270 | ENSMUST00000134670 |
| 109 | Cry1 | 10 | 85212418 | 85212918 | 4.61 | 500 | Intergenic | protein_coding | -27604 | ENSMUST00000020227 | ENSMUSG00000020038 | ENSMUST00000214805 |
| 110 | Depdc1b | 13 | 108385322 | 108385822 | 4.61 | 500 | protein_coding-intron (ENSMUST00000051594, intron 9 of 10) | protein_coding | 28146 | ENSMUST00000172402 | ENSMUSG00000021697 | ENSMUST00000171178 |
| 111 | Eri1 | 8 | 35461822 | 35462322 | 4.61 | 500 | Intergenic | protein_coding | 19336 | ENSMUST00000211536 | ENSMUSG00000031527 | ENSMUST00000033927 |
| 112 | Fam111a | 19 | 12544961 | 12545461 | 4.61 | 500 | protein_coding-promoter-TSS (ENSMUST00000139270) | protein_coding | -529 | ENSMUST00000139270 | ENSMUSG00000024691 | ENSMUST00000144662 |
| 113 | H2-Ob | 17 | 34230825 | 34231325 | 4.61 | 500 | Intergenic | protein_coding | -7828 | ENSMUST00000095342 | ENSMUSG00000041538 | ENSMUST00000173764 |
| 114 | Palld | 8 | 61737347 | 61737847 | 4.61 | 500 | protein_coding-intron (ENSMUST00000034057, intron 2 of 19) | protein_coding | 22492 | ENSMUST00000121493 | ENSMUSG00000058056 | ENSMUST00000034057 |
| 115 | Paxip1 | 5 | 27691848 | 27692348 | 4.61 | 500 | protein_coding-intron (ENSMUST00000071500, intron 16 of 25) | protein_coding | 59197 | ENSMUST00000197625 | ENSMUSG00000002221 | ENSMUST00000002291 |
| 116 | Bhlhb9 | X | 135813311 | 135813811 | 4.59 | 500 | protein_coding-intron (ENSMUST00000207433, intron 1 of 10) | protein_coding | 15967 | ENSMUST00000207433 | ENSMUSG00000072964 | ENSMUST00000180025 |
| 117 | Megf9 | 4 | 70645094 | 70645594 | 4.59 | 500 | Intergenic | protein_coding | -110349 | ENSMUST00000037881 | ENSMUSG00000039270 | ENSMUST00000037881 |
| 118 | Scfd1 | 12 | 51455555 | 51456055 | 4.59 | 500 | Intergenic | protein_coding | 32875 | ENSMUST00000218138 | ENSMUSG00000020952 | ENSMUST00000219434 |
| 119 | Csrp2 | 10 | 110916006 | 110916506 | 4.59 | 500 | Intergenic | protein_coding | -3533 | ENSMUST00000020403 | ENSMUSG00000020186 | ENSMUST00000219188 |
| 120 | Dis3 | 14 | 99081390 | 99081890 | 4.59 | 500 | protein_coding-intron (ENSMUST00000042471, intron 15 of 20) | protein_coding | -2957 | ENSMUST00000228756 | ENSMUSG00000033166 | ENSMUST00000228643 |
| 121 | Haus8 | 8 | 71288270 | 71288770 | 4.59 | 500 | protein_coding-intron (ENSMUST00000071935, intron 1 of 39) | protein_coding | -15586 | ENSMUST00000035960 | ENSMUSG00000035439 | ENSMUST00000157039 |
| 122 | Pdk3 | X | 93828903 | 93829403 | 4.59 | 500 | protein_coding-intron (ENSMUST00000045748, intron 1 of 11) | protein_coding | 3048 | ENSMUST00000045748 | ENSMUSG00000035232 | ENSMUST00000154717 |
| 123 | Rbbp7 | X | 162791250 | 162791750 | 4.59 | 500 | protein_coding-intron (ENSMUST00000041370, intron 4 of 9) | protein_coding | 16922 | ENSMUST00000143681 | ENSMUSG00000031353 | ENSMUST00000138791 |
| 124 | Dsp | 13 | 38190887 | 38191387 | 4.54 | 500 | protein_coding-intron (ENSMUST00000124830, intron 22 of 23) | protein_coding | 12770 | ENSMUST00000145151 | ENSMUSG00000054889 | ENSMUST00000145151 |
| 125 | Mctp2 | 7 | 72447735 | 72448235 | 4.54 | 500 | Intergenic | protein_coding | -141377 | ENSMUST00000206755 | ENSMUSG00000032776 | ENSMUST00000205410 |
| 126 | Phf6 | X | 52917227 | 52917727 | 4.54 | 500 | protein_coding-intron (ENSMUST00000078944, intron 3 of 10) | protein_coding | 5182 | ENSMUST00000154864 | ENSMUSG00000025626 | ENSMUST00000179014 |
| 127 | Rabgap1l | 1 | 160511423 | 160511923 | 4.54 | 500 | protein_coding-intron (ENSMUST00000028049, intron 13 of 20) | protein_coding | -57900 | ENSMUST00000194696 | ENSMUSG00000026721 | ENSMUST00000195114 |
| 128 | Arl6 | 16 | 59739235 | 59739735 | 4.52 | 500 | protein_coding-intron (ENSMUST00000068860, intron 14 of 17) | protein_coding | -100094 | ENSMUST00000099646 | ENSMUSG00000022722 | ENSMUST00000099646 |
| 129 | Cox7c | 13 | 86066671 | 86067171 | 4.52 | 500 | Intergenic | protein_coding | -20017 | ENSMUST00000131011 | ENSMUSG00000017778 | ENSMUST00000017922 |
| 130 | Magi3 | 3 | 104230398 | 104230898 | 4.52 | 500 | Intergenic | protein_coding | -10274 | ENSMUST00000064371 | ENSMUSG00000052539 | ENSMUST00000199201 |
| 131 | 1110059E24Rik | 19 | 21592377 | 21592877 | 4.52 | 500 | protein_coding-intron (ENSMUST00000177577, intron 1 of 2) | protein_coding | 11692 | ENSMUST00000177605 | ENSMUSG00000035171 | ENSMUST00000177577 |
| 132 | Arhgef12 | 9 | 43057777 | 43058277 | 4.52 | 500 | protein_coding-intron (ENSMUST00000072767, intron 1 of 41) | protein_coding | -15395 | ENSMUST00000213566 | ENSMUSG00000059495 | ENSMUST00000072767 |
| 133 | Ccdc138 | 10 | 58494720 | 58495220 | 4.52 | 500 | protein_coding-TTS (ENSMUST00000003310) | protein_coding | -2978 | ENSMUST00000036576 | ENSMUSG00000038010 | ENSMUST00000036576 |
| 134 | Cd83 | 13 | 43812588 | 43813088 | 4.52 | 500 | Intergenic | protein_coding | 27653 | ENSMUST00000221359 | ENSMUSG00000015396 | ENSMUST00000221359 |
| 135 | Dusp10 | 1 | 183987018 | 183987518 | 4.52 | 500 | Intergenic | protein_coding | -26034 | ENSMUST00000139839 | ENSMUSG00000039384 | ENSMUST00000139839 |
| 136 | Gipc1 | 8 | 83648531 | 83649031 | 4.52 | 500 | Intergenic | protein_coding | -3896 | ENSMUST00000019577 | ENSMUSG00000019433 | ENSMUST00000211985 |
| 137 | Gpr34 | X | 13625268 | 13625768 | 4.52 | 500 | protein_coding-TTS (ENSMUST00000124710) | protein_coding | -6571 | ENSMUST00000041708 | ENSMUSG00000040229 | ENSMUST00000096492 |
| 138 | Luzp1 | 4 | 136529361 | 136529861 | 4.52 | 500 | protein_coding-intron (ENSMUST00000063021, intron 1 of 3) | protein_coding | 8216 | ENSMUST00000063021 | ENSMUSG00000001089 | ENSMUST00000105849 |
| 139 | Ncam1 | 9 | 49781036 | 49781536 | 4.52 | 500 | protein_coding-intron (ENSMUST00000114476, intron 1 of 15) | protein_coding | 17472 | ENSMUST00000192584 | ENSMUSG00000039542 | ENSMUST00000217121 |
| 140 | Ndufs4 | 13 | 114338928 | 114339428 | 4.52 | 500 | protein_coding-intron (ENSMUST00000022286, intron 2 of 4) | protein_coding | 48871 | ENSMUST00000225707 | ENSMUSG00000021764 | ENSMUST00000232101 |
| 141 | Snx25 | 8 | 46114672 | 46115172 | 4.52 | 500 | protein_coding-intron (ENSMUST00000041582, intron 3 of 18) | protein_coding | 1780 | ENSMUST00000176410 | ENSMUSG00000038291 | ENSMUST00000144244 |
| 142 | Zfp521 | 18 | 14013016 | 14013516 | 4.52 | 500 | Intergenic | protein_coding | -40479 | ENSMUST00000025288 | ENSMUSG00000024420 | ENSMUST00000234025 |
| 143 | App | 16 | 85341902 | 85342402 | 4.47 | 500 | Intergenic | protein_coding | -168386 | ENSMUST00000227723 | ENSMUSG00000022892 | ENSMUST00000228375 |
| 144 | Atg10 | 13 | 91071871 | 91072371 | 4.47 | 500 | protein_coding-intron (ENSMUST00000022119, intron 3 of 7) | protein_coding | 53489 | ENSMUST00000223729 | ENSMUSG00000021619 | ENSMUST00000224449 |
| 145 | Chd9 | 8 | 91013046 | 91013546 | 4.47 | 500 | protein_coding-intron (ENSMUST00000048665, intron 23 of 39) | protein_coding | 21733 | ENSMUST00000211552 | ENSMUSG00000056608 | ENSMUST00000209474 |
| 146 | Hmgn2 | 4 | 133946972 | 133947472 | 4.47 | 500 | Intergenic | protein_coding | 20021 | ENSMUST00000123234 | ENSMUSG00000003038 | ENSMUST00000100472 |
| 147 | Ncald | 15 | 37808411 | 37808911 | 4.47 | 500 | Intergenic | protein_coding | -16091 | ENSMUST00000150453 | ENSMUSG00000051359 | ENSMUST00000137944 |
| 148 | Prune2 | 19 | 17180394 | 17180894 | 4.47 | 500 | protein_coding-intron (ENSMUST00000087689, intron 11 of 19) | protein_coding | -27607 | ENSMUST00000225838 | ENSMUSG00000039126 | ENSMUST00000224059 |
| 149 | Thg1l | 11 | 45963272 | 45963772 | 4.47 | 500 | Intergenic | protein_coding | -8028 | ENSMUST00000011398 | ENSMUSG00000011254 | ENSMUST00000133238 |
| 150 | Ttk | 9 | 83838161 | 83838661 | 4.47 | 500 | protein_coding-TTS (ENSMUST00000189727) | protein_coding | 3645 | ENSMUST00000191484 | ENSMUSG00000038379 | ENSMUST00000186277 |
| 151 | Pde4d | 13 | 108977144 | 108977644 | 4.46 | 500 | protein_coding-intron (ENSMUST00000122041, intron 2 of 16) | protein_coding | 117228 | ENSMUST00000177907 | ENSMUSG00000021699 | ENSMUST00000117879 |
| 152 | Ccdc18 | 5 | 108172196 | 108172696 | 4.46 | 500 | protein_coding-intron (ENSMUST00000047677, intron 12 of 28) | protein_coding | 10767 | ENSMUST00000195973 | ENSMUSG00000056531 | ENSMUST00000047677 |
| 153 | Pds5b | 5 | 150735977 | 150736477 | 4.46 | 500 | protein_coding-intron (ENSMUST00000016569, intron 7 of 34) | protein_coding | -11595 | ENSMUST00000133738 | ENSMUSG00000034021 | ENSMUST00000038900 |
| 154 | Skap2 | 6 | 51818179 | 51818679 | 4.46 | 500 | Intergenic | protein_coding | 53500 | ENSMUST00000205174 | ENSMUSG00000059182 | ENSMUST00000078214 |
| 155 | Ufsp2 | 8 | 45986990 | 45987490 | 4.46 | 500 | protein_coding-intron (ENSMUST00000034051, intron 7 of 11) | protein_coding | -5385 | ENSMUST00000210608 | ENSMUSG00000031634 | ENSMUST00000210081 |
| 156 | Mitf | 6 | 98005283 | 98005783 | 4.4 | 500 | protein_coding-intron (ENSMUST00000043628, intron 5 of 8) | protein_coding | 13612 | ENSMUST00000203938 | ENSMUSG00000035158 | ENSMUST00000203884 |
| 157 | Pola1 | X | 93466986 | 93467486 | 4.4 | 500 | protein_coding-intron (ENSMUST00000006856, intron 32 of 36) | protein_coding | 23710 | ENSMUST00000142479 | ENSMUSG00000006678 | ENSMUST00000006856 |
| 158 | Rpl36al | 12 | 69177448 | 69177948 | 4.4 | 500 | protein_coding-intron (ENSMUST00000110621, intron 3 of 3) | protein_coding | 6288 | ENSMUST00000110620 | ENSMUSG00000049751 | ENSMUST00000110620 |
| 159 | Tnfaip3 | 10 | 18989678 | 18990178 | 4.39 | 500 | Intergenic | protein_coding | 18917 | ENSMUST00000154749 | ENSMUSG00000019850 | ENSMUST00000154749 |
| 160 | Lzts1 | 8 | 69165577 | 69166077 | 4.39 | 500 | protein_coding-TTS (ENSMUST00000181890) | protein_coding | 16403 | ENSMUST00000209353 | ENSMUSG00000036306 | ENSMUST00000185176 |
| 161 | Mpp6 | 6 | 50196265 | 50196765 | 4.39 | 500 | protein_coding-TTS (ENSMUST00000167063) | protein_coding | 2099 | ENSMUST00000167319 | ENSMUSG00000038388 | ENSMUST00000203415 |
| 162 | Rhoh | 5 | 65874863 | 65875363 | 4.33 | 500 | protein_coding-intron (ENSMUST00000031106, intron 2 of 3) | protein_coding | 11495 | ENSMUST00000202471 | ENSMUSG00000029204 | ENSMUST00000202591 |
| 163 | Sacs | 14 | 61159952 | 61160452 | 4.33 | 500 | protein_coding-intron (ENSMUST00000089394, intron 2 of 9) | protein_coding | 12150 | ENSMUST00000119943 | ENSMUSG00000048279 | ENSMUST00000119943 |
| 164 | Casp4 | 9 | 5326956 | 5327456 | 4.32 | 500 | protein_coding-TTS (ENSMUST00000163086) | protein_coding | 3858 | ENSMUST00000163086 | ENSMUSG00000033538 | ENSMUST00000160064 |
| 165 | Asb4 | 6 | 5419843 | 5420343 | 4.32 | 500 | protein_coding-intron (ENSMUST00000043294, intron 3 of 5) | protein_coding | 29666 | ENSMUST00000183358 | ENSMUSG00000042607 | ENSMUST00000043294 |
| 166 | Cd59b | 2 | 104073012 | 104073512 | 4.32 | 500 | protein_coding-intron (ENSMUST00000090429, intron 1 of 3) | protein_coding | 2196 | ENSMUST00000111130 | ENSMUSG00000068686 | ENSMUST00000129749 |
| 167 | Pnpt1 | 11 | 29057359 | 29057859 | 4.32 | 500 | Intergenic | protein_coding | -73135 | ENSMUST00000154924 | ENSMUSG00000020464 | ENSMUST00000131527 |
| 168 | Cep57 | 9 | 13814739 | 13815239 | 4.26 | 500 | protein_coding-intron (ENSMUST00000034398, intron 6 of 10) | protein_coding | 1408 | ENSMUST00000124524 | ENSMUSG00000031922 | ENSMUST00000150893 |
| 169 | Mipep | 14 | 60850229 | 60850729 | 4.26 | 500 | protein_coding-intron (ENSMUST00000063562, intron 16 of 18) | protein_coding | 65841 | ENSMUST00000223709 | ENSMUSG00000021993 | ENSMUST00000224635 |
| 170 | Zdhhc21 | 4 | 82851341 | 82851841 | 4.26 | 500 | protein_coding-promoter-TSS (ENSMUST00000139401) | protein_coding | -777 | ENSMUST00000139401 | ENSMUSG00000028403 | ENSMUST00000030110 |
| 171 | Slc35f5 | 1 | 125504045 | 125504545 | 4.19 | 500 | Intergenic | protein_coding | -56300 | ENSMUST00000190037 | ENSMUSG00000026342 | ENSMUST00000027580 |
| 172 | Cenpi | X | 134327813 | 134328313 | 4.06 | 500 | protein_coding-intron (ENSMUST00000081064, intron 9 of 22) | protein_coding | 16065 | ENSMUST00000144678 | ENSMUSG00000031262 | ENSMUST00000081064 |

H3K4me1 - enhancer

| | Gene.Name | Chr | Start | End | Peak.Score | Focus.Ratio.Region.Size | Detailed.Annotation | Gene.Type | Distance.to.TSS | Nearest.PromoterID | Entrez.ID | Nearest.Unigene |
|---|---|---|---|---|---|---|---|---|---|---|---|---|
| 1 | Gfod1 | 13 | 43296629 | 43296951 | 4.93 | 0.681 | protein_coding-intron (ENSMUST00000055341, intron 1 of 1) | protein_coding | 6615 | ENSMUST00000222779 | ENSMUSG00000051335 | ENSMUST00000222779 |
| 2 | Baiap2 | 11 | 119960077 | 119960513 | 4.79 | 1.474 | protein_coding-TTS (ENSMUST00000146960) | protein_coding | 274 | ENSMUST00000146960 | ENSMUSG00000025372 | ENSMUST00000146566 |
| 3 | Lrp8 | 4 | 107806338 | 107806802 | 4.73 | 1.385 | lncRNA-exon (ENSMUST00000097930, exon 2 of 2) | protein_coding | 3702 | ENSMUST00000146552 | ENSMUSG00000028613 | ENSMUST00000123140 |
| 4 | Lhfpl2 | 13 | 94094786 | 94095108 | 4.65 | 1.234 | protein_coding-intron (ENSMUST00000054274, intron 1 of 3) | protein_coding | 486 | ENSMUST00000221096 | ENSMUSG00000045312 | ENSMUST00000223423 |
| 5 | Frmd4b | 6 | 97278309 | 97278711 | 4.61 | 0.693 | Intergenic | protein_coding | 12751 | ENSMUST00000203486 | ENSMUSG00000030064 | ENSMUST00000113353 |
| 6 | Uba7 | 9 | 107974608 | 107974930 | 4.61 | 0.705 | protein_coding-promoter-TSS (ENSMUST00000035216) | protein_coding | -736 | ENSMUST00000176037 | ENSMUSG00000032596 | ENSMUST00000176037 |
| 7 | Arsb | 13 | 93758178 | 93758664 | 4.4 | 0.718 | Intergenic | protein_coding | -13209 | ENSMUST00000091403 | ENSMUSG00000042082 | ENSMUST00000091403 |
| 8 | Muc1 | 3 | 89231740 | 89232235 | 4.38 | 0.667 | protein_coding-TTS (ENSMUST00000146844) | protein_coding | 516 | ENSMUST00000139206 | ENSMUSG00000042784 | ENSMUST00000146844 |
| 9 | Cdc6 | 11 | 98911162 | 98911728 | 4.29 | 0.605 | protein_coding-exon (ENSMUST00000092706, exon 5 of 12) | protein_coding | 3294 | ENSMUST00000093937 | ENSMUSG00000017499 | ENSMUST00000093937 |
| 10 | Rest | 5 | 77249766 | 77250088 | 4.25 | 0.768 | Intergenic | protein_coding | -15564 | ENSMUST00000080359 | ENSMUSG00000029249 | ENSMUST00000113449 |
| 11 | Tpi1 | 6 | 124811129 | 124811671 | 4.11 | 0.567 | protein_coding-TTS (ENSMUST00000047760) | protein_coding | 486 | ENSMUST00000133251 | ENSMUSG00000023456 | ENSMUST00000133251 |
| 12 | Irf2 | 8 | 46743024 | 46743346 | 4.1 | 0.428 | protein_coding-intron (ENSMUST00000034041, intron 1 of 8) | protein_coding | 2919 | ENSMUST00000207571 | ENSMUSG00000031627 | ENSMUST00000210218 |
| 13 | Irf8 | 8 | 120806600 | 120807077 | 4.08 | 0.34 | protein_coding-intron (ENSMUST00000127664, intron 1 of 15) | protein_coding | 54629 | ENSMUST00000160594 | ENSMUSG00000041515 | ENSMUST00000162775 |
| 14 | Mpzl3 | 9 | 45063914 | 45064236 | 4.07 | 0.428 | protein_coding-exon (ENSMUST00000093856, exon 2 of 2) | protein_coding | 4826 | ENSMUST00000187113 | ENSMUSG00000070305 | ENSMUST00000114664 |
| 15 | Calm3 | 7 | 16919882 | 16920348 | 3.96 | 0.428 | protein_coding-intron (ENSMUST00000019514, intron 1 of 5) | protein_coding | -2912 | ENSMUST00000173139 | ENSMUSG00000019370 | ENSMUST00000172594 |
| 16 | Pkib | 10 | 57650050 | 57650447 | 3.94 | 0.331 | protein_coding-promoter-TSS (ENSMUST00000066028) | protein_coding | -733 | ENSMUST00000066028 | ENSMUSG00000019876 | ENSMUST00000176225 |
| 17 | Tsen15 | 1 | 152377253 | 152377739 | 3.91 | 0.529 | protein_coding-intron (ENSMUST00000015124, intron 3 of 4) | protein_coding | -5414 | ENSMUST00000161717 | ENSMUSG00000014980 | ENSMUST00000159270 |
| 18 | Ttc3 | 16 | 94453125 | 94453522 | 3.87 | 0.282 | protein_coding-intron (ENSMUST00000117648, intron 37 of 45) | protein_coding | 2197 | ENSMUST00000141192 | ENSMUSG00000040785 | ENSMUST00000129536 |
| 19 | Arhgef7 | 8 | 11774357 | 11774764 | 3.84 | 0.365 | protein_coding-intron (ENSMUST00000033908, intron 2 of 17) | protein_coding | -7967 | ENSMUST00000211409 | ENSMUSG00000031511 | ENSMUST00000098938 |
| 20 | Socs1 | 16 | 10774752 | 10775154 | 3.82 | 0.453 | Intergenic | protein_coding | 10583 | ENSMUST00000229866 | ENSMUSG00000038037 | ENSMUST00000038099 |
| 21 | Ankrd33b | 15 | 31332435 | 31332757 | 3.79 | 0.327 | protein_coding-intron (ENSMUST00000044324, intron 1 of 2) | protein_coding | -7347 | ENSMUST00000227391 | ENSMUSG00000022237 | ENSMUST00000110410 |
| 22 | Dtx4 | 19 | 12501008 | 12501421 | 3.77 | 0.29 | protein_coding-exon (ENSMUST00000045521, exon 1 of 9) | protein_coding | 240 | ENSMUST00000045521 | ENSMUSG00000039982 | ENSMUST00000045521 |
| 23 | Susd1 | 4 | 59355681 | 59356003 | 3.76 | 0.239 | protein_coding-intron (ENSMUST00000040166, intron 10 of 16) | protein_coding | 7140 | ENSMUST00000136077 | ENSMUSG00000038578 | ENSMUST00000136562 |
| 24 | 4930438A08Rik | 11 | 58289533 | 58289939 | 3.71 | 0.277 | protein_coding-exon (ENSMUST00000108834, exon 5 of 7) | protein_coding | 14676 | ENSMUST00000208022 | ENSMUSG00000069873 | ENSMUST00000208022 |
| 25 | Ifih1 | 2 | 62647211 | 62647663 | 3.66 | 0.214 | Intergenic | protein_coding | -1182 | ENSMUST00000176431 | ENSMUSG00000026896 | ENSMUST00000028259 |
| 26 | Itpa | 2 | 130669565 | 130669979 | 3.65 | 0.315 | protein_coding-intron (ENSMUST00000103193, intron 1 of 7) | protein_coding | 1907 | ENSMUST00000126714 | ENSMUSG00000074797 | ENSMUST00000103193 |
| 27 | Gpx3 | 11 | 54903578 | 54903900 | 3.64 | 0.29 | protein_coding-intron (ENSMUST00000082430, intron 1 of 4) | protein_coding | 813 | ENSMUST00000149324 | ENSMUSG00000018339 | ENSMUST00000149324 |
| 28 | Mmp28 | 11 | 83466996 | 83467318 | 3.64 | 0.302 | Intergenic | protein_coding | -4086 | ENSMUST00000108137 | ENSMUSG00000020682 | ENSMUST00000119346 |
| 29 | Rad18 | 6 | 112692147 | 112692544 | 3.63 | 0.127 | protein_coding-intron (ENSMUST00000068487, intron 2 of 13) | protein_coding | 4259 | ENSMUST00000113182 | ENSMUSG00000030254 | ENSMUST00000142079 |
| 30 | Slc7a8 | 14 | 54760594 | 54761024 | 3.63 | 0.277 | protein_coding-intron (ENSMUST00000022787, intron 1 of 10) | protein_coding | 21121 | ENSMUST00000022787 | ENSMUSG00000022180 | ENSMUST00000022787 |
| 31 | Ubl5 | 9 | 20643542 | 20643864 | 3.57 | 0.164 | protein_coding-promoter-TSS (ENSMUST00000086459) | protein_coding | -368 | ENSMUST00000160682 | ENSMUSG00000084786 | ENSMUST00000161887 |
| 32 | Uhrf1 | 17 | 56310169 | 56310583 | 3.57 | 0.114 | protein_coding-intron (ENSMUST00000001258, intron 3 of 16) | protein_coding | -2287 | ENSMUST00000137876 | ENSMUSG00000001228 | ENSMUST00000001258 |
| 33 | Lin9 | 1 | 180662786 | 180663183 | 3.56 | 0.07 | protein_coding-intron (ENSMUST00000085804, intron 6 of 11) | protein_coding | 1507 | ENSMUST00000194638 | ENSMUSG00000058729 | ENSMUST00000193892 |
| 34 | Nrgn | 9 | 37544010 | 37544412 | 3.56 | 0.227 | protein_coding-TTS (ENSMUST00000002008) | protein_coding | 1927 | ENSMUST00000182070 | ENSMUSG00000053310 | ENSMUST00000182070 |
| 35 | Tcp11l1 | 2 | 104691526 | 104691848 | 3.56 | 0.177 | protein_coding-intron (ENSMUST00000028597, intron 5 of 9) | protein_coding | -7421 | ENSMUST00000129571 | ENSMUSG00000027175 | ENSMUST00000137428 |
| 36 | Dmkn | 7 | 30781928 | 30782353 | 3.55 | 0.188 | Intergenic | protein_coding | 5739 | ENSMUST00000041703 | ENSMUSG00000060962 | ENSMUST00000054427 |
| 37 | Rps6ka4 | 19 | 6840715 | 6841037 | 3.55 | 0.151 | protein_coding-promoter-TSS (ENSMUST00000025903) | protein_coding | -240 | ENSMUST00000237442 | ENSMUSG00000024952 | ENSMUST00000236334 |
| 38 | Copz2 | 11 | 96860657 | 96860979 | 3.54 | 0.151 | protein_coding-intron (ENSMUST00000018816, intron 8 of 8) | protein_coding | 6687 | ENSMUST00000147710 | ENSMUSG00000018672 | ENSMUST00000155696 |
| 39 | Golgb1 | 16 | 36918861 | 36919183 | 3.54 | 0.063 | protein_coding-exon (ENSMUST00000039855, exon 13 of 21) | protein_coding | -5435 | ENSMUST00000152864 | ENSMUSG00000034243 | ENSMUST00000140266 |
| 40 | Tlr2 | 3 | 83841216 | 83841715 | 3.54 | 0.101 | protein_coding-intron (ENSMUST00000029623, intron 1 of 2) | protein_coding | 302 | ENSMUST00000029623 | ENSMUSG00000027995 | ENSMUST00000029623 |
| 41 | Cdc42bpa | 1 | 180119173 | 180119570 | 3.53 | 0.078 | protein_coding-TTS (ENSMUST00000152582) | protein_coding | -4395 | ENSMUST00000145274 | ENSMUSG00000026490 | ENSMUST00000135056 |
| 42 | Mapkapk3 | 9 | 107259736 | 107260143 | 3.53 | 0.202 | protein_coding-exon (ENSMUST00000155860, exon 6 of 6) | protein_coding | 1997 | ENSMUST00000141674 | ENSMUSG00000032577 | ENSMUST00000035194 |
| 43 | Psen2 | 1 | 180290810 | 180291132 | 3.51 | 0.163 | Intergenic | protein_coding | -27533 | ENSMUST00000133340 | ENSMUSG00000010609 | ENSMUST00000111105 |
| 44 | Tpm3 | 3 | 90086437 | 90086842 | 3.51 | 0.214 | protein_coding-promoter-TSS (ENSMUST00000133281) | protein_coding | 268 | ENSMUST00000149115 | ENSMUSG00000027940 | ENSMUST00000143281 |
| 45 | Slc25a24 | 3 | 109126054 | 109126376 | 3.5 | 0.227 | protein_coding-intron (ENSMUST00000029477, intron 1 of 9) | protein_coding | 3053 | ENSMUST00000140786 | ENSMUSG00000040322 | ENSMUST00000140786 |
| 46 | B230118H07Rik | 2 | 101581166 | 101581488 | 3.49 | 0.163 | protein_coding-intron (ENSMUST00000090513, intron 5 of 6) | protein_coding | 3607 | ENSMUST00000138903 | ENSMUSG00000027165 | ENSMUST00000177007 |
| 47 | Wee1 | 7 | 110128449 | 110128894 | 3.49 | 0.126 | protein_coding-intron (ENSMUST00000033326, intron 5 of 10) | protein_coding | 6625 | ENSMUST00000033326 | ENSMUSG00000031016 | ENSMUST00000033326 |
| 48 | Gramd1b | 9 | 40296887 | 40297284 | 3.47 | 0.085 | protein_coding-TTS (ENSMUST00000118159) | protein_coding | 9910 | ENSMUST00000155265 | ENSMUSG00000040111 | ENSMUST00000118159 |
| 49 | Oas1g | 5 | 120880569 | 120880980 | 3.45 | 0.164 | protein_coding-intron (ENSMUST00000086368, intron 3 of 6) | protein_coding | -1171 | ENSMUST00000161809 | ENSMUSG00000066861 | ENSMUST00000086368 |
| 50 | AB124611 | 9 | 21529837 | 21530249 | 3.44 | 0.113 | protein_coding-intron (ENSMUST00000076326, intron 3 of 7) | protein_coding | 3815 | ENSMUST00000076326 | ENSMUSG00000057191 | ENSMUST00000086361 |
| 51 | Smtn | 11 | 3535662 | 3536075 | 3.44 | 0.075 | lncRNA-TTS (ENSMUST00000125275) | protein_coding | 1530 | ENSMUST00000110011 | ENSMUSG00000020439 | ENSMUST00000136243 |
| 52 | Ttf2 | 3 | 100973242 | 100973564 | 3.41 | 0.101 | Intergenic | protein_coding | -3740 | ENSMUST00000076941 | ENSMUSG00000033222 | ENSMUST00000151697 |
| 53 | Il12rb1 | 8 | 70811061 | 70811383 | 3.4 | 0.189 | protein_coding-TTS (ENSMUST00000212146) | protein_coding | -1118 | ENSMUST00000212251 | ENSMUSG00000000791 | ENSMUST00000212146 |
| 54 | N4bp1 | 8 | 86851585 | 86851907 | 3.39 | 0.189 | protein_coding-exon (ENSMUST00000210029, exon 1 of 3) | protein_coding | 674 | ENSMUST00000210029 | ENSMUSG00000031652 | ENSMUST00000034074 |
| 55 | Pdlim4 | 11 | 54053334 | 54053738 | 3.39 | 0.189 | Intergenic | protein_coding | 11032 | ENSMUST00000151948 | ENSMUSG00000020388 | ENSMUST00000093109 |
| 56 | Lman1 | 18 | 66027161 | 66027558 | 3.38 | 0.056 | Intergenic | protein_coding | -4779 | ENSMUST00000143990 | ENSMUSG00000041891 | ENSMUST00000155895 |
| 57 | Rttn | 18 | 88990255 | 88990652 | 3.38 | 0.056 | protein_coding-TTS (ENSMUST00000235314) | protein_coding | 4559 | ENSMUST00000235314 | ENSMUSG00000023066 | ENSMUST00000236313 |
| 58 | Ndufs3 | 2 | 90896270 | 90896722 | 3.38 | 0.114 | protein_coding-intron (ENSMUST00000005647, intron 6 of 6) | protein_coding | 1958 | ENSMUST00000140248 | ENSMUSG00000005510 | ENSMUST00000152059 |
| 59 | Tmem106a | 11 | 101584198 | 101584520 | 3.38 | 0.151 | protein_coding-intron (ENSMUST00000039581, intron 3 of 8) | protein_coding | -1638 | ENSMUST00000128659 | ENSMUSG00000034947 | ENSMUST00000128614 |
| 60 | Panx1 | 9 | 15042809 | 15043131 | 3.35 | 0.189 | protein_coding-intron (ENSMUST00000056755, intron 1 of 3) | protein_coding | 827 | ENSMUST00000169288 | ENSMUSG00000031934 | ENSMUST00000056755 |
| 61 | Btbd16 | 7 | 130782741 | 130783138 | 3.33 | 0.049 | protein_coding-intron (ENSMUST00000048453, intron 3 of 15) | protein_coding | 8772 | ENSMUST00000208593 | ENSMUSG00000040298 | ENSMUST00000208593 |
| 62 | Pdcd10 | 3 | 75526782 | 75527179 | 3.32 | 0.049 | protein_coding-intron (ENSMUST00000029424, intron 5 of 7) | protein_coding | -5410 | ENSMUST00000160196 | ENSMUSG00000027835 | ENSMUST00000029424 |
| 63 | Mlec | 5 | 115167132 | 115167454 | 3.32 | 0.088 | Intergenic | protein_coding | -9114 | ENSMUST00000112121 | ENSMUSG00000048578 | ENSMUST00000124678 |
| 64 | Ezh2 | 6 | 47573363 | 47573760 | 3.31 | 0.028 | protein_coding-intron (ENSMUST00000081721, intron 3 of 19) | protein_coding | 4099 | ENSMUST00000092648 | ENSMUSG00000029687 | ENSMUST00000081721 |
| 65 | Pi4k2b | 5 | 52759368 | 52759765 | 3.3 | 0.049 | protein_coding-intron (ENSMUST00000031081, intron 7 of 9) | protein_coding | 4964 | ENSMUST00000145825 | ENSMUSG00000029186 | ENSMUST00000031081 |
| 66 | Sertad2 | 11 | 20631348 | 20631670 | 3.29 | 0.063 | protein_coding-promoter-TSS (ENSMUST00000109586) | protein_coding | -470 | ENSMUST00000109586 | ENSMUSG00000049800 | ENSMUST00000109585 |
| 67 | Slco4a1 | 2 | 180456237 | 180456646 | 3.29 | 0.063 | protein_coding-intron (ENSMUST00000038259, intron 1 of 11) | protein_coding | 107 | ENSMUST00000139902 | ENSMUSG00000038963 | ENSMUST00000128367 |
| 68 | Ube2c | 2 | 164773254 | 164773576 | 3.29 | 0.05 | protein_coding-TTS (ENSMUST00000088248) | protein_coding | 3512 | ENSMUST00000001439 | ENSMUSG00000001403 | ENSMUST00000001439 |
| 69 | Coro7 | 16 | 4666712 | 4667120 | 3.28 | 0.202 | protein_coding-intron (ENSMUST00000038552, intron 6 of 27) | protein_coding | -4994 | ENSMUST00000130125 | ENSMUSG00000039637 | ENSMUST00000143723 |
| 70 | Nfkbia | 12 | 55457054 | 55457376 | 3.28 | 0.177 | Intergenic | protein_coding | 34595 | ENSMUST00000219869 | ENSMUSG00000021025 | ENSMUST00000219869 |
| 71 | Rhou | 8 | 123663607 | 123664058 | 3.27 | 0.189 | protein_coding-TTS (ENSMUST00000045487) | protein_coding | 9903 | ENSMUST00000045487 | ENSMUSG00000039960 | ENSMUST00000136615 |
| 72 | Rhobtb1 | 10 | 69213593 | 69213915 | 3.26 | 0.113 | protein_coding-promoter-TSS (ENSMUST00000170048) | protein_coding | 560 | ENSMUST00000067908 | ENSMUSG00000019944 | ENSMUST00000164876 |
| 73 | Adam3 | 8 | 24689776 | 24690173 | 3.25 | 0.056 | protein_coding-intron (ENSMUST00000033958, intron 16 of 21) | protein_coding | -2011 | ENSMUST00000167431 | ENSMUSG00000031553 | ENSMUST00000167703 |
| 74 | Syne1 | 10 | 5490634 | 5491031 | 3.25 | 0.056 | protein_coding-intron (ENSMUST00000041639, intron 2 of 31) | protein_coding | 59846 | ENSMUST00000215295 | ENSMUSG00000096054 | ENSMUST00000215467 |
| 75 | Asf1b | 8 | 83967756 | 83968078 | 3.25 | 0.101 | protein_coding-TTS (ENSMUST00000124913) | protein_coding | 2870 | ENSMUST00000124174 | ENSMUSG00000005470 | ENSMUST00000005607 |
| 76 | Dctd | 8 | 48111357 | 48111754 | 3.24 | 0.028 | protein_coding-intron (ENSMUST00000033966, intron 1 of 5) | protein_coding | 1344 | ENSMUST00000174379 | ENSMUSG00000031562 | ENSMUST00000174379 |
| 77 | Cdca7 | 2 | 72509349 | 72509751 | 3.23 | 0.151 | Intergenic | protein_coding | 24443 | ENSMUST00000137880 | ENSMUSG00000055612 | ENSMUST00000127344 |
| 78 | Dgcr6 | 16 | 18065495 | 18065898 | 3.23 | 0.138 | protein_coding-promoter-TSS (ENSMUST00000066027) | protein_coding | 537 | ENSMUST00000076757 | ENSMUSG00000003531 | ENSMUST00000143343 |
| 79 | Shb | 4 | 45529718 | 45530040 | 3.23 | 0.151 | protein_coding-exon (ENSMUST00000061986, exon 1 of 6) | protein_coding | 451 | ENSMUST00000061986 | ENSMUSG00000044813 | ENSMUST00000146236 |
| 80 | Sco1 | 11 | 67055295 | 67055692 | 3.22 | 0.07 | protein_coding-intron (ENSMUST00000092996, intron 2 of 5) | protein_coding | 1718 | ENSMUST00000136984 | ENSMUSG00000069844 | ENSMUST00000108690 |
| 81 | Tnr | 1 | 159790848 | 159791245 | 3.22 | 0.056 | protein_coding-intron (ENSMUST00000111669, intron 1 of 21) | protein_coding | 17276 | ENSMUST00000193325 | ENSMUSG00000015829 | ENSMUST00000192457 |
| 82 | Gnptab | 10 | 88422004 | 88422326 | 3.21 | 0.101 | protein_coding-intron (ENSMUST00000020251, intron 7 of 20) | protein_coding | -7401 | ENSMUST00000132738 | ENSMUSG00000035311 | ENSMUST00000130301 |
| 83 | Arf3 | 15 | 98763670 | 98763992 | 3.2 | 0.101 | protein_coding-promoter-TSS (ENSMUST00000053183) | protein_coding | -617 | ENSMUST00000129354 | ENSMUSG00000051853 | ENSMUST00000129354 |
| 84 | Insig1 | 5 | 28118594 | 28118916 | 3.18 | 0.114 | Intergenic | protein_coding | 44089 | ENSMUST00000199728 | ENSMUSG00000045294 | ENSMUST00000059155 |
| 85 | S1pr2 | 9 | 20972437 | 20972759 | 3.18 | 0.151 | protein_coding-intron (ENSMUST00000054197, intron 1 of 1) | protein_coding | 1380 | ENSMUST00000216499 | ENSMUSG00000043895 | ENSMUST00000214333 |
| 86 | Snd1 | 6 | 28667801 | 28668198 | 3.17 | 0.035 | protein_coding-intron (ENSMUST00000001460, intron 11 of 23) | protein_coding | -36910 | ENSMUST00000163772 | ENSMUSG00000001424 | ENSMUST00000165151 |
| 87 | AI413582 | 17 | 27566378 | 27566700 | 3.17 | 0.126 | protein_coding-promoter-TSS (ENSMUST00000080190) | protein_coding | -783 | ENSMUST00000154473 | ENSMUSG00000062753 | ENSMUST00000145183 |
| 88 | Nos2 | 11 | 78920960 | 78921282 | 3.17 | 0.075 | protein_coding-intron (ENSMUST00000018610, intron 1 of 26) | protein_coding | 334 | ENSMUST00000018610 | ENSMUSG00000020826 | ENSMUST00000214397 |
| 89 | Celsr1 | 15 | 86034489 | 86034811 | 3.15 | 0.101 | protein_coding-promoter-TSS (ENSMUST00000016172) | protein_coding | -873 | ENSMUST00000016172 | ENSMUSG00000016028 | ENSMUST00000227450 |
| 90 | Cflar | 1 | 58748455 | 58748860 | 3.15 | 0.101 | protein_coding-intron (ENSMUST00000069333, intron 8 of 9) | protein_coding | 18105 | ENSMUST00000123032 | ENSMUSG00000026031 | ENSMUST00000156870 |
| 91 | Dlg2 | 7 | 92191159 | 92191556 | 3.14 | 0.007 | protein_coding-intron (ENSMUST00000074273, intron 13 of 22) | protein_coding | 10576 | ENSMUST00000207798 | ENSMUSG00000052572 | ENSMUST00000239136 |
| 92 | Snx25 | 8 | 46150095 | 46150417 | 3.14 | 0.075 | protein_coding-intron (ENSMUST00000041582, intron 1 of 18) | protein_coding | 1449 | ENSMUST00000041582 | ENSMUSG00000038291 | ENSMUST00000144244 |
| 93 | Oas3 | 5 | 120759056 | 120759453 | 3.12 | 0.028 | protein_coding-intron (ENSMUST00000044833, intron 11 of 15) | protein_coding | -1599 | ENSMUST00000200795 | ENSMUSG00000032661 | ENSMUST00000044833 |
| 94 | Slc31a2 | 4 | 62259659 | 62260056 | 3.12 | 0.028 | Intergenic | protein_coding | -2705 | ENSMUST00000107468 | ENSMUSG00000066152 | ENSMUST00000107467 |
| 95 | Snrnp35 | 5 | 124490437 | 124490841 | 3.08 | 0.101 | protein_coding-TTS (ENSMUST00000136567) | protein_coding | 7181 | ENSMUST00000111453 | ENSMUSG00000029402 | ENSMUST00000111453 |
| 96 | Igf2bp3 | 6 | 49093118 | 49093515 | 3.07 | 0.049 | protein_coding-exon (ENSMUST00000203544, exon 10 of 10) | protein_coding | 895 | ENSMUST00000205057 | ENSMUSG00000029814 | ENSMUST00000205057 |
| 97 | Atm | 9 | 53499063 | 53499460 | 3.06 | 0.056 | protein_coding-intron (ENSMUST00000118282, intron 19 of 61) | protein_coding | 34705 | ENSMUST00000118282 | ENSMUSG00000034218 | ENSMUST00000232179 |
| 98 | Mogat2 | 7 | 99238812 | 99239134 | 3.06 | 0.126 | protein_coding-promoter-TSS (ENSMUST00000064231) | protein_coding | -354 | ENSMUST00000064231 | ENSMUSG00000052396 | ENSMUST00000064231 |
| 99 | Srsf3 | 17 | 29044664 | 29045067 | 3.06 | 0.126 | Intergenic | protein_coding | 5171 | ENSMUST00000233105 | ENSMUSG00000071172 | ENSMUST00000233885 |
| 100 | Clic5 | 17 | 44173044 | 44173447 | 3.05 | 0.063 | protein_coding-intron (ENSMUST00000239215, intron 1 of 5) | protein_coding | -15339 | ENSMUST00000024755 | ENSMUSG00000023959 | ENSMUST00000239215 |
| 101 | Kctd1 | 18 | 15048028 | 15048475 | 3.04 | 0.063 | lncRNA-TTS (ENSMUST00000234445) | protein_coding | 13638 | ENSMUST00000234685 | ENSMUSG00000036225 | ENSMUST00000233993 |
| 102 | Pgap2 | 7 | 102236538 | 102236860 | 3.04 | 0.05 | protein_coding-TTS (ENSMUST00000126914) | protein_coding | 796 | ENSMUST00000134621 | ENSMUSG00000030990 | ENSMUST00000120119 |
| 103 | Atg10 | 13 | 91029396 | 91029793 | 3.01 | 0.056 | protein_coding-intron (ENSMUST00000022119, intron 4 of 7) | protein_coding | 96016 | ENSMUST00000223729 | ENSMUSG00000021619 | ENSMUST00000223729 |
| 104 | Gbp4 | 5 | 105152445 | 105152842 | 3.01 | 0.056 | Intergenic | protein_coding | -13057 | ENSMUST00000100962 | ENSMUSG00000079363 | ENSMUST00000196677 |
| 105 | Dhrs3 | 4 | 144929505 | 144929827 | 3.01 | 0.05 | Intergenic | protein_coding | 10787 | ENSMUST00000133265 | ENSMUSG00000066026 | ENSMUST00000128926 |

### H3K4me1 - super enhancer

| | Gene.Name | Chr | Start | End | Peak.Score | Focus.Ratio.Region.Size | Detailed.Annotation | Gene.Type | Distance.to.TSS | Nearest.PromoterID | Entrez.ID | Nearest.Unigene |
|---|---|---|---|---|---|---|---|---|---|---|---|---|
| 1 | Cars | 7 | 143593396 | 143601472 | 24.8 | 63.135 | protein_coding-promoter-TSS (ENSMUST00000154022) | protein_coding | -682 | ENSMUST00000154022 | ENSMUSG00000010755 | ENSMUST00000105909 |
| 2 | Tnnt1 | 7 | 4508146 | 4518601 | 22.3 | 58.867 | protein_coding-TTS (ENSMUST00000163560) | protein_coding | 995 | ENSMUST00000166161 | ENSMUSG00000064179 | ENSMUST00000168111 |
| 3 | Arhgef10 | 8 | 14979323 | 14989576 | 21.2 | 53.178 | protein_coding-intron (ENSMUST00000084207, intron 26 of 28) | protein_coding | -5910 | ENSMUST00000162444 | ENSMUSG00000071176 | ENSMUST00000163062 |
| 4 | Paqr4 | 17 | 23733625 | 23742277 | 15.1 | 14.039 | protein_coding-TTS (ENSMUST00000233776) | protein_coding | 2447 | ENSMUST00000024702 | ENSMUSG00000023909 | ENSMUST00000233776 |
| 5 | Capn5 | 7 | 98175229 | 98180549 | 14.6 | 10.418 | protein_coding-intron (ENSMUST00000040971, intron 1 of 12) | protein_coding | 385 | ENSMUST00000040971 | ENSMUSG00000035547 | ENSMUST00000040971 |
| 6 | Gpc1 | 1 | 92831914 | 92835167 | 14.2 | 10.515 | protein_coding-intron (ENSMUST00000045970, intron 1 of 8) | protein_coding | 1571 | ENSMUST00000191373 | ENSMUSG00000034220 | ENSMUST00000045970 |
| 7 | Pcp4l1 | 1 | 171189673 | 171196504 | 12.9 | 13.053 | protein_coding-intron (ENSMUST00000111332, intron 1 of 2) | protein_coding | 3180 | ENSMUST00000111332 | ENSMUSG00000038370 | ENSMUST00000111332 |
| 8 | Prdx5 | 19 | 6906941 | 6910730 | 11.9 | 12.997 | protein_coding-promoter-TSS (ENSMUST00000088257) | protein_coding | -534 | ENSMUST00000149261 | ENSMUSG00000024953 | ENSMUST00000025904 |
| 9 | BC051142 | 17 | 34443659 | 34449828 | 11.2 | 11.581 | protein_coding-intron (ENSMUST00000078615, intron 11 of 19) | protein_coding | -1577 | ENSMUST00000097348 | ENSMUSG00000057246 | ENSMUST00000097348 |
| 10 | Pomp | 5 | 147840323 | 147845124 | 9.47 | 7.233 | Intergenic | protein_coding | -17738 | ENSMUST00000031654 | ENSMUSG00000029649 | ENSMUST00000202621 |
| 11 | Naa40 | 19 | 7229653 | 7235972 | 8.95 | 8.354 | protein_coding-intron (ENSMUST00000025675, intron 3 of 7) | protein_coding | 8222 | ENSMUST00000236628 | ENSMUSG00000024764 | ENSMUST00000236769 |
| 12 | Zbtb8os | 4 | 129345204 | 129346571 | 7.1 | 6.306 | protein_coding-intron (ENSMUST00000106047, intron 5 of 5) | protein_coding | 1947 | ENSMUST00000140272 | ENSMUSG00000057572 | ENSMUST00000119480 |
| 13 | Xrcc3 | 12 | 111810652 | 111811537 | 6.63 | 3.213 | protein_coding-TTS (ENSMUST00000133679) | protein_coding | 1107 | ENSMUST00000127281 | ENSMUSG00000021287 | ENSMUST00000127281 |
| 14 | Hic2 | 16 | 17238524 | 17248631 | 6.47 | 3.505 | protein_coding-intron (ENSMUST00000090190, intron 1 of 2) | protein_coding | 9913 | ENSMUST00000232082 | ENSMUSG00000050240 | ENSMUST00000090190 |
| 15 | Ctdspl | 9 | 119036634 | 119037373 | 5.81 | 4.889 | protein_coding-intron (ENSMUST00000073109, intron 6 of 7) | protein_coding | 4002 | ENSMUST00000174132 | ENSMUSG00000047409 | ENSMUST00000073109 |
| 16 | Mlh1 | 9 | 111250318 | 111250999 | 5.61 | 34.015 | protein_coding-intron (ENSMUST00000035079, intron 9 of 18) | protein_coding | -2913 | ENSMUST00000123713 | ENSMUSG00000032498 | ENSMUST00000123869 |
| 17 | Schip1 | 3 | 68494894 | 68495617 | 5.55 | 3.249 | protein_coding-exon (ENSMUST00000029346, exon 2 of 8) | protein_coding | 1047 | ENSMUST00000182719 | ENSMUSG00000027777 | ENSMUST00000170788 |
| 18 | Cbx1 | 11 | 96791581 | 96792508 | 5.36 | 2.582 | protein_coding-intron (ENSMUST00000018810, intron 1 of 3) | protein_coding | 2795 | ENSMUST00000079702 | ENSMUSG00000018666 | ENSMUST00000079702 |
| 19 | Ifih1 | 2 | 62620013 | 62620525 | 4.94 | 2.561 | protein_coding-intron (ENSMUST00000028259, intron 4 of 15) | protein_coding | 19204 | ENSMUST00000176388 | ENSMUSG00000026896 | ENSMUST00000176388 |
