## Additional_File_1 for "Inducible mechanisms of disease tolerance provide an alternative strategy of acquired immunity to malaria"

| | progenitors [bone marrow & spleen] |
|---|---|
| BV421 | **CD71** |
| BV605 | **Sca 1** |
| FITC | **CD34** |
| PerCP EF710 | **c Kit** |
| PE | **CD27** |
| PE Cy7 | ***CD16/32**** |
| APC [lineage] | **CD3**<br>**CD4**<br>**CD19**<br>**NK1.1**<br>**Ter119**<br>**CD8**<br>**Ly6G**<br>**CD11c**<br>**CD11b** |

| | red pulp Mϕ [spleen] |
|---|---|
| BV421 | **F4/80** |
| BV605 | **CD11b** |
| FITC | *auto*** |
| PerCP EF710 | **B220** |
| PE | **CD11c** |
| APC [lineage] | **CD3**<br>**CD4**<br>**CD19**<br>**NK1.1**<br>**Ter119**<br>**Ly6G** |

| | myeloid cells [bone marrow] |
|---|---|
| BV421 | **Ly6C** |
| BV605 | **CD115** |
| FITC | **Ly6G** |
| PerCP EF710 | **c Kit** |
| PE | **CD135** |
| PE Cy7 | **CD11b** |
| APC [lineage] | **CD3**<br>**CD4**<br>**CD19**<br>**NK1.1**<br>**Ter119**<br>**CD8**<br>**CD11c** |

| | iron recycling Mϕ [bone marrow] |
|---|---|
| BV421 | **F4/80** |
| BV605 | **CD115** |
| FITC | **CD11b** |
| PerCP EF710 | **VCAM-1** |
| PE | **CD11c** |
| PE Cy7 | **CD169** |
| APC [lineage] | **CD3**<br>**CD4**<br>**CD19**<br>**NK1.1**<br>**Ter119**<br>**Ly6G** |

| | myeloid cells [blood & spleen] |
|---|---|
| BV421 | **Ly6C** |
| BV605 | **CD115** |
| FITC | **IAb** |
| PerCP EF710 | **Ly6G** |
| PE | **CD11c** |
| PE Cy7 | **CD11b** |
| APC [lineage] | **CD3**<br>**CD4**<br>**CD19**<br>**NK1.1**<br>**Ter119** |

| | patrolling monocytes [blood] |
|---|---|
| BV421 | **Ly6C** |
| BV605 | **CD115** |
| FITC | **IAb** |
| PerCP EF710 | **Cx3Cr1** |
| PE | **Nr4a1***** |
| PE Cy7 | **CD11b** |
| APC [lineage] | **CD3**<br>**CD4**<br>**CD19**<br>**NK1.1**<br>**Ter119**<br>**CD8**<br>**Ly6G** |

| antibody details | | |
|---|---|---|
| | **clone** | **supplier** |
| **B220** | RA3-6B2 | ebioscience |
| **CD3** | 145-2C11 | biolegend |
| **CD4** | RM4-5 | biolegend |
| **CD8** | 53-6.7 | biolegend |
| **CD11b** | M1/70 | biolegend |
| **CD11c** | N418 | biolegend |
| **CD16/32** | 19 | ebioscience |
| **CD19** | 6D5 | biolegend |
| **CD27** | LG.7F9 | ebioscience |
| **CD34** | RAM34 | ebioscience |
| **CD43** | S11 | biolegend |
| **CD64** | X54-5/7.1 | biolegend |
| **CD71** | RI7217 | biolegend |
| **CD115** | AFS98 | biolegend |
| **CD135 = Flt3** | A2F10 | ebioscience |
| **CD169** | 3D6.112 | biolegend |
| **ckit = CD117** | 2B8 | ebioscience |
| **Cx3Cr1** | SA011F11 | biolegend |
| **F4/80** | BM8 | biolegend |
| **IAb** | AF6-120.1 | biolegend |
| **IL7Ra** | A7R34 | biolegend |
| **Ly6C** | HK1.4 | biolegend |
| **Ly6G** | 1A8-Ly6g | ebioscience |
| **NK1.1** | PK136 | biolegend |
| **Nr4a1***** | 12.14 | ebioscience |
| **Sca1** | D7 | biolegend |
| **SiglecF** | 1RNM44N | ebioscience |
| **Ter119** | Ter119 | biolegend |
| **VCAM-1** | 429 | biolegend |

Mϕ macrophages
*no unconjugated CD16/32
**auto-fluorescent
***intracellular stain

[bone marrow] **iron recycling macrophages (Mϕ)**

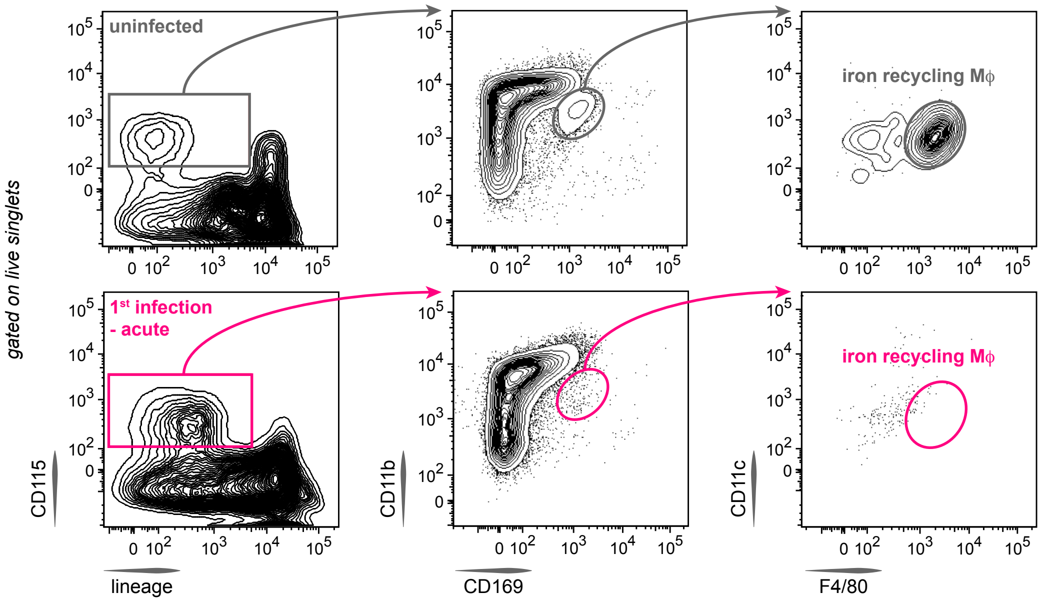


[blood] **patrolling monocytes**

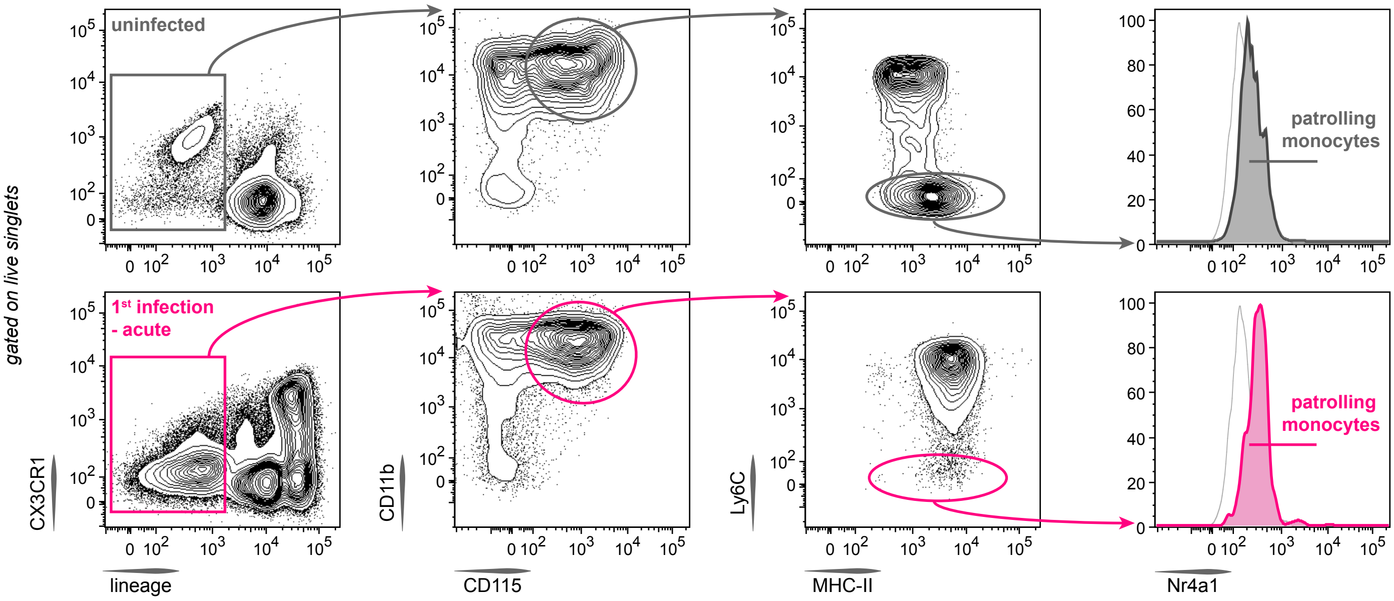


[spleen] **red pulp macrophages (Mϕ)**

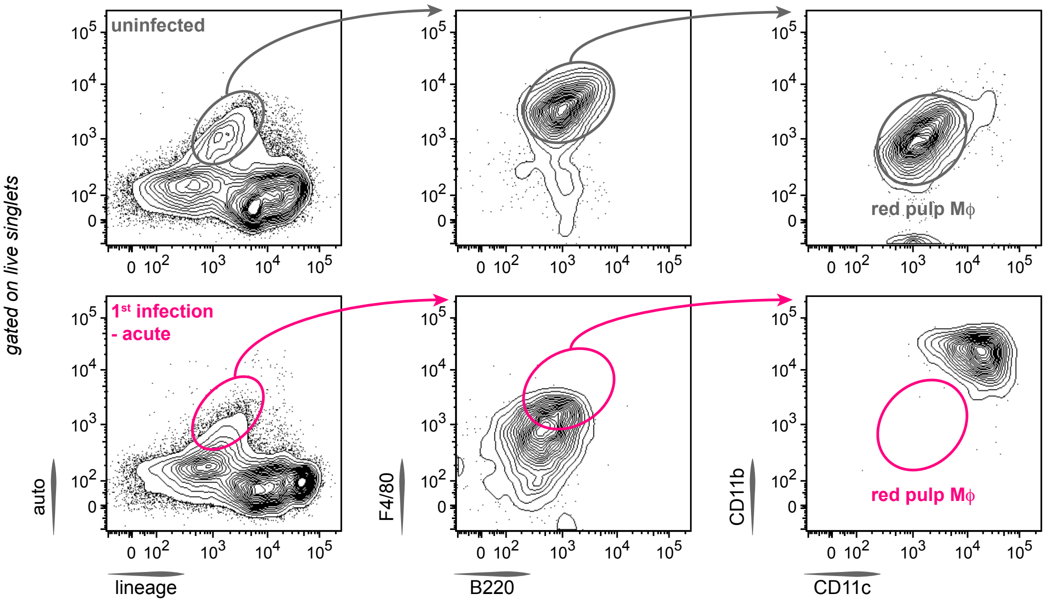

[bone marrow] **monocytes** & **neutrophils**

**uninfected**

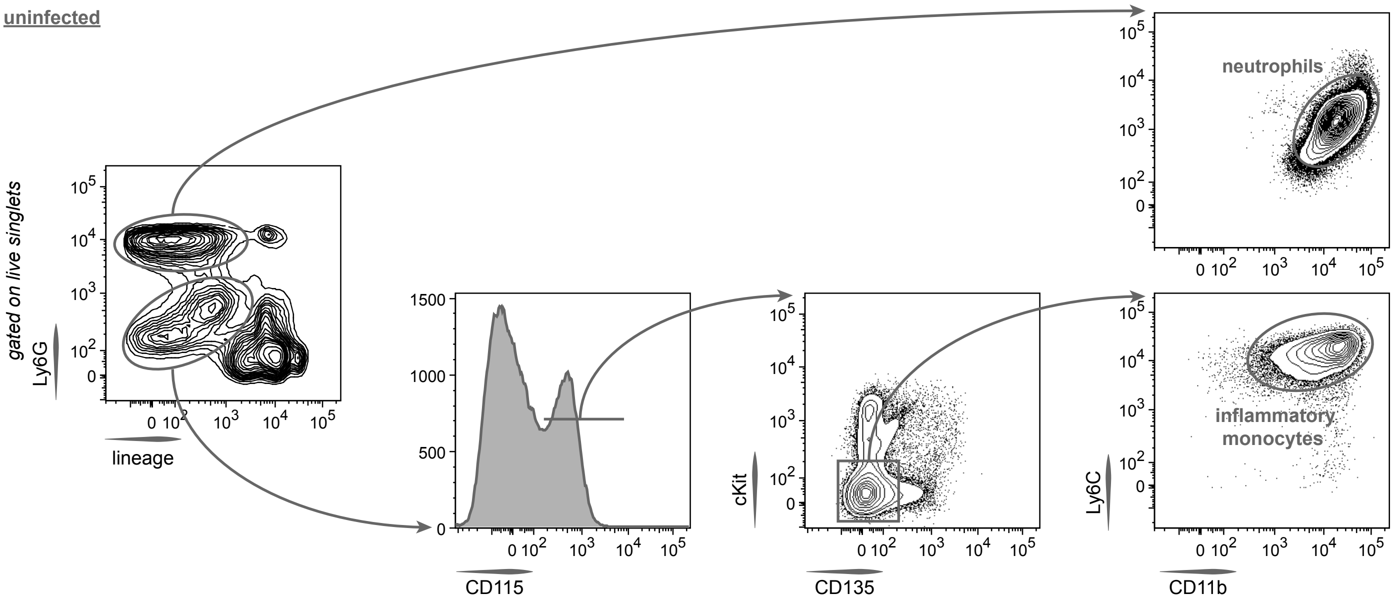


**1st infection - acute**

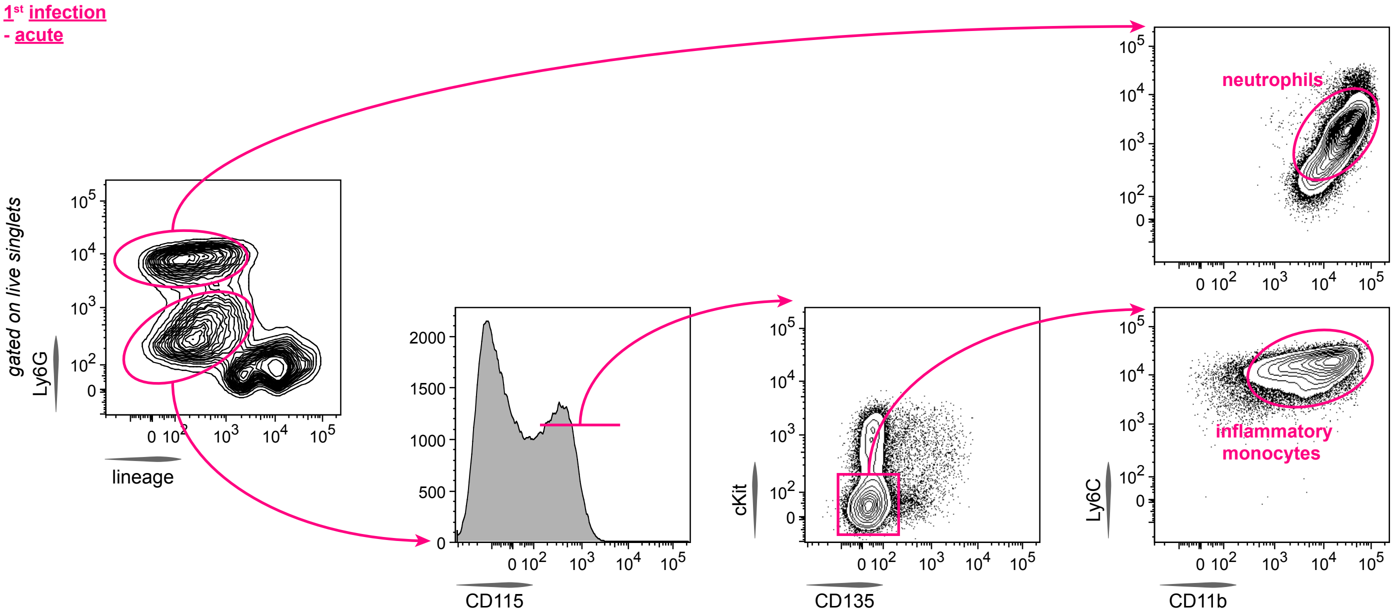

[blood & spleen] **monocytes** & **neutrophils**

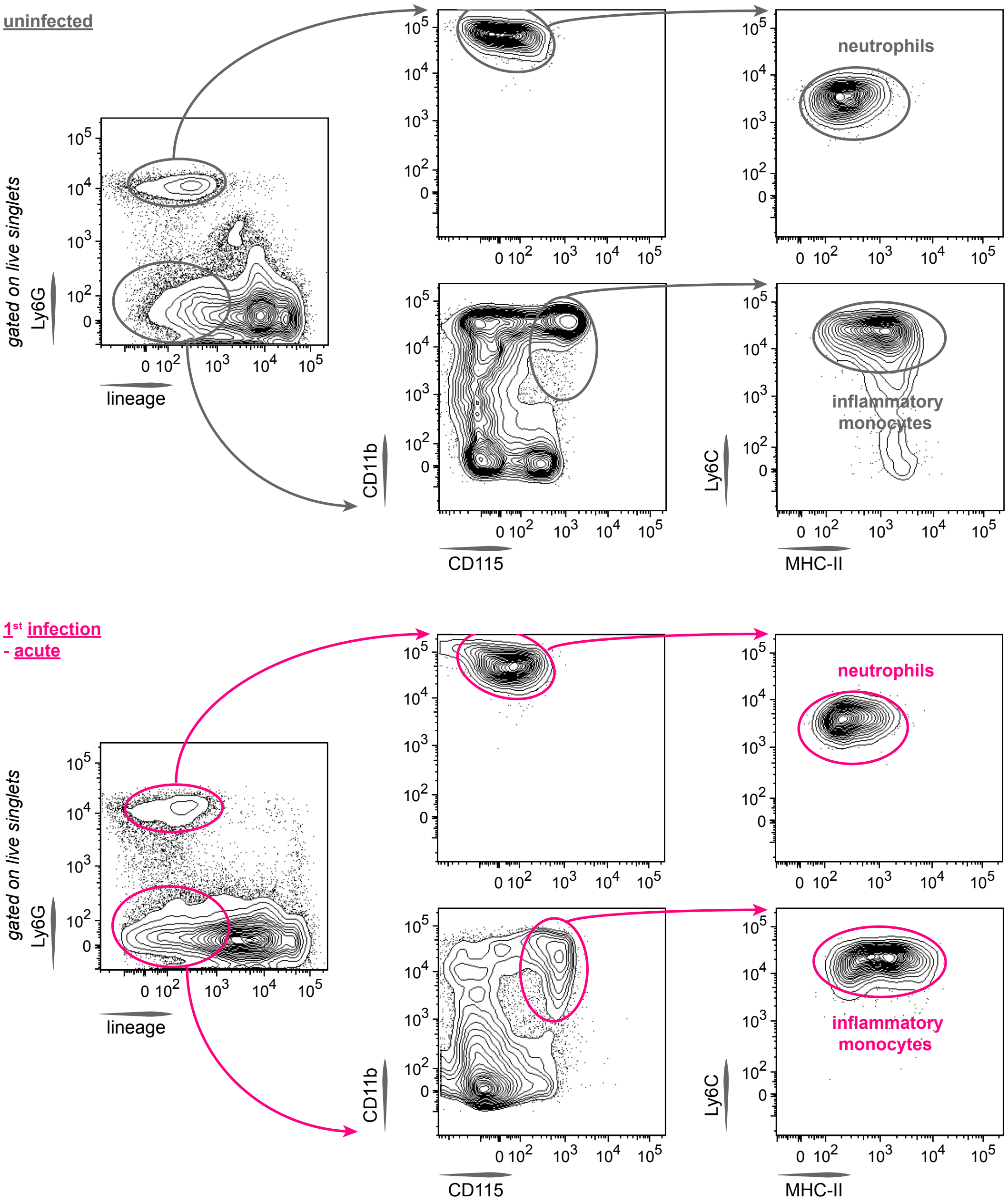

[bone marrow & spleen] **myeloid** & **erythroid progenitors**

<u>**uninfected**</u>

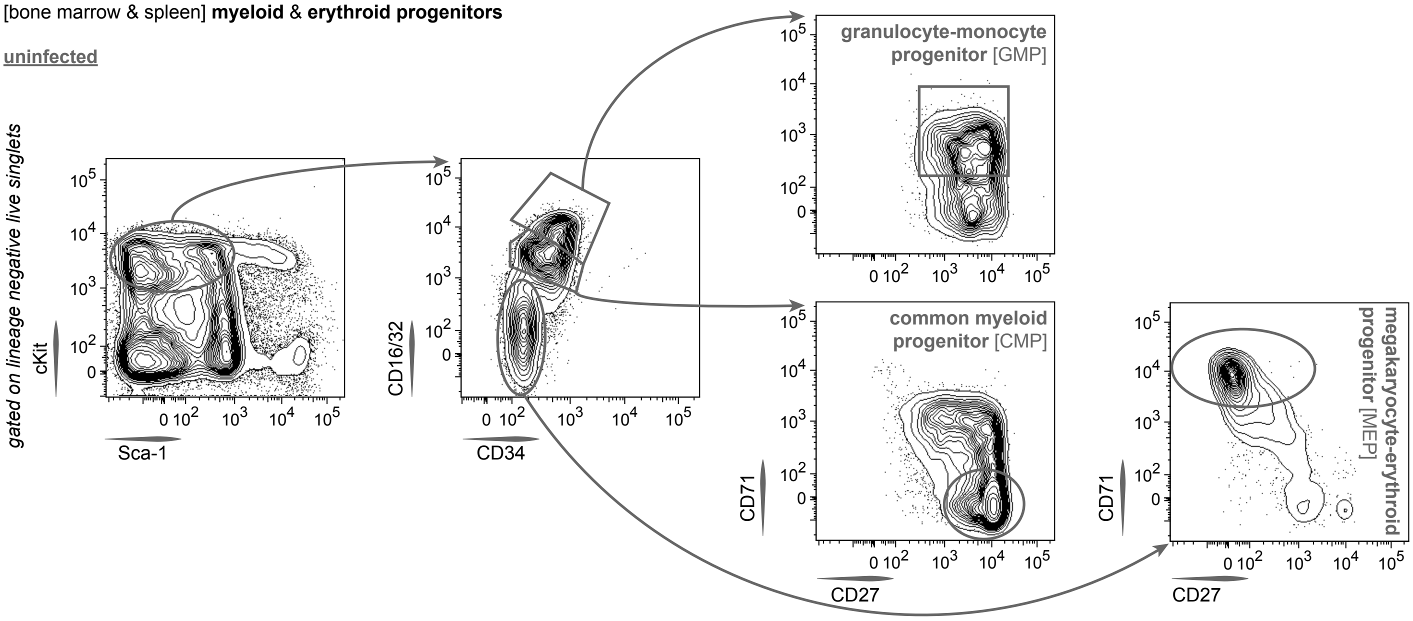


<u>**1st infection - acute**</u>

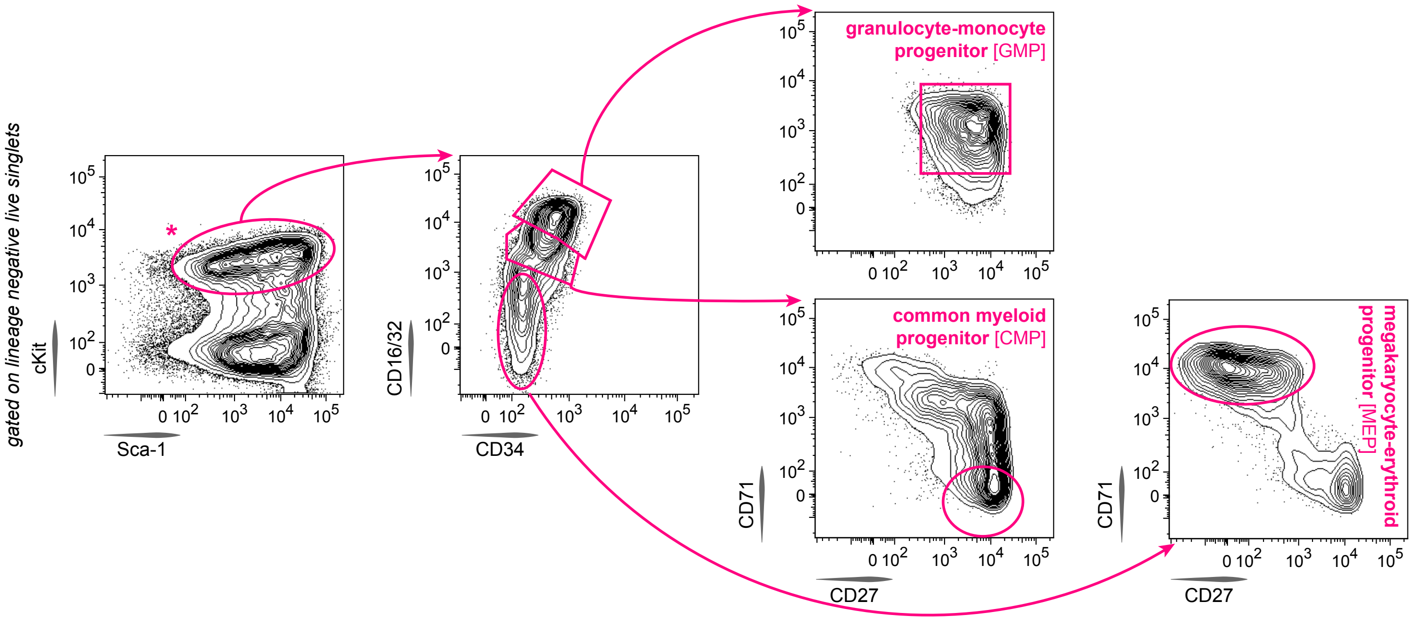
